# Lipoproteins modulate the uptake and biological function of cationic cell-penetrating peptides across the animal lineage

**DOI:** 10.64898/2026.08.23.746575

**Authors:** Sovanny R. Taylor, Temilade V. Oluwasesin, Muthu Raj Salaikumaran, Borna Novak, Adam Briner, Roshan Ailani, Pilar I. Andrade, Paulo L. Onuchic, Olivia M. S. Carmo, Garam Kim, Jacob A. Blum, Mason Galliver, Catherine Stuart, Nicolas Mendez, Vedant Patel, Guo-Teng Liang, Ronan O’Connell, Mohammad Majharul Islam, Zeinab Kashaniasl, Maria Jose Villar, Yingqiong Cao, Xi-Lei Zeng, Lucas Porta, Haoyun Yang, Gabe Hohensee, Misha Kavalur, Rachel N. Arey, Sarah E. Blutt, Ewan K. S. McRae, Radbod Darabi, Lilei Zhang, Kathryn Jones, Simon G. Pfisterer, Chad Johnston, Jeroen Pollet, Mirian A. F. Hayashi, Aaron D. Gitler, Hyun-Kyoung Lee, Alex S. Holehouse, Steven J. Ludtke, Steven Boeynaems

**Affiliations:** Department of Molecular and Human Genetics, Baylor College of Medicine, Houston, TX, USA; Jan and Dan Duncan Neurological Research Institute, Texas Children’s Hospital, Houston, TX, USA; Department of Biochemistry and Molecular Pharmacology, Baylor College of Medicine, Houston, TX, USA; Department of Biochemistry and Molecular Biophysics, Washington University School of Medicine, St. Louis, MO, USA; Center of Biochemistry Condensates (CBC), Washington University in St. Louis, St. Louis, MO, USA; Medical Scientist Training Program, Washington University School of Medicine, St. Louis, MO, USA; Clem Jones Centre for Ageing Dementia Research (CJCADR), Queensland Brain Institute (QBI), The University of Queensland, Brisbane, Australia; Departments of Biomolecular Sciences and Molecular Neuroscience, Weizmann Institute of Science, Rehovot, Israel; Department of Pediatrics, Section of Neurology, Baylor College of Medicine, Houston, TX, USA; Department of Genetics, Stanford University School of Medicine, Stanford, CA, USA; Broad Institute of MIT and Harvard, Cambridge, MA, USA; Center for Precision Environmental Health, Baylor College of Medicine, Houston, TX, USA; Department of Neuroscience, Baylor College of Medicine, Houston, TX, USA; Department of Anatomy, Faculty of Medicine, University of Helsinki, Helsinki, Finland; Institute of Muscle Biology and Cachexia (IMBC), University of Houston, Houston, TX, USA; Department of Pharmacological and Pharmaceutical Sciences, College of Pharmacy, University of Houston, Houston, TX, USA; Division of Pediatric Tropical Medicine, Baylor College of Medicine, Houston, TX, USA; Department of Molecular Virology and Microbiology, Baylor College of Medicine, Houston, TX, USA; Department of Pharmacology, Escola Paulista de Medicina (EPM), Universidade Federal de São Paulo (UNIFESP), São Paulo, Brazil; Center for RNA Therapeutics, Houston Methodist Research Institute (HMRI), Houston, TX, USA; Louisiana State University Health Sciences Center, New Orleans, LA, USA; Department of Molecular and Cellular Biology, Baylor College of Medicine, Houston, TX, USA; Department of Biochemistry in Medicine, Weill Cornell Medical College, Cornell University, New York, NY, USA; Department of Medicine, Baylor College of Medicine, Houston, TX, USA; Department of Integrative Physiology, Baylor College of Medicine, Houston, TX, USA; Chan Zuckerberg Biohub, San Francisco, CA, USA; Therapeutic Innovation Center (THINC), Baylor College of Medicine, Houston, TX, USA; Center for Alzheimer’s and Neurodegenerative Diseases, Baylor College of Medicine, Houston, TX, USA; Dan L. Duncan Comprehensive Cancer Center, Baylor College of Medicine, Houston, TX, USA

**Keywords:** cell-penetrating peptide, LDLR, LDL, lipoprotein, venom, antimicrobial peptide, C9orf72, amyotrophic lateral sclerosis, frontotemporal dementia, endocytosis

## Abstract

Cationic cell-penetrating peptides (+CPPs) are pervasive killer peptides produced by all animals and found in innate immune systems, venoms and neurodegenerative diseases. Despite their ubiquity, the mechanisms underlying their uptake remain opaque and intensely debated. Here, we interrogate +CPPs spanning 600 million years of animal evolution and identify endocytosis as a conserved/convergent uptake mechanism across cell types and organisms, at physiological concentrations. By combining multiplex imaging, genetic screening, cryo-electron tomography, and biophysical methods, we uncover that +CPPs seemingly universally enter eukaryote cells via “hitchhiking” on lipoproteins. We further show that the interaction of +CPPs with lipoproteins modulates their antimicrobial function. Combined, our findings establish a unified molecular framework describing a pan-eukaryote cell penetrance mechanism. As hyperlipidemia is a common comorbidity, our findings have direct implications for human health. Lastly, the insights gleaned from this work highlight design principles that may inform the future engineering of peptide-based therapeutics and delivery vehicles.

**eTOC BLURB:** Cationic cell-penetrating peptides are pervasive killer peptides in immunity, venoms and neurodegenerative disease, but their physiologically relevant molecular uptake mechanisms remain unresolved. Lipoprotein hitchhiking emerges as a conserved and convergent cell penetrance mechanism spanning the entire animal lineage and their protist pathogens, with direct implications to human health.

**HIGHLIGHTS:**

- Cationic cell-penetrating peptides are endocytosed across organisms and cell types.
- Endolysosomal uptake and escape constitute a functionally relevant entry route.
- Cationic peptides hitchhike on lipoprotein particles to gain cellular entry.
- Lipoproteins modulate the biological activity of cationic antimicrobial peptides.

## INTRODUCTION

Every living cell secretes peptides—whether it concerns peptides in molecular defense and attack systems or signaling peptides that talk to neighbors near and far. For these messages to be received, peptides need to interact with their target cells. While binding to a cell-surface receptor may be sufficient to relay the message (e.g., insulin, semaglutide), many require cellular uptake to reach their intracellular targets. These are cell-penetrating peptides, a diverse group of bioactive peptides capable of crossing cellular membranes from the extracellular space. Particularly prominent are the cationic cell-penetrating peptides (+CPPs), a class solely defined by their overall highly positive net charge^1–3^. These peptides are pervasive throughout biology, but perhaps most well-known for their role as killer peptides—both as antimicrobial peptides (AMPs) in innate immune systems and venom peptides from many venomous animals, and even as neurotoxic peptides found in human neurodegenerative diseases^4^. Given that such peptides can be fused to other proteins and broadly interact with nucleic acids and other anionic polymers^5^, they have even been explored as protein/gene/drug delivery vehicles^6–9^. Yet, despite their importance and ubiquity across biology, pathology and technology, the mechanisms by which +CPPs enter cells remain incompletely understood and highly debated.

The 1988 discovery that extracellular HIV-1 Tat protein (or fragments thereof) could enter cells^10,11^ launched the study of +CPPs. Since then, there has been an intense focus on identifying the precise mechanism of uptake, taking both passive and active processes into consideration (**Fig. 1A**). Early studies proposed direct membrane translocation, where +CPPs interact with and cross the plasma membrane in an energy-independent manner^11–14^. Later work demonstrated that some observations interpreted as direct translocation were influenced by fixation artifacts or experimental conditions. Specifically, concentration seems to be important. Historically, most studies in the field have been performed at relatively high peptide concentrations, well above what would be expected concentrations in some of their physiological contexts (**Table S1**, **Fig. S1A**, **Supplemental Text A**). In these instances, the peptide accumulation at the membrane can indeed promote spontaneous membrane destabilization and passive influx^15–21^, but whether this occurs at lower concentrations remains unclear. Other evidence argues for a contribution of various active endocytic processes^8,22–27^, especially at lower peptide concentrations (**Table S1, Fig. S1A**). Progress has been made in understanding the regulation of this uptake, leading to the identification of several endosomal trafficking genes^23,28–31^. Multiple studies have also demonstrated a role for the glycocalyx—a dense forest of surface-bound glycosaminoglycans^27,32,33^ and glycoRNAs^34^ that can trap +CPPs at the cell surface and facilitate their uptake. But these molecules are not classic endocytic receptors as they lack intracellular signaling domains that can initiate endocytosis by themselves. Rather, they are widely considered as co-receptors that aspecifically interact with canonical endocytic receptors, which enables the subsequent uptake of bound ligands^35–37^. Consequently, despite decades of investigation, it remains unclear which canonical receptors assist in +CPP endocytosis.

**Figure 1:**
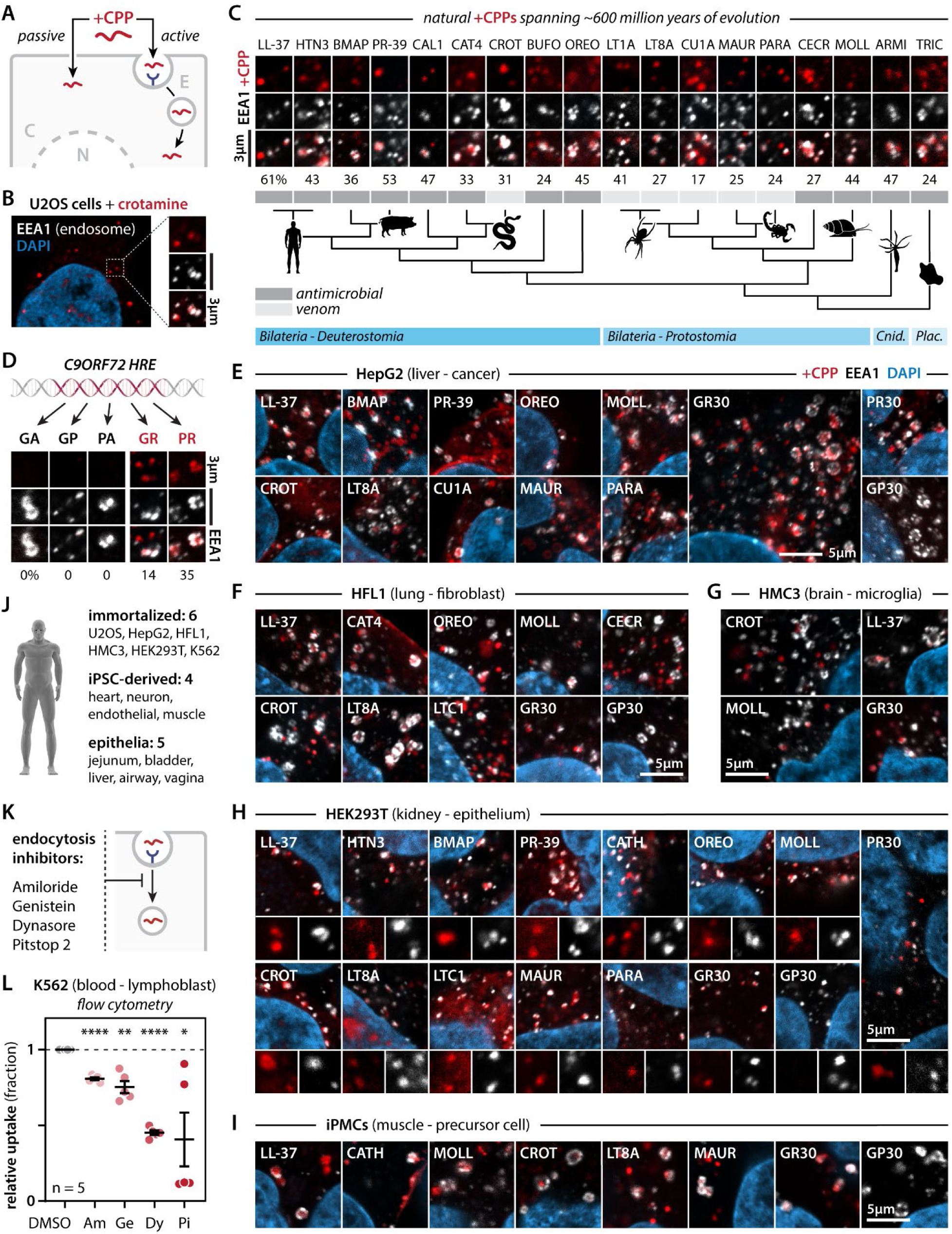
Naturally occurring +CPPs are commonly endocytosed across human cells. **(A)** Scheme illustrating passive and active +CPP uptake mechanisms. **(B)** Crotamine localizes to EEA1+ early endosomes in U2OS cells (30 min, 500 nM). Representative image. **(C)** Phylogenetically diverse natural +CPPs localize to EEA1+ early endosomes in U2OS cells. Representative zoomed images of vesicles. Numbers indicate the average percentage of peptide+ puncta that overlap with EEA1 staining. Full quantification (see also **Fig. S2A**). N = 3 replicates, n = 2,740 cells total. Abbreviations refer to *Cnidaria* and *Placozoa*. **(D)** Cationic, but not neutral, repeat peptides encoded by the C9ORF72 hexanucleotide repeat expansion (HRE) localize to EAA1+ early endosomes. Representative images. Numbers indicate average percentage of overlap (see also **Fig. S2A**). N = 3 replicates, n = 2,740 cells total. **(E–I)** Representative images of example +CPPs localizing to EEA1+ early endosomes across human cell lines, including HepG2 (**E**), HFL1 (**F**), HMC3 (**G**), HEK293T (**H**), and iPSC-derived skeletal muscle precursor cells (**I**). **(J)** List of the immortalized cell lines, iPSC-derived cell types and epithelial organoid monolayers that were found to endocytose +CPPs in this study (see also Fig. 2, **Fig. S2B-F**). **(K)** Scheme illustrating the pharmacological inhibition of endocytosis. **(L)** Flow-cytometric quantification of crotamine uptake in K562 cells following treatment with the endocytosis inhibitors Amiloride (Am), Genistein (Ge), Dynasore (Dy), and Pitstop 2 (Pi). Cellular fluorescence intensity was normalized to DMSO-treated controls. N = 5 replicates, n = 1,125,000 cells total. Repeated measures one-way ANOVA, Holm-Šídák multiple comparison test. *p < 0.05, **p < 0.01, ****p < 0.0001.

Another issue that has hindered our mechanistic understanding of +CPP uptake is that most studies have been typically limited to one or a few peptides (**Table S1**). These are often synthetic peptides (e.g., poly-arginine) or protein fragments (e.g., transcription factor fragments like Tat derived from the HIV-1 Tat protein and penetratin derived from a fly Hox gene). While such studies have provided important insights into potential cellular entry mechanisms, they may not capture the diversity of naturally occurring +CPPs. Indeed, thousands of cationic killer peptides have been described^38^. Likewise, most uptake studies have solely been performed in immortalized mammalian cell lines (**Table S1**). As a result, it remains uncertain whether the observed uptake mechanisms accurately reflect the biological pathways used by host defense peptides, venom peptides, and disease-associated +CPPs in their relevant cellular and *in vivo* contexts.

Here, we set out to investigate how natural +CPPs enter (eukaryote) cells (lacking a cell wall) in physiologically relevant conditions and whether there are conserved/convergent mechanisms at play. We did so by first compiling a phylogenetically diverse panel of natural +CPPs spanning 600 million years of animal evolution and human pathology. We demonstrate that endocytosis represents a dominant uptake mechanism in our +CPP panel and is replicated across different cell types and organisms at physiologically relevant concentrations. By combining multiplex imaging and CRISPRi genetic screening with biophysical methods and cryo-electron tomography, we have gained detailed insights into +CPP uptake that bridge the molecular and organismal scales. In doing so, we identified lipoprotein hitchhiking as a novel uptake mechanism that is shared across animal +CPPs and their eukaryote target cells. Lipoprotein uptake is one of the busiest endocytic routes for most multicellular eukaryote cells or for their single-celled parasites, making it an ideal target for +CPPs to gain cellular entry. Further demonstrating its relevance, we find that lipoproteins also modulate the physiological functions of +CPPs with direct implications to human health and drug delivery. In all, we put forward a new framework to study, understand and regulate +CPP function that informs the biology and pathology of killer peptides and highlights how we can leverage them for therapy and technology.

## RESULTS

### Assembling a phylogenetically diverse natural +CPP library

To interrogate the uptake mechanism for natural +CPPs, we assembled a library of cationic killer peptides spanning the *Eumetazoan* lineage—comprising *Placozoa*, *Cnidaria* and *Bilateria*— based on targeted literature searches for specific lineages and from a >3,300 member database^38^ covering natural AMPs. Unless when already commercially available, we opted for linear peptide sequences to avoid non-native folds of chemically synthesized products. Peptides were tagged with a C-terminal alkyne group to enable click labeling with various tags. We additionally included five peptides implicated in human neurodegenerative disease. Intronic *C9orf72* hexanucleotide repeat expansions are the major genetic cause of amyotrophic lateral sclerosis and frontotemporal dementia^39,40^. Unconventional translation of both sense and antisense RNA strands produces five different dipeptide repeats: two cationic ones, poly-(glycine-arginine) or GR and poly(proline-arginine) or PR, two neutral alanine-containing ones, namely GA and PA, and a GP peptide produced from both strands^41–43^. PR and GR have been previously described by us and others to be cell-penetrating^31,44–46^, and we have additionally shown that they mimic the biophysical mechanism of several cationic venom peptides^4^. Thus, these cationic and neutral repeat peptides will serve as “positive” and “negative” controls in this study. For the repeat peptides we chose a length of 30 repeats, as is common in the field. However, for the highly aggregation-prone amyloid-forming GA peptide we could only obtain peptides that were fifteen repeats long. Combined, all peptides make up a natural peptide library of 20 +CPPs (one folded, nineteen linear) with three neutral controls (see **Table S2** for details and abbreviations). Comparing our library to ∼1,700 annotated linear natural +CPPs indicates that they are representative of the natural variation in length and net charge (**Fig. S1B**). Thus, we are confident our library will enable us to discover general uptake mechanisms that may be broadly shared by animal +CPPs.

### Natural +CPPs are generally endocytosed across human cell types

In this study, we performed uptake assays for our peptides in the nanomolar range, to match their estimated physiological concentrations and to stay well below their acute toxic dose (**Supplemental Text A**). This approach allows us to differentiate true uptake from spontaneous influx in damaged/dead cells. We treated human U2OS osteosarcoma cancer cells, commonly used for microscopy studies given their large size, with 250 - 500 nM of fluorescently labeled Crotamine— a rattlesnake venom peptide known to be cell-penetrating^47^. 30 min after treatment, virtually all peptide accumulated in cytosolic puncta, with 31% that were positive for the early endosomal marker EEA1 (**Fig. 1B-C**, **Fig. S2A**). This argues that crotamine is endocytosed. Vesicle structures that were EEA1 negative suggest that the peptide further traffics through the endolysosomal system after its initial uptake, which we will test in more detail below. Expanding this experiment to our entire +CPP library, we surprisingly found that all partially localized to EEA1positive endosomes (17-61% of cytosolic puncta). This was the case for AMPs and venom peptides (**Fig. 1C**, **Fig. S2A**) and for the cationic neurodegenerative disease peptides (**Fig. 1D**, **Fig. S2A**). Importantly, none of the neutral control peptides were endocytosed (**Fig. 1D**, **Fig. S2A**). Note that for all peptides, except the folded crotamine, we click label peptides post-fixation and *in situ*, ruling out any effects of large bulky dye molecules on +CPP uptake. Importantly, for all experiments throughout this paper, we also include each time a no-peptide control condition to rule out non-specific click labeling (**Fig. S1C**). To assess if these results translated beyond U2OS cells, we first tested the colocalization of +CPPs with EEA1 in other immortalized human cell lines with different tissue origins (**Fig. S1D**), specifically: HepG2 (liver cancer, **Fig. 1E**), HFL1 (lung fibroblast, **Fig. 1F**), HMC3 (microglia, **Fig. 1G**), and HEK293T (kidney epithelium, **Fig. 1H**). Next, we assessed +CPPs uptake in human induced pluripotent stem cell-derived (iPSC) cell types, including precursor skeletal muscle cells (iPMCs, **Fig. 1I**), cardiomyocytes (**Fig. S2B**), endothelial cells (**Fig. S2C**), and cortical neurons (iNeurons, **Fig. 2**). Lastly, we repeated these experiments in tissue stem cell-derived organoid monolayers, including liver (**Fig. S2D**), vagina (**Fig. S2E**), bladder (**Fig. S2F**), airway epithelium (**Fig. 2**), and jejunum epithelium (**Fig. 2**). Combined, we find that +CPPs localize to endosomes in all tested human cell lines, cell types and organoids (**Fig. 1J**).

**Figure 2:**
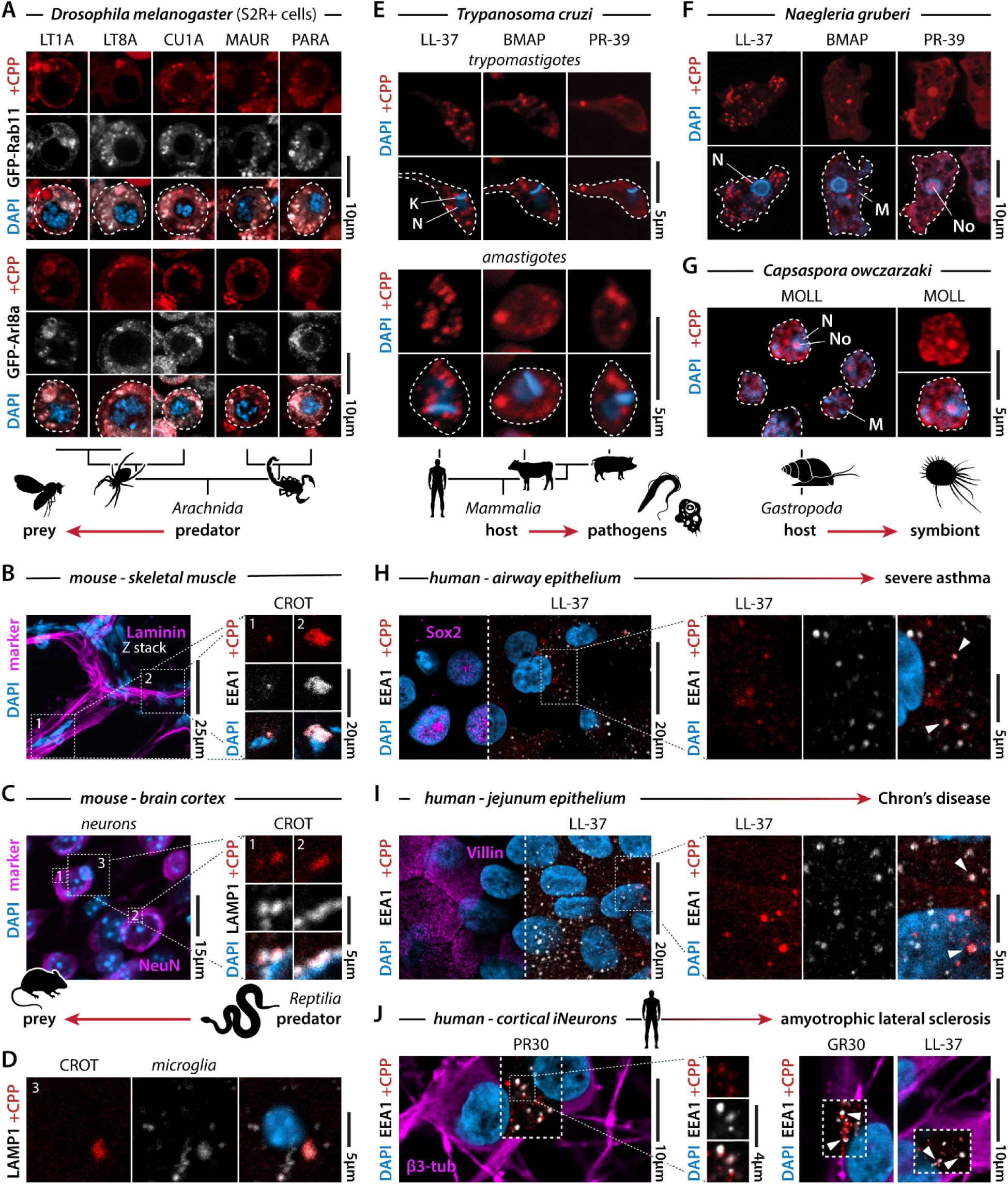
+CPPs are endocytosed in their physiological contexts. **(A)** Spider and scorpion venom peptides partially overlap with Rab11+ endosomes or Arl8a+ lysosomes in *Drosophila melanogaster* S2R+ cells (1–2 μM, 30 min). Representative images. **(B)** Cy3-labeled crotamine (6 ng) was stereotactically injected in the gastrocnemius of C57BL/6 mice and localized to EEA1+ early endosomes. Laminin marks basal lamina of muscle fibers. Representative images. Overview image is a Z-stack. **(C-D)** Cy3-labeled crotamine (6 ng) was stereotactically injected in the motor cortex of C57BL/6 mice and localized to neuronal (**C**) and microglial (**D**) LAMP1+ lysosomes. Microglia were identified based on their morphology and nuclear size. NeuN marks neurons. Representative images. Overview image is a Z-stack. **(E)** Cathelicidins localize to vesicle-like structures in *Trypanosoma cruzi* H1 trypomastigotes and amastigotes (2 μM, 60 min). K = kinetoplast. N = nucleus. Representative images. **(F)** Cathelicidins localize to food vacuole-like structures in *Naegleria gruberi* cells (1–2 μM, 60 min). N = nucleus. M = mitochondria. No = nucleolus. Representative images. **(G)** Molluscidin localizes to food vacuole-like structures in *Capsaspora owczarzaki* cells (2 μM, 30 min). N = nucleus. M = mitochondria. No = nucleolus. Representative images. **(H)** LL-37 localizes to EEA1+ early endosomes (arrowheads) in human tissue-derived airway (of nasal origin) epithelial organoid monolayers (1 μM, 60 min). SOX2 marks airway epithelium and airway progenitor cells. Representative images. Overview image is a Z-stack. **(I)** LL-37 localizes to EEA1+ early endosomes (arrowheads) in human tissue-derived jejunum epithelial organoid monolayers (1 μM, 60 min). Villin marks intestinal epithelial cells. Representative images. Overview image is a Z-stack. **(J)** PR30, GR30 and LL-37 localize to EEA1+ early endosomes (arrowheads) in human iPSC-derived cortical neurons (1 μM, 60 min). β3-tubulin marks neurons. Representative images.

To further corroborate that this localization is indeed the result of active endocytosis, we assessed the effect of established endocytosis inhibitors^48^ on the uptake of crotamine in K562 cells (125 nM), a lymphoblast suspension cell line, using flow cytometry. Amiloride, Genistein, Dynasore and Pitstop 2 all significantly reduced crotamine uptake (**Fig. 1K-L**). In all, we find strong evidence that +CPPs are commonly endocytosed across human cell types.

### +CPP endocytosis is replicated in the physiological target cells, tissues and organisms

While the above list of human cell systems was useful to demonstrate that endocytosis is a common phenomenon, obviously, most of our +CPPs did not evolve to target those specific cells. Instead, our library targets a wide range of different organisms and cell types, dependent on their specific predator-prey, host-pathogen, or disease contexts. To assess the physiological relevance of our findings, we decided to test for each +CPP class (i.e., AMP, venom, and disease) a few case studies in a setting that better mimicked their respective physiological context.

As humans universally cohabit with spiders, few predator-prey relationships are more iconic than those between spiders and flies. Latarcins^49^, Lt1a and Lt8a, are cationic venom peptides found in the venom of ant spiders from the genus *Lachesana*. Cupiennins^50^, like Cu1a, are venom peptides from the tiger bromeliad spider, *Cupiennius salei*. We cultured *Drosophila melanogaster* S2R+ cells that were engineered to express GFP-tagged Rab11 and Arl8a to mark recycling endosomes and lysosomes, respectively, both of which are downstream of early endosomes. We treated these cells with the spider venom peptides. As expected, all peptides partially colocalized with both Rab11-positive and Arl8-positive vesicles (**Fig. 2A**, **Fig. S1C**), arguing they are similarly endocytosed and likely traffic through the endolysosomal pathway—as we had observed in human cells (**Fig. 1**). The same results were obtained when we tested two scorpion venom peptides, Parabutoporin^51^ and Mauriporin^52^, from the highly venomous Schlechter’s thicktail and Moroccan black fat-tailed scorpions (**Fig. 2A**).

Moving on to vertebrate venoms, we pursued crotamine, a myotoxin found in *Crotalus* rattlesnake venoms^47^. As we cannot have snakes bite mice in the lab for obvious ethical reasons, we decided to model a snake bite using stereotactic injection. We delivered 6 ng of crotamine (a sub-physiological dose to avoid overt cell death, see **Supplemental Text A**) into the gastrocnemius muscle and brain cortex. In muscle, we observed crotamine vesicles that were positive for EEA1 (**Fig. 2B**). In brain, we found that crotamine was present in LAMP1-positive lysosomes in both neurons and microglia (**Fig. 2C-D**). These results indicate that crotamine is endocytosed in the *in vivo* context of a prey animal.

Next, we turned to AMPs. Cathelicidins^53^ are conserved across vertebrates, but highly divergent in their sequence. This family includes human LL-37, bovine Bmap-27, and porcine PR-39. Parasites from the genus *Trypanosoma* (Phylum *Euglenozoa*) infect and cause disease in humans and livestock^54–56^. For uptake studies, we isolated two of the three *Trypanosoma cruzi* life stages present in parasite-infected mammalian cell cultures: motile trypomastigotes and non-motile amastigotes. Vesicle-like structures were observed in both stages for all three peptides (**Fig. 2E**, **Fig. S1C**) and they were morphologically identical to endosomal structures described by others^57,58^. Certain members of the *Naegleria*^59,60^ genus (Phylum *Percolozoa*), popularly known as brain-eating amoebas, also infect humans and livestock. Culturing and treating (the non-infectious) *Naegleria gruberi*, we found that all three peptides localized to structures that appeared as food vacuoles^61^ (**Fig. 2F**, **Fig. S1C**), the amoeba equivalent of endolysosomes. Invertebrates also produce AMPs. We used *Capsaspora owczarzaki* (Clade *Filasterea*), a single-celled close relative of animals that colonizes mollusks^62^, as our case study. Molluscidin^63^ is a conserved mollusk AMP and readily accumulated within *Capsaspora* food vacuole-like structures^64^ (**Fig. 2G**, **Fig. S1C**). Although we are technically limited in assessing the precise nature of these vesicles in all three protist species, our observations strongly suggest that +CPPs may use similar endocytic routes as we observed above for (in)vertebrate systems. We will provide additional evidence supporting this conclusion, at least for *Naegleria*, in a later section.

While LL-37 is an AMP that has evolved to kill invading pathogens, it also moonlights as a proinflammatory signaling factor that is implicated during infection and in autoimmune disease^65^. This activity is dependent on its uptake by human cells and typically implicates endosomal innate immune receptors^66,67^, highlighting its required endocytosis to drive inflammation. LL-37 is upregulated and contributes to maladaptive inflammation in the gut of Chron’s disease patients^68^ and in the lung sputum of severe asthma patients^69^. When we grew and treated tissue stem cell-derived airway and small intestine epithelial organoids, we found that LL-37 colocalized with EEA1 in both (**Fig. 2H-I**). Thus, these findings confirm, in human organoid models, that LL-37 is endocytosed in the relevant tissues.

Lastly, we tested the uptake of the neurodegenerative disease-related GR and PR peptides in human iPSC-derived cortical neurons, which is the relevant neuronal population in frontotemporal dementia. We made identical observations on +CPP colocalization with EEA1 (**Fig. 2J**). We further extended these findings to LL-37 (**Fig. 2J**), which more recently has also been implicated in neurodegenerative^70^ and neuroinflammatory^71^ conditions.

Despite their sequence variation and disparate evolutionary origins, all examined +CPPs localized to endolysosomal(-like) compartments following cellular entry. Moreover, they did so across diverse biological contexts that are directly relevant for predator–prey and host–pathogen interactions, and human disease. While we only focused on animal peptides, our selected target organisms span over one billion years of eukaryote evolution. Collectively, these findings indicate that endocytosis is a widely shared mechanism for +CPP entry in eukaryotes.

### +CPPs traffic through the endolysosomal pathway and escape

In the above experiments (**Fig. 1-2**), we observed that peptide-laden vesicles never fully colocalized with a single marker. We interpret this as a sign of endolysosomal trafficking. Indeed, endocytosis encompasses a complex network of trafficking routes that can direct cargo toward recycling, degradation, secretion, or provide access to intracellular compartments upon escape. We therefore sought to determine whether cationic peptides indeed navigate these intracellular trafficking routes following uptake. To systematically map peptide trafficking after internalization, we employed iterative indirect immunofluorescence imaging (4i), a multiplex imaging approach that enables visualization of peptide localization relative to dozens of intracellular markers within the same cells^72–76^ (**Fig. 3A**). Using this strategy, we profiled peptide localization across 26 markers spanning the entire endomembrane system alongside other subcellular compartments (**Fig. 3B**). This allowed us to generate a comprehensive atlas of intracellular peptide trafficking while simultaneously assessing alternative subcellular destinations (**Fig. 3C**). We selected three representative peptides—crotamine, LL-37 and GR30—and treated U2OS cells with 250 nM of peptide followed by fixation and 4i analysis. We also included GP30 as a negative control. After imaging, we created composite maps of all subcellular markers, which we could now use as an atlas to identify the +CPP-positive vesicles (see **Material & Methods**). In brief, we segmented individual +CPP-positive vesicles for each cell across all experimental conditions and calculated the enrichment of each of the 26 markers in the vesicle relative to the immediate surrounding cytoplasm. After stringent quality control (see **Material & Methods**), we retained a total of 30,586 cytoplasmic vesicles from 2,630 cells.

**Figure 3:**
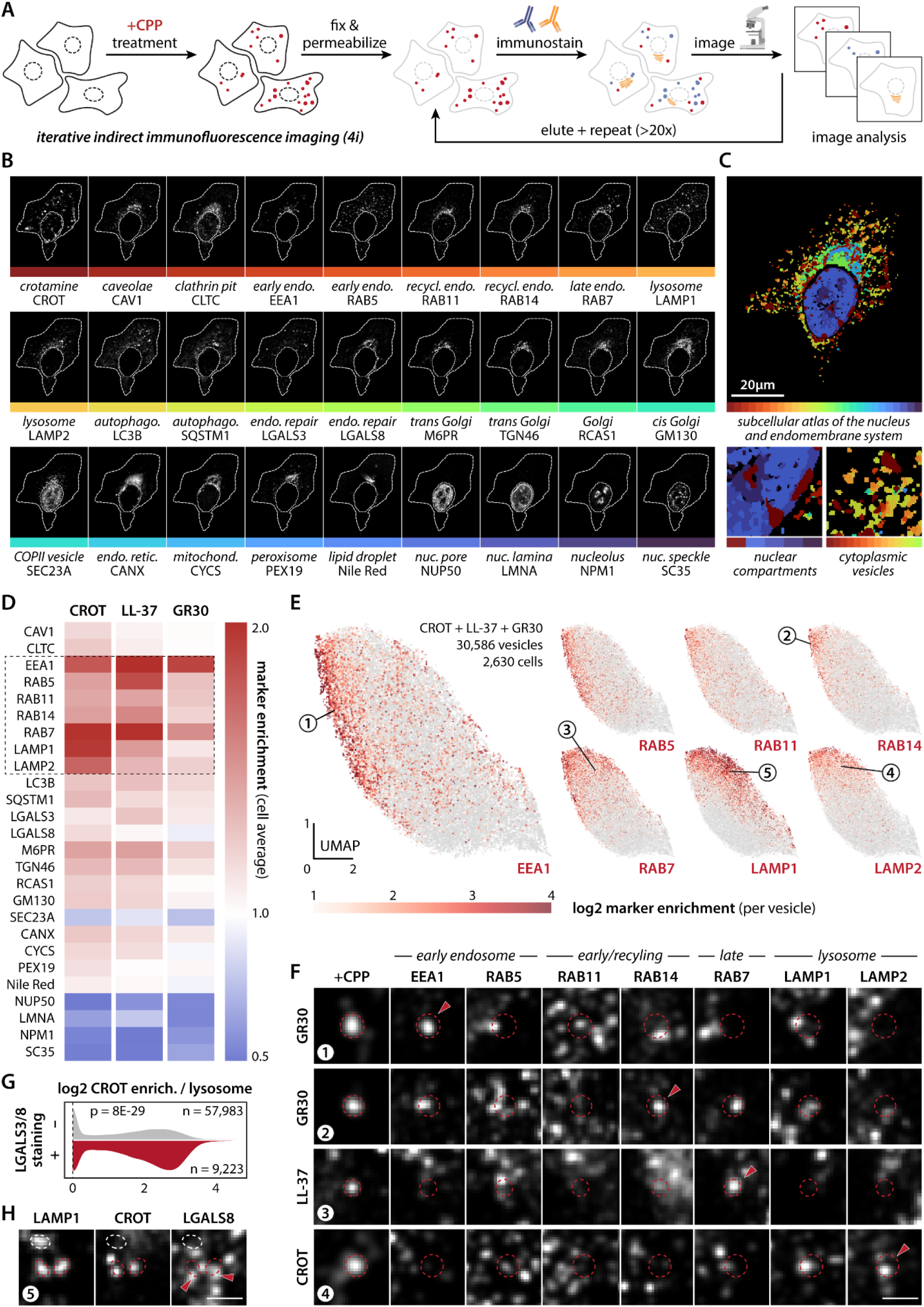
Multiplex immunostaining maps the endolysosomal trafficking and escape of +CPPs. (A) Scheme illustrating the iterative indirect immunofluorescence imaging (4i) workflow. **(B)** Representative confocal images of the same crotamine-treated cell across successive 4i rounds. Crotamine (CROT) and 26 markers of the endomembrane system and other subcellular compartments are shown. Dashed white lines indicate cell and nuclear boundaries. **(C)** Representative subcellular atlas generated from 26 segmented marker images (red to purple) overlayed with a segmented crotamine mask (dark red). Zoom highlights the spatial organization of nuclear compartments and cytoplasmic vesicular structures within an individual cell. **(D)** Heatmap showing the enrichment of 26 markers within peptide+ puncta (cytoplasmic and nuclear) for cells treated with crotamine, LL-37 or GR30. Vesicle enrichment values were averaged per cell. Values around one 1 indicate no enrichment, while higher and lower values indicate strong average enrichment or depletion, respectively. N = 9 replicates, N = 2,789 cells total, and n = 37,968 peptide+ puncta total. Dashed box highlights strong enrichment of EEA1, RAB7, LAMP1 and other endolysosomal markers across peptide conditions. **(E)** UMAP cluster of 30,586 peptide+ cytoplasmic vesicles from 2,630 analyzed cells based on enrichment values for seven endolysosomal markers. Overlapping and contrasting enrichment gradients can be observed across the vesicle population. **(F)** Representative single-vesicle images corresponding to the numbered locations in the UMAP cluster (**E**). Images show peptide signal (+CPP) and the indicated endolysosomal markers. Red dashed circles indicate the outline of the peptide+ vesicle. Arrowheads highlight the major enriched marker. Scale bar = 3 μm. **(G)** LAMP1+ lysosomes with detectable LGALS3/8 enrichment (red; n = 9,223 lysosomes) have on average a higher crotamine intensity than those without LGALS3/8 enrichment (gray; n = 57,983 lysosomes), indicative of lysosomal membrane damage and peptide escape (see also **Fig. S3A-B**). Two-sided Kolmogorov– Smirnov test. **(H)** Representative image of crotamine+ (red dashed line) and crotamine(white dashed line) lysosomes. Crotamine+ lysosomes show a stronger association with LGALS8 staining. Scale bar = 3 μm.

To gain a high-level understanding of the major subcellular compartments that were sampled by the trafficking peptides, we averaged the enrichment scores of all vesicles+puncta per cell for each experimental condition. As expected, we saw a strong per cell enrichment for the early endosomal marker EAA1 across all peptides (**Fig. 3D**). The same was true for the late endosomal marker RAB7. Encouragingly, we had previously identified *RAB7A* knockdown in a CRISPR screen to suppress the toxicity of the PR peptide in mouse primary neurons^31^. Thus, RAB7 function seems to modulate the toxicity of +CPPs by regulating their uptake and/or trafficking. Other markers that were most enriched were the early endosomal RAB5, the recycling endosomal RAB11 and RAB14, and the lysosomal LAMP1/2. While all peptides strongly colocalized with EEA1, we did observe some subtle differences between them. For example, GR30 showed a lower enrichment of late endolysosomal markers, in line with its slower uptake by cells (**Fig. 3D**). Additionally, crotamine and LL-37 showed differential enrichment for lysosomal markers. These findings suggest that these different peptides may have slightly different trafficking routes (or degradation rates) post endocytosis. Lastly, while nuclear markers were depleted on the whole cell level and serve as negative controls (**Fig. 3D**), at the imaged time points we already observe cells with localization of the peptides to subnuclear compartments (**Fig. 3C**), as we reported before^4^.

For a more granular analysis of our trafficking data, we compiled all cytoplasmic vesicles and clustered them using dimensionality reduction (Uniform Manifold Approximation and Projection or UMAP, see **Material & Methods**). In contrast to defined clusters, we found that all vesicles grouped together with specific endolysosomal markers forming overlapping or opposing gradients across the population (**Fig. 3E**). This result is in line with the dynamic nature of endolysosomes, which instead of populating discrete states sample a continuous distribution along their trafficking route, where individual vesicles gradually change “identity” by exchanging effector proteins^77–79^. For example, the early endosomal EEA1 and lysosomal LAMP1 most highly enriched in opposing sides of the population, with gradients of other markers (e.g., late endosomal RAB7) bridging them—following the expected endosome-to-lysosome transition. Note that while RAB14 is commonly known as a recycling endosomal marker, it has recently been implicated in an alternative endosomal route that is used by other +CPPs^28^. We find that RAB14 often co-enriches with +CPP-positive early/late endosomes at the population level (**Fig. 3E**). This shared involvement is also observed when investigating individual vesicles (i.e., vesicle #2 in **Fig. 3F**). Thus, as hypothesized, we find strong data that +CPPs traffic through the entire endolysosomal cascade after their cellular uptake.

While all the above data overwhelmingly support that +CPPs are endocytosed, does this matter for their function? Since we observe these peptides in lysosomes, one could argue that the endocytic route presents a functional dead end, where peptides are trapped and degraded before they can kill the cell. We and others have shown that several +CPPs kill cells by entering the nucleus and perturbing RNA metabolism^4,44^. Also here, we find that +CPPs localize to RNA-rich nuclear bodies like nucleoli and nuclear speckles (**Fig. 3B-C**). Thus, to reach their target site, +CPPs must undergo endolysosomal escape—a process where the lysosomal membrane ruptures and releases its contents^80,81^. Damaged lysosomes expose luminal glycans to the cytosol, which are bound by galectins that regulate repair or clearance and are used as indicators of endosomal escape^82–84^. However, galectins are also implicated in a number of other processes, such as cell cycle, cell growth, and inflammatory responses which continually occur in cells and are unrelated to escape^85,86^. To assess galectin recruitment associated with endolysosomal damage, rather than galectin puncta arising from other biological processes, we segmented all lysosomes in our multiplexed dataset (n = 67,206) and quantified the enrichment of galectin-3 and galectin-8 for each of them. Lysosomes enriched for either galectin exhibited a higher peptide intensity across all peptides examined (**Fig. 3G-H**; **Fig. S3A**). These findings suggest that lysosomes with higher peptide loads are more likely to recruit galectins, consistent with increased endolysosomal membrane damage and activation of the membrane repair response. We obtained identical results for RAB7+ late endosomes (**Fig. S3B**). In all, we conclude that endocytosis is a common *and* functionally relevant entry route for killer peptides.

### Lipoprotein receptors mediate +CPP entry

To our surprise, we found that unrelated natural +CPPs from across the animal lineage all have convergently evolved the capability to undergo endocytosis, and that they do so in their physiological context. Yet, an important question remains: which are the implicated endocytic receptors?

We sought to identify +CPP uptake modulators using a genome-wide CRISPR interference (CRISPRi) screen (**Fig. 4A**). To this end, we modified a screening approach that we previously used to assess modulators of intracellular protein levels^87^. In brief, we introduced a CRISPRi guide library in K562 cells and treated them with a low concentration (125 nM) of crotamine, which we used as our case study. We also confirmed that crotamine uptake was dose-dependent and therefore fluorescence intensity could be correlated to uptake levels (**Fig. 4B**; **Fig. S4A**). Next, we sorted cells by fluorescence-activated cell sorting into three different populations based on their fluorescence intensity (i.e., top and bottom 10% + middle 80%; **Fig. 4A**) followed by guide RNA sequencing and casTLE enrichment analysis^88^. We identified a total of 571 genes whose knock-down significantly reduced or increased crotamine uptake, covering a variety of biological pathways (**Table S3-4**, **Fig. S4B-D**, **Supplemental Text B**). Encouragingly, some of our hits overlapped with those from other +CPP uptake^21^ or toxicity^31^ screens, indicating that we are discovering relevant modifiers of +CPP biology (**Fig. 4E**).

**Figure 4:**
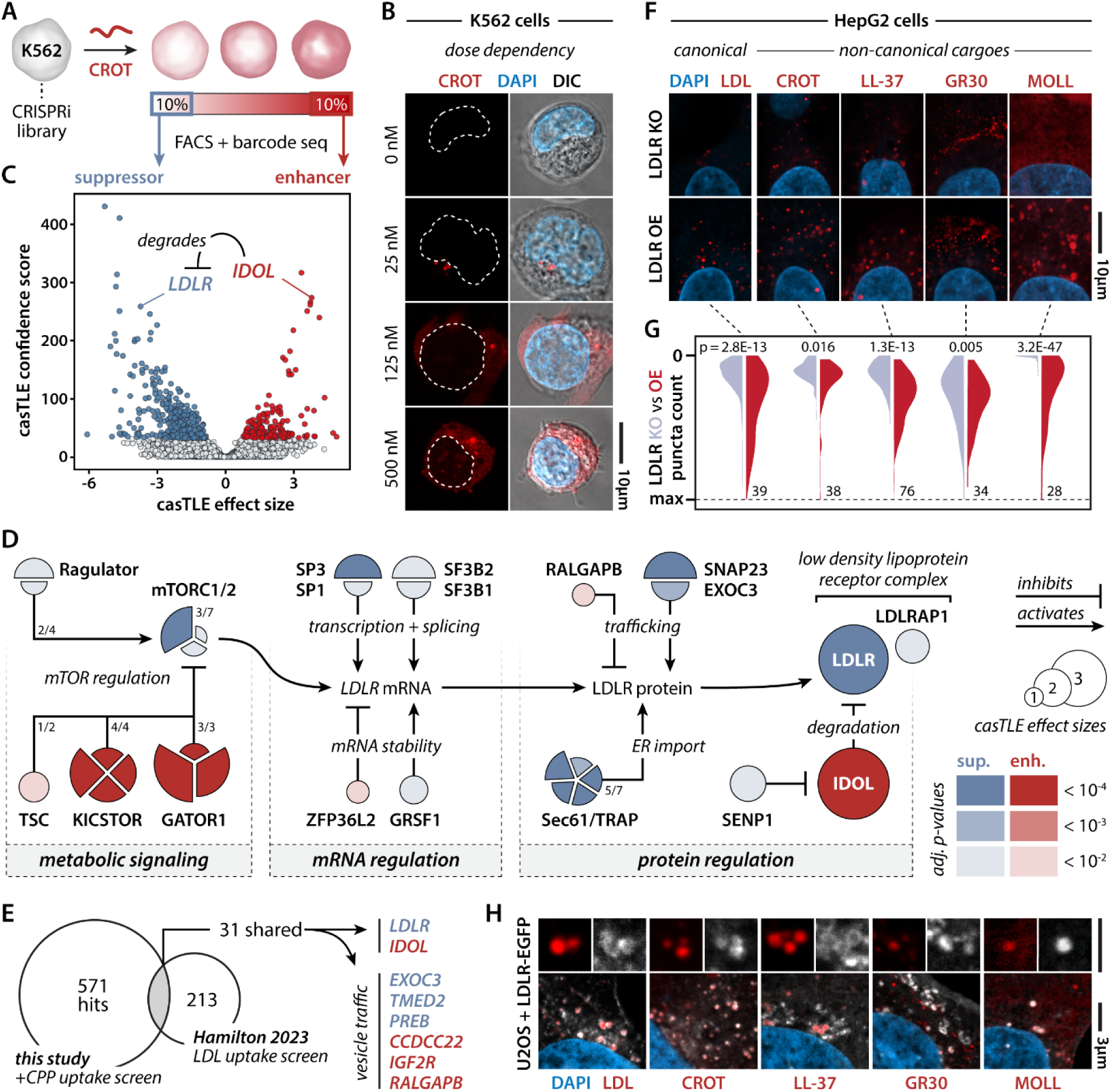
A pooled CRISPRi screen identifies LDLR as a dose-dependent modulator of +CPP uptake. **(A)** Scheme of the CRISPRi screen setup. A K562 CRISPRi library was treated with fluorescently labeled crotamine (125 nM, 60 min), sorted for top and bottom 10% cells based on fluorescence intensity by FACS, and subjected to barcode (i.e., sgDNA) sequencing. **(B)** Representative confocal images showing dose-dependent crotamine uptake in K562 cells (0–500 nM; see also **Fig. S4A**). Dashed lines outline the nucleus. **(C)** Volcano plot highlighting the significant enhancers (red) and suppressors (blue) of crotamine uptake. *LDLR* and its negative post-translational regulator *IDOL* are highlighted. **(D)** Screen hits are enriched for genes involved in metabolic signaling, and *LDLR* gene and LDLR protein expression and regulation. Color intensity corresponds to the adjusted p-value. Fractions indicate the number of complex subunits identified as hits in the screen. **(E)** Crotamine-uptake screen (CRISPRi, K562) hits overlap with LDL-uptake screen hits (CRISPR, HepG2). **(F)** Representative images of LDLR-knockout (LDLR KO) and LDLR-overexpressing (LDLR OE) HepG2 cells treated with LDL or the indicated +CPPs. **(G)** LDLR dose-dependently promotes LDL and +CPP endocytosis. Quantification of per-cell cargo+ puncta counts between LDLR-KO cells (blue) and LDLR-OE cells (red). Two-sided Kolmogorov–Smirnov test (see also (**Fig. S5A-F**). Violin plots were scaled relatively to the maximal puncta/cell count identified in each experimental comparison. Numbers indicate maximal puncta/cell count. n = 90-226 cells per condition. **(H)** LDL, crotamine, LL-37, GR30, and Molluscidin localize to LDLR-EGFP+ vesicles in U2OS cells. Representative images.

We identified only one canonical endocytic receptor whose knock-down reduced crotamine uptake: the *low-density lipoprotein receptor* (*LDLR*), a cell-surface receptor that mediates lipoprotein uptake (**Fig. 4C**). Adding to its significance, knockdown of *inducible degrader of the LDL receptor* (*IDOL*, also known as *MYLIP*) increased uptake, indicating that LDLR is a dose-dependent mediator of crotamine cell entry. Given its importance to cardiovascular and metabolic disease, there is a rich literature on LDLR regulation. Investigating our hit lists, we found several genes that had previously been directly implicated in all aspects of LDLR’s life cycle and the upstream nutrient sensing pathways (**Fig. 4D**, **Supplemental Text C**). This convergence on LDLR was further exemplified by several hits being shared with those identified in an LDL uptake screen in HepG2 cells^89^ (**Fig. 4E**).

To validate LDLR’s role in +CPP uptake, we used HepG2 cell lines where LDLR was knocked out or overexpressed, respectively. Given its liver origin, an organ with a central role in regulating lipoprotein metabolism, this cell line has been commonly used by the field to study LDL uptake. In line with our screen results, the uptake of crotamine, LL-37, GR30 and molluscidin was modulated by LDLR levels and followed the same trend as the uptake of LDL itself (**Fig. 4F-G**, **Fig. S5A-F**).

Our findings suggest that LDLR plays a direct role in the uptake of +CPPs. If this were the case, one would expect that LDLR would colocalize with these peptides in endolysosomal vesicles. When we expressed EGFP-tagged LDLR in U2OS cells and treated them with our four example +CPPs, we indeed found that they colocalized to vesicles (**Fig. 4H**). This colocalization was indistinguishable from the colocalization we observed for LDL and LDLR, arguing that this most likely presents the canonical endocytic route of LDL(R).

### +CPPs are co-endocytosed with lipoproteins

If +CPPs indeed use LDLR for cellular entry, we need to consider its canonical ligands. Lipoprotein receptors bind to and endocytose lipoproteins, which are abundant lipid transport particles that circulate in biofluids and the extracellular space. We envision two potential mechanisms of how LDLR could mediate +CPP entry. First, +CPPs and lipoproteins may compete for the same receptor, where uptake of one prevents the uptake of the other. Second, +CPPs may directly interact with the lipoproteins, which will enable their joint uptake. We call these two processes “lipoprotein receptor hijacking” and “lipoprotein hitchhiking”, respectively (**Fig. 5A**). Of note, the latter process would provide an explanation as to how endocytosis is actually initiated, as the small peptides are unlikely to interact with the receptor in a manner similar to the much larger lipoproteins (**Fig. 5B**).

**Figure 5:**
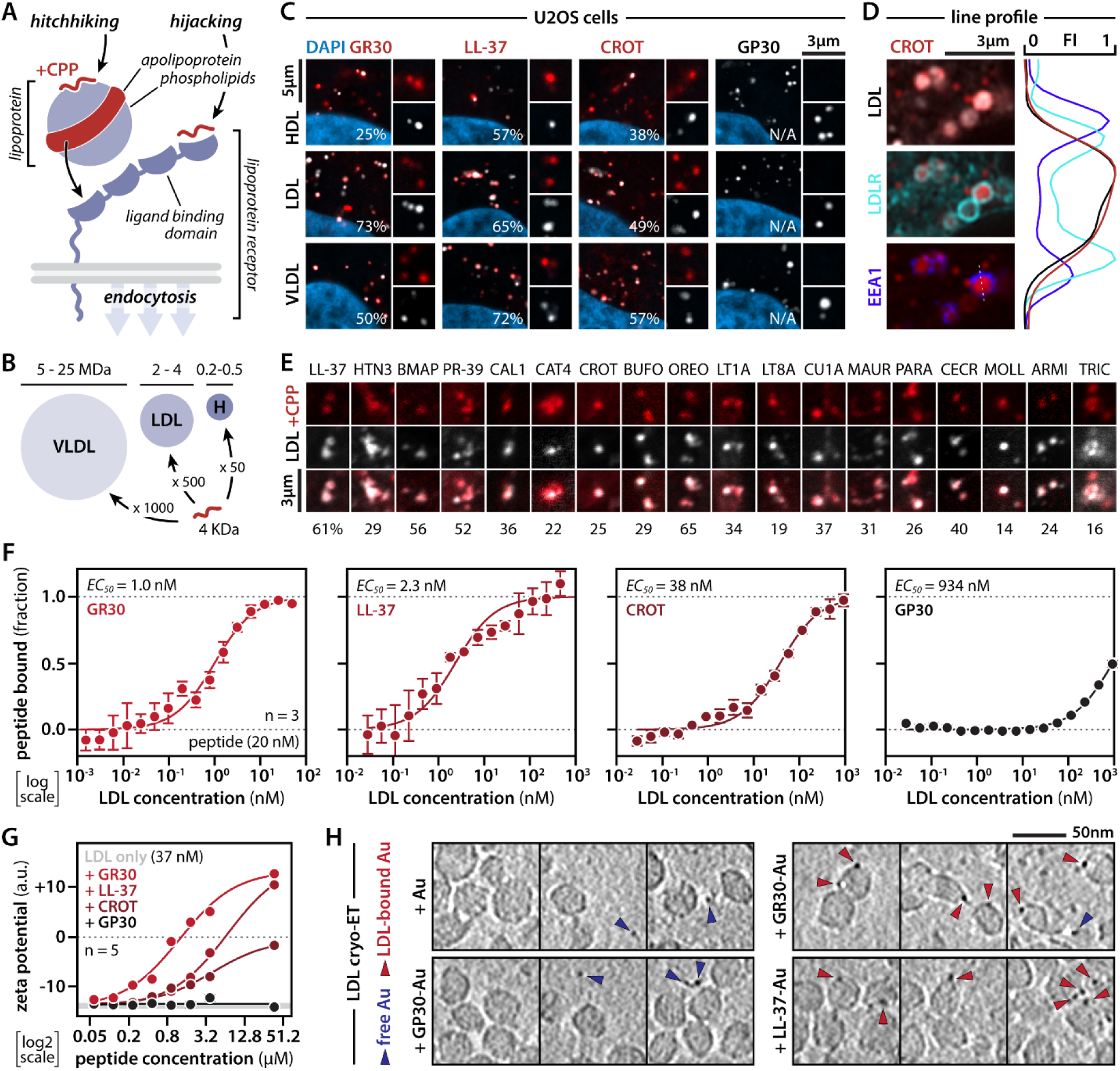
+CPPs directly bind to and co-endocytose with lipoproteins. **(A)** Scheme illustrating two non-mutually exclusive models of lipoprotein receptor-mediated +CPP uptake, namely lipoprotein hitchhiking and lipoprotein receptor hijacking. **(B)** Scheme comparing the approximate molecular masses of lipoprotein species to a 36 amino acid peptide. Approximate mass differences relative to the peptide are indicated. **(C)** GR30, LL-37 and crotamine, but not GP30, colocalize with lipoproteins to vesicles in U2OS cells (500 nM, 60 min). Representative images. Percentages indicate the average overlap between peptide+ puncta and lipoproteins (see also **Fig. S6A**). N = 3 replicates; n = 1,818 cells total. N/A indicates that overlap was not detected. **(D)** Crotamine localizes with LDL in LDLR+/EEA1+ early endosomes. Representative image with line plot. Fluorescence intensities were normalized from 0 to 1 along the indicated dashed line. (see also **Fig. S6C**) **(E)** All tested natural +CPPs colocalize with LDL into vesicles in U2OS cells. N = 3; n = 2,958 cells total. Percentages indicate the average overlap between peptide+ puncta and lipoproteins (see also **Fig. S6B,D**). Representative images. **(F)** LDL-binding curves for GR30, LL-37, crotamine and GP30 generated using spectral-shift and microscale thermophoresis measurements. The fraction of peptide bound was measured across increasing LDL concentrations at a fixed peptide concentration of 20 nM. Apparent EC_50_ values are indicated. N = 34 replicates, error bars = SEM. (See **Table S8**) **(G)** Zeta potential (i.e., surface charge) of LDL alone (grey box indicates mean +SEM) or in the presence of increasing concentrations of GR30, LL-37, crotamine or GP30. LDL was maintained at ∼37 nM (100 µg/ml). N = 5, error bars = SEM. **(H)** Cryo-electron tomography indicates that nanogold-labeled GR30 and LL-37, but not GP30 or nanogold itself, bind to the surface of LDL particles. Representative images (see also **Fig. S6F**, **Supplemental Video 1-4**). Red and blue arrowheads indicate LDL-associated and free nanogold particles, respectively.

To gain insight into which of these mechanisms would be the predominant one, we treated cells with both +CPPs and fluorescently labeled lipoproteins (i.e., very low density VLDL, low density LDL, and high density HDL) that were purified from human serum (**Fig. 5B-C**, **Fig. S6A**). We observed robust colocalization between our entire +CPP library and lipoproteins (**Fig. 5D-E**, **Fig. S6B-D**). Moreover, we found that +CPPs colocalized with LDL and LDLR in early endosomes (**Fig 5D**, **Fig. S6C**). This finding is in line with a hitchhiking mechanism rather than a competition model. It also explains why all lipoproteins, not only the canonical LDLR cargo LDL, co-endocytose with +CPPs.

### +CPPs directly bind to lipoproteins

Our functional data propose that +CPPs may bind to lipoproteins. The surface of these particles consists of a monolayer of anionic phospholipids with a “belt”^90^ formed by apolipoproteins that have cationic binding sites for lipoprotein receptors (**Fig. 5A**). Microscale thermophoresis/ spectral shift (**Fig. 5F**) and gel shift assays (**Fig. S6E**) demonstrated the direct interaction of +CPPs with lipoproteins. If +CPPs bind to the anionic lipoprotein surface, we reasoned that the overall surface charge of the lipoprotein particles would change. Zeta potential measurements indeed revealed that +CPPs dose-dependently first neutralized and later even inverted lipoprotein surface charge for our most charged peptides (**Fig. 5G**).

As our biophysical and biochemical methods provided robust evidence for +CPP-lipoprotein interactions, we wondered if we could directly visualize them. Over a decade ago, we reported the structure of the LDL-LDLR complex using cryo-electron microscopy^91^. Yet, despite the increased resolution of current microscopes, our peptides are typically too small to observe. To circumvent this, we took advantage of clickable gold nanoparticles to tag our +CPPs. When we incubated LDL particles with nanogold-labeled GR30 and LL-37, we confirmed that these nanogold particles were bound to the LDL surface (**Fig. 5H**, **Fig. S6F, Supplemental Video 1-2**). Importantly, this was not the case for nanogold particles by themselves or those that were clicked to GP30 (**Fig. 5H**, **Fig. S6F, Supplemental Video 3-4**). Thus, we conclude that +CPPs can directly bind to lipoproteins, which leads to their joint uptake.

Our *in vitro* experiments highlight that +CPPs can bind lipoproteins, but do they actually do so *in vivo*? Given their importance to metabolic and cardiovascular disease^92,93^, many studies have isolated human serum lipoproteins for proteomics analysis. Across datasets, LL-37 is commonly identified as natively associated with HDL^94^, LDL^95,96^, VLDL^95,97^ and chylomicrons^97^. This association was further supported by the increased interaction of LL-37 and LDL in the arteries and plasma of atherosclerosis patients, which even served as a biomarker for disease severity^98^. Besides LL-37, we identified three other cationic AMPs (or their pro-proteins) in these datasets (i.e., CXCL4^94,96,97^, CXCL7^96,97^, DEFA1^94^). These findings support that +CPPs are readily bound by lipoproteins in the physiological *in vivo* context.

### Lipoproteins modulate the biological function of +CPPs

Having established that lipoproteins interact with and modulate the uptake of +CPPs, we wondered if the same was true for their bioavailability and killer activity. Given its established interaction with lipoproteins in humans^94–97^, we focused on LL-37. This AMP potently kills *E. coli* cells. However, co-treating with increasing doses of lipoproteins completely blocked its bactericidal activity (**Fig. 6A**). In other words, lipoproteins act as a sink, sequestering this AMP away from the bacterial plasma membrane (**Fig. 6B**). Note that we saw complete inhibition of LL-37 killing at LDL levels that were at least two orders of magnitude below the lower bound physiological plasma concentration (i.e., ∼50 mg/dl). Others had previously observed that addition of serum can inhibit AMPs in antibacterial assays^99^, yet why remained unclear. Our finding suggests that lipoproteins are at least in part responsible for this effect.

**Figure 6:**
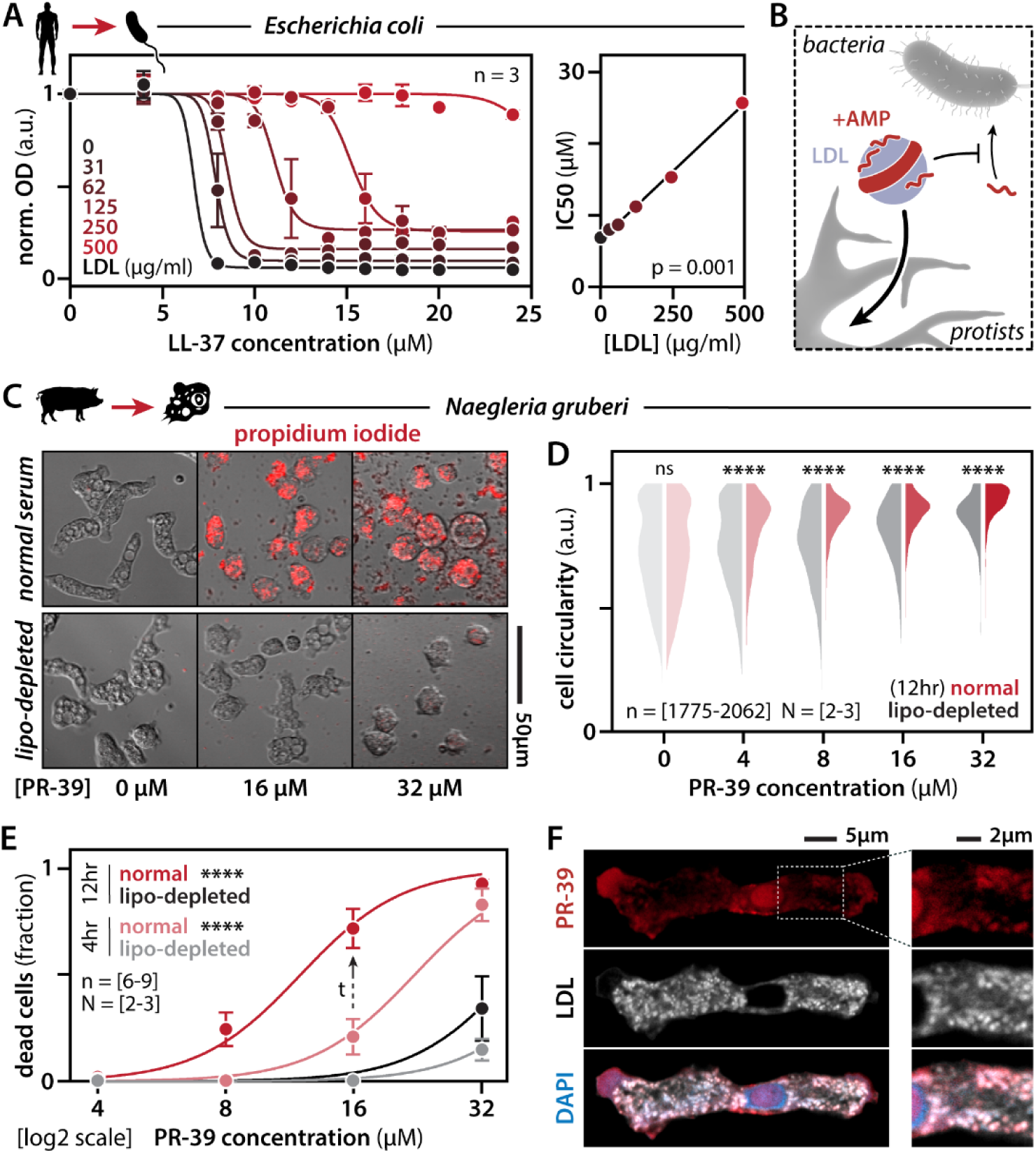
Lipoproteins differentially modulate the antimicrobial activity of cathelicidins in bacteria and protists. **(A)** LDL addition dose-dependently counteracts the antimicrobial activity of LL-37 in *Escherichia coli*. (Left) Bacterial growth is shown as normalized optical density (OD). N = 3, error bars = SEM. Curve fits are variable slope (four parameter) dose response curves. (Right) LL-37 half-maximal inhibitory concentrations (IC_50_) linearly correlates with the LDL concentration. **(B)** Proposed model in which lipoproteins sequester cationic antimicrobial peptides (+AMPs), limiting their activity against bacteria while facilitating their uptake by and activity in protists. **(C-E)** PR-39 treatment induces cyst formation (**D**) and cell death (propidium idodide staining, **E**) in *Naegleria gruberi* in a manner dependent on the presence of lipoproteins. (**D**) N = 2-3 replicates, n = 1775-2062 cells total. Two-sided Kolmogorov–Smirnov test. ns = non-significant, ****p < 0.0001. (**E**) N = 2-3 biological replicates, n = 6-9 technical replicates total, error bars = SEM. Curve fits are variable slope (four parameter) dose response curves. Mixed-effects model. ****p < 0.0001. **(F)** PR-39 (2 uM) colocalizes with fluorescently labelled LDL (10 ug/ml) in *N. gruberi* food vacuoles after 60 min. PR-39 also targets the nucleolus at this time point.

Besides bacteria, AMPs also target parasites. While we had already shown that LL-37 could be endocytosed by *N. gruberi* (**Fig. 2E**), it did not kill them (**Fig. S7A-D**). We moved on to the porcine cathelicidin PR-39 that is more cationic in nature and targeted the nucleolus of these protists (**Fig. 2E**), which we have previously shown to be important for +CPP killing activity^4^. Treating *Naegleria* with PR-39 rapidly induced them to switch from their amoeboid state into a stress-associated cyst-like state (**Fig. 6C-D**). At higher PR-39 concentrations, these cysts stained positive for propidium iodide, indicative of their death (**Fig. 6C, E**). As these amoebas are axenically cultured with serum as a food source, we repeated our experiment with serum from which lipoproteins were depleted (lipo-depleted). This significantly *reduced* toxicity (**Fig. 6C-E**). Contrary to bacteria, these protists endocytose lipoproteins as a major energy source in the host environment. Thus, the binding of AMPs like PR-39 to lipoproteins could accelerate their uptake and boosts their killer activity. Indeed, when we co-treated these cells with fluorescently labeled LDL particles, we found that they co-localized with PR-39 in the vesicle-like structures we had seen above (**Fig. 6F**, **Fig. 2E**), confirming that these are *bona fide* food vacuoles.

Together, these findings show that physiologically relevant levels of lipoproteins modulate the toxicity of AMPs and that the direction of the effect is dependent on whether the target pathogens endocytose lipoproteins or not (**Fig. 6B**).

## DISCUSSION

Despite three decades of intense study, the mechanisms governing the uptake of cationic cell-penetrating peptides (+CPPs) have remained controversial. Many studies have provided often conflicting evidence on the main routes of cellular uptake, being either passive or active. We attribute the discrepancies to countless experimental variables between these studies that often may not represent well their native context. A major one is the reliance on high peptide concentrations that are often orders of magnitude higher than one would expect in nature (**Fig. S1A**, **Table S1**, **Supplemental Text A**). Additionally, the “monoculture” of experimental systems, with most studies using just a handful of synthetic peptides in a handful of cancer cell lines (**Table S1**), limits our ability to assess which mechanisms may be shared across natural +CPPs and relevant target cells and organisms. To address these current limitations, we decided to take a different approach, where we (1) aggregated a representative library of natural +CPPs from 17 species spanning 600 million years of animal evolution, (2) paired several of them with their physiological targets, and (3) performed our assays at physiologically relevant concentrations. In total, our study tests twenty +CPPs in fifteen human cell lines/types and five additional organisms.

We found that endocytosis is the predominant mode of uptake of natural +CPPs at nanomolar concentrations and across cell types and organisms (**Fig. 1-2**). While we do not rule out the existence or potential contributions of passive mechanisms, we propose they may be largely limited to high +CPP concentrations that destabilize the membrane (**Fig. S1A**). Even though +CPP concentrations can acutely spike to such high levels (e.g., consider the local crotamine concentration directly after a snake bite), these peptides are typically present at substantially lower concentrations or rapidly become diluted within tissues and biofluids (**Supplemental Text A)**.

To highlight that the endocytosed pool of +CPPs is the functionally relevant pool, we show that +CPPs traffic through and eventually escape from the endolysosomal pathway (**Fig. 3**) before accumulating at their site of action (e.g., the nucleus^4^). This is an important point as the endolysosomal accumulation has been suggested as a site of toxin neutralization through degradation, rather than a relevant trafficking route. In line with this, numerous other biological, pathological and therapeutic entities undergo endolysosomal escape^80,81^. These include viruses^100^, infectious prions and prion-like aggregates^101^, antisense oligonucleotides^102^, siRNAs^103^, mRNA vaccines^104^, and transfected plasmids^105^. As the biophysical mechanism of this process remains elusive, we propose +CPPs as a useful model system for its interrogation in the years to come. A major advantage of these peptides is that we have exquisite control over their precise (sequence) chemistry via synthesis.

Even though endocytosis had been demonstrated as a possible mechanism for +CPP uptake, we lacked the identity of the implicated receptors. While polyanions on the cell surface had already been implicated, these are merely co-receptors that dock peptides to the cell rather than initiating endocytosis themselves^35–37^. Identically, viruses have evolved similar entry mechanisms, where electrostatic tethering to the glycocalyx promotes infection by reducing the search for the endocytic entry receptor from a 3D to a 2D problem^106^. By leveraging genetic screens, multiplex imaging, and a range of cellular, biochemical and biophysical assays, we provide evidence for a mechanism implicating lipoprotein receptors (**Fig. 4**) that is conserved across animals and even extends to their protist pathogens. This +CPP uptake mechanism seems to have repeatedly and convergently evolved across the eukaryote lineage. As all animals use lipoprotein particles to share and distribute lipids amongst their cells and tissues, these particles are recognized by a variety of lipoprotein and related receptors that endocytose them. Additionally, single-celled protist pathogens and symbionts have evolved similar receptors to steal lipoproteins from their animal hosts as a food source. Thus, lipoprotein uptake constitutes one of the most common and busiest endocytic routes across many eukaryote species, making it an ideal target for +CPPs.

As lipoprotein receptors bind to cationic apolipoproteins embedded in the surface of lipoprotein particles, one could expect that our +CPPs bind the anionic ligand binding domains to hijack these receptors (i.e., receptor hijacking). Yet, based on our cellular assays, +CPPs and lipoproteins do not seem to compete for receptor binding. This could mean that +CPPs may bind accessible receptor domains not bound to the lipoproteins. However, we find strong evidence that the most likely explanation is one of lipoprotein hitchhiking, where the +CPPs first bind lipoproteins (**Fig. 5**), which subsequently control their uptake. This was the case for the three main human lipoprotein species (i.e., HDL, LDL and VLDL). Even though our genetic screen only identified LDLR as a receptor hit (**Fig. 4**), this is likely a cell type-dependent effect. Lipoproteins can be taken up via their canonical receptor (e.g., LDL via LDLR), but more often than not by a host of other lipoprotein and scavenger receptors. Our experiments using HepG2 LDLR knock-out cells confirm this. While these cells had reduced LDL and +CPP uptake (**Fig. 4F-G**), loss of LDLR did not completely prevent their uptake. Performing transcriptomics analysis, we find that these knock-out cells strongly upregulated lipid metabolism genes (fold enrichment = 2.6, p-value = 1.77E-10), including several lipoprotein and scavenger receptors, as a likely compensatory mechanism (**Table S5-7**, **Supplemental Text D**). This finding indicates that several receptors may accommodate +CPP uptake, as long as they are compatible with lipoprotein uptake. Further evidence for this interpretation comes from our protist studies. Also here, we found that +CPPs are co-endocytosed with lipoproteins, even though protist lipoprotein receptors are evolutionary unrelated to animal LDLR. While our data does not rule out a role for other classes of endocytic receptors and cargoes, we provide compelling evidence that lipoproteins and their receptors are major players in +CPP uptake that are shared across a divergent set of eukaryote lineages. Our study also provides a mechanistic explanation as to why serum addition was previously observed by others to modulate +CPP uptake^8,20,22^.

In line with our results, available proteomics data indicates that human +CPPs are natively bound to lipoproteins in the bloodstream^94–97^, which has also been confirmed in disease contexts^98,107–109^. Hyperlipidemia, or the elevated serum levels of “bad cholesterol” in the form of LDL particles, is one of the most common co-morbidities in Western populations (i.e., 1 in 3 of US adults)^110–112^. This urges us to explore if the association of lipoproteins with +CPPs may affect not only their uptake but also their canonical function. As a case study, we focused on the cathelicidin family of AMPs (**Fig. 6**). In humans and livestock these peptides are produced in response to infection. LDL, at levels that were two orders of magnitude below the physiological serum concentration, strongly inhibited bacterial killing as the lipoproteins competed for +CPP binding with the bacterial membrane. However, for protist pathogens that are dependent on lipoprotein uptake as an energy source, we found that the presence of lipoproteins boosted the antimicrobial activity of +CPPs. Here, lipoprotein hitchhiking would promote the protist’s uptake of AMPs. Thus, the native presence of AMPs on lipoproteins in the bloodstream may provide a mechanism for animals to essentially weaponize these particles as they are a critical food source of invading pathogens. While future work should evaluate this mechanism in *in vivo* models, our findings could already have immediate implications in drug delivery. Some of our first-line drugs against diseases like trypanosomiasis suffer from off-target effects and poor pathogen targeting and efficacy^113^. Primary amebic meningoencephalitis remains largely untreatable and near universally fatal^114^. One could imagine leveraging custom lipoprotein-like delivery vehicles that would direct these drugs to the parasites (and the liver for detoxification of the off-target fraction while sparing other tissues). Our finding that cathelicidins from other mammals are more potent at killing protist parasites may also lay the foundation for rational design studies that further enhance the clinical potential of such peptides.

In summary, our study builds on the incredibly rich literature of the cell-penetrance field as it takes prior observations on both passive and active uptake mechanisms and interrogates them across physiologically relevant experimental conditions and model systems. In doing so, we demonstrate endocytosis as a physiological *and* functionally relevant +CPP uptake route. Next, we provide a unified mechanistic framework to understand uptake—in part—as the result of lipoprotein hitchhiking. Our mechanism explains observations by others regarding the modulation of +CPP uptake by serum^8,20,22,99^, and identifies the biophysical interactions responsible. We then extend it to clades that span the eukaryote lineage. In addition to providing important insights into the biology of +CPPs, our study may also motivate moving beyond model system monocultures and leverage evolutionary diversity as a guide to identify conserved/convergent mechanisms. Lipoprotein hitchhiking could be specifically relevant for neurodegenerative diseases, as we know that lipoprotein metabolism is deeply implicated in their genetics and pathophysiology^115–117^. Additionally, cationic cell-penetrating AMPs, including LL-37, have been recently implicated in these disorders^70,118–120^. Besides the *C9orf72* cationic repeat peptides in this study, emerging evidence indicates that lipoprotein and scavenger receptors facilitate the uptake of other pathological protein aggregates^121,122^, enabling their prion-like spreading. Future work should attempt to further disentangle the role of the lipoproteins themselves (and APOE risk/protective variants) in this process and explore whether cationic AMPs also contribute to this. Some of the insights gained here address why tissues and cell types with high rates of lipoprotein uptake are preferentially targeted by +CPPs *in vivo*. For example, in mice, crotamine was shown to preferentially accumulate in liver and kidneys^123^, tumors^124^, and target adipose tissue^125^. Lastly, our findings on the uptake mechanisms of human +CPPs and the increased antiparasitic activity of orthologous sequences may guide the future design of novel peptide-based drug delivery vehicles and therapeutics.

## LIMITATIONS OF THIS STUDY

In this study we have aimed at capturing a broad range of eukaryote peptides and target species. Obviously, it is impossible to cover the entire lineage. Therefore, we have focused on eukaryote target species lacking a cell wall, which would provide an additional barrier against +CPP entry. To gather a large enough library of peptides we also opted for chemical synthesis, as cationic peptides are toxic when produced in bacteria and are hard to purify without anionic contaminants like RNA and lipopolysaccharide that would affect their uptake and toxicity. Thus, we mostly limited ourselves to linear peptides given (1) the difficulties of synthesizing folded peptides and (2) the fact that for most peptides from non-traditional model organisms we do not even have an experimentally validated structure. To address this, future work could be focused on testing our findings with additional folded peptides and include cells from eukaryote lineages that have cell walls. Even though we find that lipoproteins are an important carrier for +CPPs, these particles are not equally abundant in all biofluids. We do not rule out that other hitchhiking targets may be identified in other biofluids or other organisms. Lastly, while we find that several of our +CPPs undergo endolysosomal escape, the biophysical mechanism by which they do so remains incompletely resolved. Identifying it is well beyond the scope of this study. Thus, we hope that the approaches, reagents, and model systems developed in our work may prove a useful starting point for the field to start interrogating the evolutionary and molecular basis of endolysosomal escape.

## ACKNOWLEDGEMENTS

The authors would like to thank all members of the Boeynaems lab, the Gitler and Holehouse labs, the Water and Life Interface Institute (WALII), THINC, CAND, NRI, IDPSIG, and FNZ for helpful discussions and feedback. We would like to thank Furqan Fazal, Stephanie A. Pangas, Jason E. Lee, Thomas A. Cooper, Ignatia Van den Veyver, Joshua M. Shulman, Nora Vanegas Arroyave, Lindsay Burrage, Hugo Bellen, Brendan Lee, and Huda Y. Zoghbi for guidance, mentorship, and feedback. We would like to thank Margarite Divenko and Jason C. Mills for sharing reagents and Tyler Jackson and Hongjie Li for their thoughtful input. We would like to thank Joshua A. Riback for helpful suggestions on the manuscript. Our eternal gratitude goes to Olga Rybina-Willis for her logistical and administrative support and for being the best admin, colleague and friend anyone could have ever wished for. This work is dedicated to her memory. **S.R.T** would like to thank Jeremy Purvis and Wayne Stallaert for their mentorship and encouragement to pursue a scientific career and David Nelson for his support in the Cancer and Cell Biology graduate program. **S.B.** is a CPRIT Scholar in Cancer Research. Work in the **S.B.** lab is supported by CPRIT (RR220094), NSF (DBI # 2213983, WALII), NIH (DP2NS142714, R01NS138605), DOW CDMRP (HT94252510231), the Kleberg Foundation, the Welch Foundation, the Rainwater Foundation, the CurePSP Foundation, the Association for Frontotemporal Degeneration, and the Frick Foundation. **S.R.T.** has been supported by NIH training and education grants to BCM (Initiative for Maximizing Student Development Program R25GM056929-26, Cancer and Cell Biology Graduate Program T32GM136560, and Clinical Translational Research Program T32GM136554).

**O.M.S.C.** is supported by the NSF Postdoctoral Research Fellowships in Biology Program under Grant No. 2410403. **S.J.L**. lab is funded by the NIH (R35GM151999). **M.S.** and **S.J.L.** are partially funded by 2021FIB-41 from the Arnold and Mabel Beckman Foundation. CryoEM data was collected using equipment purchased under CPRIT Core Facility Award RP190602. **E.K.S.M.** is a Pew Biomedical Scholar and a CPRIT Scholar (RR230015). **M.A.G.** and **C.W.J.** are supported by grants from the Welch Foundation (10006508) and CPRIT (RR210066). **C.W.J.** is a CPRIT Scholar in Cancer Research. **R.R.A** is funded by 1F31NS145624 from the NIH. The generation of the cell lines and our work has been supported by the grants: La Caixa Foundation (LCF/PR/HP23/52330032) and EU Horizon RIA: 101155885-2, FH-EARLY. **M.A.F.H.** and **L.P.** are/were funded by FAPESP (Fundação de Amparo à Pesquisa do Estado de São Paulo) [#2022/00527-8; 2019/08287-3], FINEP (04.16.0054.02), CAPES, and CNPq. This project was supported by the Texas Medical Center Digestive Disease Center (P30 DK056338) GEMS Core and the BCM 3D Organoid Core. This project was supported by the Cytometry and Cell Sorting Core at Baylor College of Medicine with funding from the CPRIT Core Facility Support Award RP240432 and the NIH (CA125123, OD036336, and OD038251), and the assistance of Joel M. Sederstrom.

## SUPPLEMENTAL TEXT

### A. Estimating the physiological concentration of +CPPs

While sensitive assays (e.g., ELISA, MSD, quantitative mass spectrometry) exist, estimating the physiological levels of peptides remains challenging. This is especially true when one tries to measure the true *local* concentration at their site of action. Peptides can often only be assayed in the biofluid they were produced in or dilute into—giving us upper and lower ranges, respectively. Here, we will discuss three of the peptides that we have used across experiments in this study:

**(1) Crotamine** its concentration in venom ranges from 2-20 mM^126,127^ and a rattlesnake will typically inject between 0.1 - 0.4 ml of venom, before it dilutes into the surrounding tissue/biofluid. This concentration is five orders of magnitude higher than what we typically test in this study, and therefore, represents an upper range of concentrations that have physiological relevance.
**(2) LL-37** its concentration spikes at 1-4 µM in sputum^128,129^ and 33-77 nM in serum or saliva^130–132^. Yet, when secreted by immune cells this concentration will be substantially higher (allowing it to kill *E. coli* and *Naegleria* in the 5 to 15 µM range, respectively; see **Fig. 6**) before diluting into these biofluids. Note that LL-37 can already activate immune cells (via an endocytic mechanism) in the <1 µM range^133^.
**(3) Poly-GR** is found at 1.5 nM in bulk ALS patient brain^134^. But only five poly-GR inclusions (each <10µm^2^) are found per mm^2^ (=10^6^µm^2^) of cortex^135^. Thus, the local concentration in the insoluble inclusion may reach up to ∼30 µM. The soluble pool that dilutes into the cerebrospinal fluid is much lower, down to ∼2 pM^136^. Note that we estimated molar concentrations using a GR30 peptide for comparison with the other +CPPs. In reality, the length, and therefore molar mass, of individual GR peptides in patient material remain undetermined and are likely heterogeneous.

These three peptides provide good case studies of upper, intermediate and lower ranges—spanning no less than nine orders of magnitude, from mM to pM. While venom peptides can be produced at quite high molar concentrations, for most antimicrobial peptides secreted in biofluids this is substantially less. The free soluble pool of neurodegenerative disease peptides that dilute and circulate into the interstitial and cerebrospinal fluid is even much lower.

Based on these estimates, in this study, we have focused our experiments on the following concentrations ranges for +CPPs:

- **Uptake experiments:** 250 nM – 2 µM (a non-toxic range for most cell types and peptides)
- **Toxicity experiments:** 4 - 32 µM

### B. Diverse biological pathways regulate crotamine uptake

In this study, we set out to identify the endocytic receptors that mediate +CPP uptake. Through a CRISPRi screen, we singled out LDLR, whose importance was supported by other genetic hits and biochemical, biophysical and cell biological assays. Yet, the screen returned many more hits that centered around a range of biological pathways (**Fig. S4B-D**). Here, we will discuss some of the major ones and provide potential explanations for their role in +CPP uptake. While beyond the scope of this work, these pathways may form interesting starting points for follow-up studies.

**(1) Ion channels:** Knock-down of two ion channels, ATP1B3 and KCNK5, reduced crotamine uptake. Both of these were previously identified to suppress the toxicity of the +CPPs Tat^21^ and PR20/GR20^31^ (**Fig. S4B**). ATP1B3 its co-subunit, ATP1A1, was also a hit in our screen. The Tat study^21^ proposed a model where these ion channels are involved in generating the membrane potential required for the passive translocation of Tat peptides. While such spontaneous translocation was observed in the referenced study, at lower concentrations or at earlier time points, the predominant uptake mode was endocytosis^21^. Thus, it remains to be shown if the activity of these channels also modulates endocytosis or solely alternate passive mechanisms.
**(2) Mitochondria:** Knockdown of 66 mitochondrial genes reduced crotamine uptake (19% of suppressor hits; p-value = 1.70E-13; **Fig. S4C**). Twelve of these were specifically implicated in the mitochondrial respiratory chain complex I. Endocytosis is an ATP-dependent active process, therefore requiring proper mitochondrial function. Indeed, others have shown that the pharmacological inhibition of mitochondrial respiration perturbs +CPP uptake^22^.
**(3) Peroxisomes:** Knockdown of 11 peroxisomal genes reduced crotamine uptake (3% of suppressor hits; p-value = 1.91E-05; **Fig. S4C**). Peroxisomes serve important functions in lipid metabolism. Recently, it was shown that peroxisomes are needed for proper cholesterol trafficking^137^. After LDL particles are endocytosed, peroxisome-lysosome contacts are required to regulate cholesterol export from the lysosome and direction to the plasma membrane. If peroxisomes are dysfunctional, they will accumulate cholesterol in their lysosomes, essentially starving cells from cholesterol. While this lack of available cholesterol will drive increased LDLR expression via SREBP2^138^, the reduction in plasma membrane cholesterol also interferes with proper endocytosis^139^. Thus, despite the compensatory transcriptional mechanism that would promote LDL uptake, peroxisomal dysfunction is nonetheless expected to result in a net decrease in endocytosis.
**(4) Splicing factors:** Knockdown of 16 nuclear speckle genes, 7 of which were subunits of the U12 spliceosome, increased crotamine uptake (9% of enhancer hits; p-value = 3.49E-06; **Fig. S4C**). The so-called minor spliceosome regulates a specific type of introns found in just over 600 genes^140^. For two of the U12 splicing factors, ZCRB1^141^ and ZRSR2^142^, we found datasets on which genes are misspliced upon their loss of function. Interestingly, enhancer hits in our knockdown screen were significantly enriched in genes that depended on ZCRB1 (fold enrichment = 2.5; p-value = 0.03) and ZRSR2 (fold enrichment = 2.5; p-value = 0.01) for proper splicing. This was not the case for suppressor hits. Thus, the knockdown of U12 genes seems to preferentially deregulate genes that increase +CPP uptake when knocked down themselves (**Fig. S4D**). Whether there is a biological reason for this enrichment is unknown.

### C. LDLR expression is tightly regulated at multiple levels

LDLR occupies a central role in regulating lipid metabolism by mediating the uptake of LDL particles from the extracellular environment. Unsurprisingly, this process is both tightly regulated and, given its role in metabolic disease, intensely studied. Taking a deep dive into the literature, we found evidence for several of our screen hits being directly implicated in the regulation of LDLR levels and its function (**Fig. 4D**). Here, we will discuss each of these regulatory processes.

**(1) mTOR signaling:** A running joke in cell biology is that mTOR regulates everything. Since LDLR serves a prominent role in controlling the uptake of cholesterol, its transcription is tightly regulated by energy and nutrient levels. mTOR inhibition represses *LDLR* expression and the contrary is true as well^143^. We identified 13 mTOR subunits and regulators. Knockdown of subunits from the mTOR complex (i.e., MLST8, RPTOR and RICTOR) or its positive regulator Ragulator (i.e., LAMPTOR3 and LAMTOR4) decreased crotamine uptake. On the other hand, subunits of the negative regulators were enhancers, including TSC (i.e., TSC1), KICKSTOR (i.e., C12ORF66, ITFG2, KPTN and SZT2) and GATOR1 (i.e., DEPDC5, NPRL2 and NPRL3) complexes. Lastly, FKBP1A, the actual target of rapamycin, was also a hit. The gene knockdown effects on crotamine uptake of each of the above subunits correlated with their effect on mTOR activity and therefore are likely explained by their effect on *LDLR* expression.
**(2) *LDLR* gene expression.** We identified two transcription factors (i.e., SP1 and SP3^144^) and two splicing factors (i.e., SF3B1 and SF3B2^145^) that regulate *LDLR* expression/splicing. Two additional RNA-binding proteins increase (i.e., GRSF1^146^) or decrease (i.e., ZFP36L2^147^) its RNA stability. Again, the direction of their effect on crotamine uptake follows their effect of *LDLR* levels.
**(3) LDLR trafficking.** To exert its function, LDLR needs to be presented on the cell surface. First, upon translation, it needs to be imported into the endoplasmic reticulum. This is mediated by the Sec61/TRAP complex of which several subunits were suppressors (i.e., SSC1, SSC2, SCC3, SEC61B and SEC61G). This complex is known as important for LDLR surface expression^148^. Three other hits involved in exocytosis were recently also implicated in LDLR trafficking (i.e., EXOC3, RALGAPB and SNAP23^89^). Effects were congruent with LDLR surface expression.
**(4) LDLR receptor complex:** Besides LDLR, we identified its co-receptor LDLRAP1^149^. As mentioned in the main text, IDOL was an enhancer in our screen. This is consistent with its role as a mediator of LDLR degradation. Interestingly, SENP1 regulates the levels of IDOL^150^ and its knockdown had, as expected, the opposite effect on crotamine uptake.

In conclusion, at least 32 hits in our screen can be directly tied to the regulation of each aspect of LDLR expression. Together with the overlap between our hits and hits from an LDL uptake screen^89^ and the biochemical, biophysical and cell biological evidence that we provide, these results highlight the central role of the LDLR endocytic pathway in the uptake of +CPPs.

### D. Loss of LDLR function drives compensatory endocytic pathways

In our CRISPRi screen in K562 cells, we identified LDLR as the main endocytic receptor hit. But is this the sole receptor that can accommodate lipoprotein-mediated +CPP uptake? We expanded our experiments to HepG2 cells, which are commonly used in the field of LDLR biology, to investigate this. Knocking out LDLR strongly reduced but did not eliminate +CPP uptake. Compellingly, the same was true for LDL itself (**Fig. 4F-G**). This argues that additional receptors can endocytose LDL and its bound +CPPs. RNA sequencing analysis on these LDLR knock-out cells showed the upregulation of genes that are implicated in the uptake of LDL or hijacked by viruses and peptides.

**LDLRAD1 (Low density lipoprotein receptor class A domain containing 1):** Lipoprotein receptor family member. Untested role in lipoprotein uptake but has been linked to coronary heart disease disease in a GWAS study^151^.

**LDLRAD3 (Low density lipoprotein receptor class A domain containing 3):** Lipoprotein receptor family member. Untested role in lipoprotein uptake but is hijacked by amyloid-beta fibrils^122^ and several viruses^152,153^ for cell entry.

**SCARA3 (Scavenger receptor class A member 3):** Implicated in the endocytosis of peptide-DNA nanoparticles^154–157^.

**CD14 (Cluster of differentiation 14):** Involved in the detection and uptake of (modified) LDL^158,159^.

**GPC3 (Glypican 3):** Acts as a competitive inhibitor of PCSK9^160^, the latter which promotes the degradation of LDLR and other lipoprotein receptors.

## SUPPLEMENTAL TABLES

**Table S1: Literature review on experimental conditions in +CPP studies.**

**Table S2: Peptides used in this study.**

**Table S3: CRISPRi screen hits: suppressors.**

**Table S4: CRISPRi screen hits: enhancers.**

**Table S5: RNA sequencing of HepG2 LDLR KO cells.**

**Table S6: GO term enrichment: upregulated genes in HepG2 LDLR KO cells.**

**Table S7: GO term enrichment: downregulated genes in HepG2 LDLR KO cells.**

**Table S8: Microscale thermophoresis & spectral shift analyses.**

**Table S9: Recombinant DNA sequences.**

## SUPPLEMENTAL VIDEOS

**Video S1: 3D Cryo-ET rendering of nanogold-labeled GR30 bound to the LDL surface.**

**Video S2: 3D Cryo-ET rendering of nanogold-labeled LL-37 bound to the LDL surface.**

**Video S3: 3D Cryo-ET rendering of nanogold particle not bound by LDL.**

**Video S4: 3D Cryo-ET rendering of nanogold-labeled GP30 not bound by LDL.**

## METHOD DETAILS

### Mammalian Cell culture

Cells were maintained in 37C and 5% CO2 sterile incubators. All cells were tested for mycoplasma contamination monthly. Cells were maintained at 70% confluency before each passage and passaged at least once before subsequent experiments after thawing. Adherent cell lines were maintained by first rinsing cells with 1X PBS and then treating cells with 0.25% 1X trypsin-EDTA (Gibco) for a maximum of 5 minutes at 37C before quenching the reaction with complete media. Immortalized adherent cells used here included HEPG2 (Simon Pfisterer lab), HEK293T (ATCC, CRL-3216), U2OS (ATCC, MSPP-HTB96), HMC3 (ATCC, CRL-3304), and HFL1 (ATCC, CCL153), and suspension cells included K562 (ATCC, CCL-243) cells. All cells were maintained in media with FBS and penicillin–streptomycin as suggested by manufacturer.

### +CPP treatment in-vitro, peptide synthesis and click chemistry

All peptide treatments were prepared using protein lobind deepwell plates (Eppendorf, #951032107) or snap-tubes (Eppendorf, #022431064) to reduce binding artifacts. All cellvis glass-plates (P96-1.5H-N) were treated with poly-D-lysine (Sigma-Aldrich, P6407) unless stated otherwise. Peptides were resuspended in ddH2O at 1000x concentrations and diluted in the relevant media. All peptide synthesis was conducted through Biosynth. All were synthesized at >90% purity and HCL exchanged and prepped at 10mg scale in 1mg aliquots (dry). All peptides were then resuspended in diH2O and stored at -20C with minimal freeze thaws. All click chemistry reactions were done using Click-iT™ Plus Alexa Fluor™ 488/555/647 Picolyl Azide Toolkit (Invitrogen, #C10641) with the manufacturer’s instructions. After optimization, a 1:1 ratio of CUSO4:copper protectant and 500 nM target concentration of azide dye was used for all in vitro experiments. For labeling of *in vivo* tissue, 1 uM azide dye was used. However, the azide dyes were substituted for AZDYE 647 picolyl azide, AZDYE 488 picolyl azide, and AZDYE 555 (vector laboratories, #CCT1300, CCT-1276-1, CCT-1288R). All images included a no-peptide negative control subjected to the click reaction to assess background-dye staining. All uptake studies ranged from 125 nM -2 μM and is stated in the figure legends or main text.

### Normal immunofluorescent staining

For immunofluorescence staining, immortalized, ipsc-derived and organoid monolayer cells were fixed with 4% paraformaldehyde (Electron Microscopy Sciences, 15710) for 20 min and washed three times with 1× phosphate-buffered saline (PBS). Cells were permeabilized and blocked for 1 h in immunofluorescence blocking buffer containing 5% goat serum, 0.5% bovine serum albumin (BSA) (Tocris, 5217), and 0.4% Triton X-100 (Sigma-Aldrich, T8787) in PBS. Primary antibodies, listed in STAR Methods and also throughout the methods text, were diluted in blocking buffer and incubated with the cells overnight at 4°C. Cells were subsequently washed three times with 1× PBS for 5 min per wash on a shaker and incubated with Alexa Fluor Plus 488–conjugated, highly cross-adsorbed goat anti-rabbit IgG (H+L) (Thermo Fisher Scientific, A32731), and/or Alexa Fluor Plus 647–conjugated, highly cross-adsorbed goat anti-mouse IgG (H+L) (Thermo Fisher Scientific, A32728TR), and Hoechst 33258 (Sigma-Aldrich, 94403). Cells were then washed three times with 1× PBS for 5 min per wash before imaging. For mouse tissue, after sectioning onto Fisherbrand Superfrost Plus microscope slides (Fisher Scientific, 22-037-246), tissue was rinsed with 1x PBS once. Sections for staining were delineated using a pap pen and tissue was incubated with 4% paraformaldehyde for 5 minutes followed by three subsequent 5 minute 1x PBS washes. Tissue was incubated with blocking buffer mentioned above for 1 hour and primary antibodies were diluted in blocking buffer overnight at 4°C. Tissue-specific secondary steps followed the same procedure as cells followed by mounting onto 1.5# glass with prolong gold antifade (Invitrogen, P36930).

### Flow cytometry Endocytosis inhibitor

K562 cells (1 × 10^5 cells per condition) were treated with the indicated endocytosis inhibitors, including dynasore (80 µM) (EMD Millipore, 324410-10MG), EIPA/amiloride (500 µM) (SigmaAldrich, A7410-1G), Pitstop 2 (20 µM) (Sigma-Aldrich, SML1169-5MG), genistein (150 µM) (G6649-5MG), with matched DMSO vehicle controls in Eppendorf lobind 500 µL plates. Cells were incubated in medium supplemented with each inhibitor for 30 minutes, before addition of the alkyne-labeled or cy3-labeled peptide for 30 minutes. Inhibitors were maintained in the medium throughout the 30-min peptide treatment. Cells were subsequently washed with 1X PBS, fixed with 4% paraformaldehyde for 20 minutes, and subjected to copper-catalyzed azide–alkyne click chemistry using Azdye-555. Following labeling, cells were washed, resuspended in 200 µL of cell-staining buffer (Biolegend, #420201) in plastic 96-well plates (Genesee Scientific, 25-109), and analyzed by flow cytometry using an Attune Cytpix Flow Cytometer (Invitrogen, # A51844). Cells were gated based on ssc and fsc singlets for live cells and the unstained control for autofluorescence. Downstream analysis utilized FlowJo 10.10.0 to gate cells based on ssc, fsc and unstained control. Mean, median, top 10%, count, and other parameters were extracted per well into csv’s from flowjo software. Plotting and analysis of the concatenated data was done in python using custom-built scripts.

### S2R+ Drosophila melanogaster cell culture

*Drosophila melanogaster* S2R+ cells (Drosophila Genomics Resource Center, DGRC) were maintained at 25°C in Schneider’s Drosophila medium (Thermo Fisher Scientific, #21720024) supplemented with 10% heat-inactivated fetal bovine serum (Corning, #35-016-CV) and 1% penicillin– streptomycin (Thermo Fisher Scientific, #15140122). Cells were cultured under ambient atmospheric conditions without supplemental CO₂ and passaged every 3–4 days. For routine passaging, cells were detached by gentle pipetting or scraping, resuspended in fresh complete medium, and reseeded at a 1:8–1:10 dilution. The S2R+ cell lines used in this study were S2R+-Rab11GFP-14 (RRID: CVCL_UW09) and S2R+-Arl8-GFP-4 (RRID: CVCL_XF61), obtained from the DGRC.

### *Trypanasoma cruzi* cell culture and imaging

*Trypanosoma cruzi* H1 strain parasites, originally isolated from human cases in Yucatán, Mexico ^161^ were maintained in Vero cell cultures (ATCC CCL-81). Vero cells were cultured in MEM (Gibco) and 10% FBS (Gibco, 12492013) at 37°C with 5% CO₂ and were infected at 80% confluency and at a multiplicity of infection of 5. Parasites were collected from the infected cultures every 2–3 days and used immediately for experiments. To isolate trypomastigotes, the media was collected from infected mammalian cells. The media was centrifuged at 350 g for 5 minutes to pellet mammalian cell debris and the supernatant was then further centrifuged for 7 minutes at 3000 g to pellet the parasites. Because the trypomastigote pool contained sparse amastigotes (identified by morphology), likely from prematurely ruptured cells, we also assessed peptide localization in these cells. Each parasite collection represented independent experimental preparation and was not returned to the maintenance culture. 500,000 parasites were treated in each 96-well condition. Parasites were treated and prepped for imaging in PDL-coated cellvis glass plates previously mentioned and fixed with 4% paraformaldehyde for 20 minutes. After 20 minutes, plates were spun down 3000 g for 5 minutes to improve retention of the parasites to the plate. After fixation, cells were permeabilized with 0.2% PBS-T for 5 minutes and previously described click reaction was administered to visualize peptide with Hoechst staining.

### *Naegleria gruberi* cell culture and experimentation

*Naegleria gruberi* cells (strain ATCC® 30224) were cultured axenically in 25 cm² tissue-culture flasks containing 5 mL of ATCC medium 1034 (modified PYNFH medium), hereafter referred to as growth medium, and maintained at 23°C. For immunofluorescent imaging experiments, cells were seeded at an initial density of 15,000 cells and grown overnight on cellvis glass 96 well plate before peptide treatment and image processing and acquisition. For kill assays, cells were seeded and grown overnight before next-day peptide treatment. At the start of treatment, cell media was changed to media with normal FBS (Corning, 35-016-CV) or lipo-depleted FBS (Kalen biomedical, 880100-2) and imaged at 23°C at specified timepoints with 1 ug/ml of propidium iodide (Thermo Fisher Scientific, P1304MP) diluted in the media. Images were taken at 20x and stitched using zen software (ver. 3.11). CellposeSAM was used to segment cells, napari was used to manually validate the masks, and proofs were generated for each cell body (**Fig. S7A**).

### *Capasaspora owcwarzaki* cell culture and experimentation

Capsaspora owczarzaki cells (strain ATCC® 30864) were cultured axenically in 25 cm² tissueculture flasks containing 5 mL of ATCC medium 1034 (modified PYNFH medium) supplemeted with 30% heat-inactivated FBS, hereafter referred to as growth medium, and maintained at 23°C. For imaging experiments, 500,000 cells were seeded in a cellvis glass 96 well plate, fixed with 4% paraformaldehyde, click-treated and imaged as stated in confocal imaging acquisitions.

### iPSC-derived cortical neuron culture

WTC11 induced pluripotent stem cells (iPSCs) were maintained on Matrigel-coated culture vessels in StemFlex medium (Gibco, #A3349401), with medium replaced every other day. Matrigel (Corning, #354234) was diluted to 100 µg/mL in chilled KnockOut DMEM/F12 (Thermo Scientific, #12660012), applied to culture vessels, and incubated at 37°C for 30–60 min before use. Cells were passaged using Accutase (Sigma-Aldrich, #A6964) collected by centrifugation at 210 × g for 5 min, and replated in StemFlex supplemented with ROCK inhibitor (Selleckchem, #Y-27632); ROCK inhibitor was removed 48 h after plating. For neuronal differentiation, 4 × 10^6 cells were seeded onto a Matrigel-coated 10-cm dish three days before differentiation in KnockOut DMEM/F12 containing NEAA (Gibco, #11-140-050), N2 supplement (Gibco, #17502048), NT3(Gibco, #AF-450-03-100UG), BDNF(Gibco, #AF-450-02-100UG), mouse laminin(Gibco, #23017015), doxycycline(Sigma-Aldrich, #D3447), and ROCK inhibitor, followed by daily half-medium changes without ROCK inhibitor. On day 0, cells were dissociated with Accutase, centrifuged at 200 × g for 5 min, and plated onto poly-D-lysine-coated 96 well glass plates (Cellvis) in neuronal differentiation medium consisting of equal parts DMEM/F12 and Neurobasal-A supplemented with NEAA, GlutaMAX, N2, B27 without vitamin A (Gibco, # 17504044), NT-3, BDNF, mouse laminin, and doxycycline. A complete medium change was performed on day 3, followed by weekly half-medium changes without doxycycline or ROCK inhibitor. On day of peptide treatment, cells were treated with peptide diluted in neuronal differentiation medium for specified times, fixed with 4% paraformaldehyde and processed with click chemistry and normal immunofluorescence protocols using beta-3 tubulin antibody as a cell type marker (Thermo, MA1-118).

### Human airway epithelial stem/progenitor cells (nasal origin) monolayer

Human nose organoid-derived air–liquid interface (HNO-ALI) cultures were generated from tissue-derived human nasal epithelial stem/progenitor cells isolated from paired nasal wash and mid-turbinate swab samples previously described^162^. The Baylor College of Medicine 3D organoid core provided the monolayer. Three-dimensional HNOs were expanded prior to dissociation into single cells using 0.05% trypsin/0.5 mM EDTA (Invitrogen, #25300054) and passage through a 40-µm cell strainer (Falcon, #352340). Cells were seeded onto cellvis glass plates precoated with bovine type I collagen (, Gibco #A1064401). Cells were initially maintained in airway organoid medium supplemented with epidermal growth factor (EGF; PeproTech, #AF-100-15) and 10 µM Y-27632 (Sigma-Aldrich, #Y0503). After 4 days, confluent monolayers were transitioned to air– liquid interface culture with PneumaCult-ALI Medium (STEMCELL Technologies, #05001) supplied to the basolateral compartment and differentiated for 21 days to generate a polarized, pseudostratified nasal airway epithelium. +CPPs were treated for 2 hours and subsequent staining (sox2, cell signaling technology, #14962) and click chemistry was performed.

### Human jejunum organoid monolayer

Human jejunal intestinal enteroid monolayers were provided by the Baylor College of Medicine 3D organoid core. HIEs, derived from tissue-resident intestinal stem/progenitor cells isolated from human jejunal crypts, were maintained as three-dimensional cultures in WRNE medium for approximately 7 days before preparation of epithelial monolayers^163^. Cellvis glass 96 well plates were coated with collagen IV (Sigma-Aldrich, #C5533-5MG) at 33 µg/mL for 90 min at 37°C or overnight. Enteroids were recovered using DPBS without Ca²⁺ or Mg²⁺ (Invitrogen, #14190-144) containing 0.5 mM EDTA, dissociated with 0.05% trypsin-EDTA (Invitrogen, #25300054) for 4–5 min at 37°C, mechanically dispersed, and passed through a 40-µm cell strainer (Falcon, #352340). Cells were resuspended in WRNE medium containing 10 µM Y-27632 (STEMCELL Technologies, #72308) and seeded at approximately 2.5–3 × 10⁵ cells per well in 100 µL apical medium, with 600 µL medium added basolaterally. The following day, WRNE medium was replaced with differentiation medium in both compartments to promote formation of a differentiated jejunal epithelial monolayer. +CPPs were treated for 2 hours and subsequent staining (villin, abcam, ab97512) and click chemistry was performed.

### Human liver organoid monolayer

Human liver organoids were provided by the Baylor College of Medicine 3D Organoid Core. Liver organoid establishment and maintenance were performed as previously described^164. Wells of a Cellvis glass-bottom 96-well plate were coated with 100 µL collagen IV diluted 1:30 in cold water and incubated at 37 °C for 90 min. Dense 3D human liver organoids were released from growth factor-reduced, phenol red-free Matrigel (Corning, Cat# 356231) and dissociated in 300 µL 0.05% trypsin–EDTA at 37 °C for 4 min with intermittent pipetting. Digestion was quenched with two volumes of cold complete medium without growth factors (CMGF−), and the organoids were collected by centrifugation at 80 × *g* for 5 min at 4 °C. The pellet was resuspended in human liver organoid expansion medium and plated at 100 µL per collagen-coated well, using approximately one well of dense 3D organoid culture per monolayer well. Once confluent, monolayers were cultured in expansion medium supplemented with 25 ng/mL recombinant human BMP7 (PeproTech, Cat# 120-03) for 5 days, followed by human liver organoid differentiation medium for 10 days, with medium replaced every other day. Differentiated monolayers were treated with +CPPs for 2 h, after which click chemistry and immunofluorescence staining for albumin (Proteintech, Cat# 16475-1-AP) were performed.

### Human vaginal and bladder organoid monolayers

Vaginal and bladder organoid monolayers were provided by the Baylor College of Medicine 3D organoid core. 96-well plates was coated with 100 µL type IV collagen (30 µg/mL) for 90 min at 37 °C. Three-dimensional HVO or HBO cultures were recovered from Matrigel with cold PBS containing 0.5 mM EDTA, pelleted, and digested with 0.05% trypsin/0.5 mM EDTA for 8 min at 37 °C. Trypsin was neutralized with CMGF containing 10% FBS; cells were mechanically dissociated, passed through a 40-µm strainer, pelleted, and resuspended in HVO or HBOP propagation medium. Cells were plated at 100 µL per insert or well; wells additionally received 600 µL basolateral medium. After 2–3 days, confluent HVO cultures were transitioned to an air–liquid interface by removing apical medium, whereas HBO cultures received HBO differentiation medium in both compartments. Cultures were differentiated for 5 days with medium changes every 2–4 days.

+CPPs were treated for 2 hours and subsequent staining (UPK3a, Progen, 610108) and (CK5, Invitrogen, MA5-17057) and click chemistry was performed.

### iPSC-derived precursor muscle cells

Human pluripotent stem cell-derived skeletal myogenic progenitors were generated and purified by the Darabi laboratory according to the method described by Xu et al^165^. Briefly, skeletal myogenic progenitors were differentiated from human pluripotent stem cells and purified by fluorescence-activated cell sorting for the CD10⁺CD24⁻ population. The purified cells were provided to our laboratory and subsequently cultured as described.

### iPSC-derived endothelial cells

Human-induced pluripotent stem cells (hiPSCs) were differentiated into endothelial cells (ECs) using an optimized protocol adapted from established methods^166–169^. Briefly, hiPSCs were directed toward the mesodermal lineage via treatment with the Glycogen Synthase Kinase 3 beta (GSK3β) inhibitor, CHIR99021, for 2 days, followed by a 2-day endothelial specification phase using a defined combination of cytokines. This protocol yields 50% to 60% CD31⁺CD34⁺ endothelial progenitor populations. The hiPSC-derived ECs were subsequently isolated using cell sorting. Purified hiPSC-ECs expressed canonical endothelial markers—including CD31, von Willebrand factor (vWF), VE-cadherin, and Tie-2—and maintained exponential expansion capacity in vitro^166–169^. +CPPs were treated and subsequent staining (CD31, Thermo Scientific, MA3100) and click chemistry was performed.

### iPSC-derived cardiomyocytes

iPSC-CMs were generated using a modified published GiWi protocol and other published methods^170–173^. Briefly, on day 0 of differentiation, the medium was changed to RPMI 1640 (Thermo Fisher Scientific, 11875119) supplemented with 6 μM CHIR99021 (MCE, CT99021). On day 1, the medium was replaced with RPMI 1640 alone. On day 3, cells were cultured in RBA medium consisting of RPMI 1640 supplemented with 0.5 mg/mL fatty acid-free bovine serum albumin (GenDEPOT, A0100010) and 0.2 mg/mL L-ascorbic acid 2-phosphate (Wako, 321-44823), with the addition of 2 μM Wnt-C59 (MCE, HY-15659). On day 5 and every other day thereafter, the medium was changed to RBA supplemented with 5 μg/mL insulin (Sigma, I9278) to maintain iPSC-CMs. Spontaneous contractions were typically observed by day 7. To further enrich the cardiomyocyte population, cells underwent lactate selection from days 20 to 24 using glucose-free RPMI 1640 (Thermo Fisher Scientific, 11879020) supplemented with 4 mM lactate (MilliporeSigma, L7022). Cells were then maintained with RPMI1640 supplemented with 1 x B27 (Thermo Fisher Scientific, 17504044). All experiments were performed using iPSC-CMs between days 30 and 50 of differentiation. +CPPs were treated and subsequent staining (cTNT, Abcam, ab8295) and click chemistry was performed.

### Mouse husbandry

C57BL/6 mice (4-6 months old) were obtained from the Center for Comparative Medicine rodent colony at Baylor College of Medicine. Experiments used equal number of male and female mice. All mice were housed in the Animal Facility at the Neurological Research Institute at Baylor College of Medicine, under a 12-h light/dark cycle with food and water *ad libitum*. All experiments were conducted in compliance with the Guide for Care and Use of Laboratory Animals (PMID: 21595115) and approved by the Baylor College of Medicine Institutional Animal Care and Use Committee, Assurance number D16-00475.

### Mouse intramuscular injection

All mice received an intramuscular injection to the caudal thigh muscles. Mice were anesthetized with isoflurane. While anesthetized, each mouse underwent tail marking for identification and were then injected into the target muscle to a depth of approximately 2 to 4 mm. The syringe plunger was aspirated to check for inadvertent placement within a blood vessel, and then the injection was delivered. Crotamine (500 nM) was administered for 1 hour and then mouse tissue was prepped as described below. Normal mouse immunofluorescence protocol as followed using the following stain (laminin, santa cruz biotechnology, sc-5985).

### Mouse stereotaxic brain injection

Mice were anesthetized with isoflurane (5% for induction and 3% for maintenance in oxygen) and secured in a stereotaxic apparatus using ear bars. Adequate depth of anesthesia was confirmed by the absence of a pedal-withdrawal reflex, and body temperature was maintained using a heating pad. The scalp was shaved and disinfected with betadine and 70% ethanol. A midline incision was made to expose the skull, and bregma and lambda were identified. The skull was leveled in the anteroposterior and mediolateral planes before determining the injection site. A small hole was drilled above the target region. Crotamine (500 nM) was administered unilaterally at the following coordinates relative to bregma: anteroposterior, +1.0 mm; mediolateral, +1.0 mm; and dorsoventral, −0.8 mm from the dura. A total volume of 2 µL was delivered using a Hamilton syringe fitted with a 27-gauge needle. Following completion of the injection, the needle was left in place for 1 min to permit diffusion and minimize reflux along the injection tract. The needle was then slowly withdrawn over approximately 1min. The incision was closed using sutures, and mice received analgesic in accordance with the approved animal protocol. Animals were allowed to recover in a warmed cage and were monitored until fully ambulatory before being returned to their home cages. After mouse tissue collection and sectioning as described below, normal mouse immunofluorescence (NeuN, EMD Millipore, ABN78 and LAMP1, Santa Cruz Biotechnology, sc19992) was performed.

### Mouse tissue collection and sectioning

Unless otherwise specified, deeply anesthetized animals were transcardially perfused with 4% paraformaldehyde (PFA)/PBS solution. Fixed brains were extracted, followed by overnight PFA fixation and gradient sucrose dehydration. Brains were embedded in OCT (Tissue TEK) and sectioned with a cryostat (Leica) at a thickness of 15μm (attached section).

### Confocal microscopy sample acquisition and image analysis

All images were taken on Zeiss LSM 900. Any images taken and compared for relative intensity were taken at the same acquisition settings using either 63x oil objective or 63x water objective. All images taken for iterative immunofluorescence were taken using the 63x water objective, 0.45 digital zoom, and stitched together using zeiss stitching processing (Carl Zeiss Microscopy). All analyses were performed using custom-built python scripts (python version 3.13), which are linked at https://github.com/sovannytaylor/cpp-manuscript-2026. General packages used include, cellposeSAM, napari, seaborn, matplotlib, and skimage. Briefly, an initial cleanup script was used to convert czi files to numpy arrays. CellposeSAM was then used to mask cells and nuclei followed by manual validation of the masks using napari. After mask creation, depending on the analysis, puncta detection was performed by detecting objects using different thresholding methods. All datasets were normalized to the negative control GP30 for background correction. Lastly, object count, morphology, and intensity of markers within the objects were measured based on condition. Cell-level morphology quantifications were also measured for quality control purposes. All image analysis quantifications, unless stated otherwise, were cell-level based. N refers to the number of independent experiments performed and n refers to the number of cells. All statistical analyses were based on the N statistics, unless stated otherwise. For KS tests, the statistical analyses were based on cell-level distributions for each independent experiment. More information is detailed in figure legends and in the code availability.

### Airyscan processing

Super-resolution fluorescence images were acquired using a Zeiss LSM 900 confocal microscope equipped with an Airyscan 2 detector and an inverted Axio Observer 7 stand (Carl Zeiss Microscopy). Images were collected using a 63×/1.15 NA water-immersion LD C-Apochromat objective with a correction collar and 2.0× digital zoom. Four fluorescence channels were acquired in separate tracks using 405-, 488-, 561-, and 640-nm excitation at laser powers of 2.5%, 2.0%, 0.5%, and 2.0%, respectively. Emission was collected at 400–505 nm for the 405-nm channel, 450–545 nm for the 488-nm channel, 450–620 nm for the 561-nm channel, and 620–700 nm for the 640-nm channel. Detector gain was set to 750 V for the 405-nm channel and 700 V for the remaining channels, with a digital gain of 1.0 and an offset of 0.

Images were acquired at 16-bit depth in bidirectional frame-scan mode using a scan speed of 7, a pixel dwell time of 0.89 µs, and 16-fold line averaging. The pinhole was set to 5.0 Airy units for the 488-, 561-, and 640-nm channels and 5.46 Airy units for the 405-nm channel. Images were acquired at 1,156 × 1,156 pixels, corresponding to a lateral pixel size of 0.043 µm and a field of view of 49.68 × 49.68 µm. Images were acquired in two-dimensional Airyscan super-resolution mode and processed using the automatic Airyscan processing settings in ZEN software, version 3.11 (Carl Zeiss Microscopy).

### 4i (iterative, indirect, immunofluorecent imaging) experimentation

Cells were seeded in glass-bottom plates (Cellvis) coated with poly-D-lysine hydrobromide (200 µg/cm²). Cells were treated with 250 nM Cy3-labeled crotamine (SMARTOX), GR30-click or GP30-click peptide (Biosynth), or an equivalent volume of PBS as the vehicle control. Where indicated, DiI-labeled LDL (LDL-DiI; Kalen Biomedical, catalog no. 770230-9) was added at 10 µg/mL. Treatments were performed for 15, 30, 45, 60, or 90 min and were initiated in a staggered manner so that all conditions were fixed simultaneously (**Fig. S8A-C**).

Unless otherwise stated, all subsequent incubations were performed at room temperature, and samples were washed three times with phosphate-buffered saline (PBS) between steps. Cells were fixed with 4% paraformaldehyde for 20 min and permeabilized with 0.4% Triton X-100 in PBS for 10 min. Samples were stained with Hoechst and inspected for quality control in imaging buffer consisting of 700 mM N-acetyl-L-cysteine (Sigma-Aldrich, catalog no. A7250) in double-distilled water (ddH₂O), adjusted to pH 7.4.

Following quality-control imaging, samples were washed three times with ddH₂O. Hoechst was removed by three successive 10-min incubations with elution buffer containing 0.5 M L-glycine (Sigma-Aldrich, catalog no. 50046), 3 M urea (Sigma-Aldrich, catalog no. U4883), 3 M guanidine hydrochloride (Thermo Fisher Scientific, catalog no. 15502-016), and 70 mM TCEP-HCl (SigmaAldrich, catalog no. 646547) in ddH₂O, adjusted to pH 2.5. Elution was performed with gentle agitation. After elution, post-elution images were taken and wells were assessed for any residual staining leftover (**Fig. S8D**).

Samples were then incubated for 1 h in 4i-specific blocking solution containing 100 mM maleimide (Sigma-Aldrich, catalog no. 129585), 100 mM NH₄Cl (Sigma-Aldrich, catalog no. A9434), and 1% bovine serum albumin (BSA) in PBS. Primary antibodies were diluted as indicated in conventional blocking solution containing 1% BSA in PBS and incubated with the samples overnight at 4 °C. Samples were washed three times with PBS and incubated with the corresponding secondary antibodies and Hoechst for 1 h at room temperature with gentle agitation. After three additional PBS washes, samples were imaged in imaging buffer.

Images were acquired using a Zeiss LSM 900 confocal microscope equipped with an Airyscan 2 detector and an inverted Axio Observer 7 stand (Carl Zeiss Microscopy), using a 63×/1.15 NA water-immersion LD C-Apochromat objective with a correction collar. For each condition, a 2 × 2 tiled field was acquired at 0.45× digital zoom, with a lateral pixel size of 0.220 µm and a bit depth of 16 bits. Images were collected using 405-, 488-, 561-, and 640-nm excitation and the corresponding emission-detection windows. Acquisition was performed in bidirectional frame-scan mode at a scan speed of 8, with a pixel dwell time of 0.76 µs and eightfold line averaging. Laser power and detector gain were optimized separately for each staining round to accommodate differences in antibody intensity and signal abundance. Settings were kept as low as practicable to minimize photobleaching and detector saturation while maintaining detectable signal across the imaged plate. Laser power was generally maintained at or below 2.5%, with a maximum recorded setting of 2.7%, and detector gain was generally maintained at or below 750 V, with a maximum recorded setting of 800 V. Within each imaging round, acquisition settings were held constant across all experimental conditions to permit quantitative comparison. Images were stitched using the default settings for zen stitched processing. Image acquisition was performed using ZEN software, version 3.11 (Carl Zeiss Microscopy).

Antibodies included, Rabbit monoclonal anti-LAMP1 (D2D11) (Cell Signaling Technology, Cat. #9091); mouse monoclonal anti-SC35 (Abcam, Cat. #ab11826); mouse monoclonal anti-nucleophosmin (Abcam, Cat. #ab10530); rabbit monoclonal anti-LDL receptor (Abcam, Cat. #ab314008); rabbit polyclonal anti-PEX19 (Invitrogen/Thermo Fisher Scientific, Cat. #PA5-22129); mouse monoclonal anti-human cytochrome c (clone 28-37AB11) (Invitrogen, Cat. #456100); rabbit monoclonal anti-Rab11 XP (D4F5) (Cell Signaling Technology, Cat. #5589); mouse monoclonal anti-calnexin (GT1563) (Thermo Fisher Scientific, Cat. #MA5-31501); rabbit monoclonal anti-EEA1 (C45B10) (Cell Signaling Technology, Cat. #3288); mouse monoclonal anti-phospho-histone H2A.X (Ser139; D7T2V) (Cell Signaling Technology, Cat. #80312); rabbit monoclonal anti-Rab7 (D95F2) (Cell Signaling Technology, Cat. #9367); mouse monoclonal anti-VLDL receptor/VLDL-R (clone 1H10) (Abcam, Cat. #ab75591); mouse monoclonal anti-DDX6 (Sigma-Aldrich, Cat. #SAB4200837); rabbit polyclonal anti-RAB14 (Novus Biologicals, Cat. #NBP1-84720); mouse monoclonal anti-lamin B receptor/LBR (clone BBmLBR 12.F8) (Abcam, Cat. #ab232731); rabbit monoclonal anti-Rab5 (C8B1) (Cell Signaling Technology, Cat. #3547); mouse monoclonal anti-galectin-3 (A3A12) (Thermo Fisher Scientific, Cat. #MA1-940); rabbit recombinant monoclonal anti-NUP50 (JE63-92) (Thermo Fisher Scientific, Cat. #MA5-44782); mouse monoclonal anti-caveolin-1 (7C8) (Thermo Fisher Scientific, Cat. #MA3-600); rabbit polyclonal anti-SEC23A (Invitrogen, Cat. #PA5-144934); mouse monoclonal anti-p53 (DO-7) (Cell Signaling Technology, Cat. #48818); rabbit polyclonal anti-GM130 (Thermo Fisher Scientific, Cat. #PA5-95727); mouse monoclonal anti-clathrin heavy chain (X22) (Thermo Fisher Scientific, Cat. #MA1-065); rabbit polyclonal anti-p62/SQSTM1 (Proteintech, Cat. #18420-1-AP); mouse monoclonal anti-LAMP2 (H4B4) (Abcam, Cat. #ab25631); rabbit polyclonal anti-LC3B (Cell Signaling Technology, Cat. #2775); mouse monoclonal anti-nuclear pore complex (Mab414) (Abcam, Cat. #ab24609); rabbit monoclonal anti-PDI (C81H6) (Cell Signaling Technology, Cat. #3501); mouse monoclonal anti-PCNA (PC10) (Cell Signaling Technology, Cat. #2586); rabbit monoclonal anti-RCAS1 (Cell Signaling Technology, Cat. #12290); rabbit polyclonal anti-phospho-p21 (Thr145) (Abcam, Cat. #ab47300); rabbit polyclonal anti-α-tubulin (Abcam, Cat. #ab18251); rabbit polyclonal anti-lamin A (Abcam, Cat. #ab26300); rabbit recombinant monoclonal anti-AIF (D39D2) (Cell Signaling Technology, Cat. #5318); mouse monoclonal anti-cyclin D1 (DCS-6) (Santa Cruz Biotechnology, Cat. #sc-20044); goat polyclonal anti-cyclin B1 (R&D Systems, Cat. #AF6000); rabbit monoclonal anti-IGF-II receptor/CI-M6PR (D3V8C) (Cell Signaling Technology, Cat. #14364); rabbit polyclonal anti-TGN46 (Proteintech, Cat. #13573-1-AP); rabbit monoclonal anti-mTOR (7C10) (Cell Signaling Technology, Cat. #2983); rabbit polyclonal anti-galectin-8 (Proteintech, Cat. #10955-1-AP); mouse monoclonal anti-G3BP (Abcam, Cat. #ab56574); and rabbit polyclonal antiC9orf72 (Proteintech, Cat. #22637-1-AP).

The secondary antibodies used were donkey anti-rabbit Alexa Fluor 488 (Thermo Fisher Scientific, Cat. #A-21206), goat anti-rabbit Alexa Fluor 594 (Thermo Fisher Scientific, Cat. #A32740), donkey anti-mouse Alexa Fluor 647 (Thermo Fisher Scientific, Cat. #A-31571), and goat antimouse Alexa Fluor 647 (Thermo Fisher Scientific, Cat. #A32728).

### 4i Image processing and analysis

Raw images from all iterative immunofluorescence (4i) rounds were processed using a custom Python-based analysis pipeline. Images from successive rounds were spatially registered using the DAPI channel as the reference. To maximize registration accuracy, each round was aligned to its temporally adjacent round, thereby minimizing morphological differences between images. Transformations were estimated using the phase cross-correlation algorithm and applied uniformly to all fluorescence channels within the corresponding round. The transformed images were retained without further modification for downstream analysis and quality-control visualization. Prior to registration, a Laplacian-of-Gaussian (LoG) filter was applied to each DAPI image to enhance nuclear boundaries and other structural features, which improved registration performance. Registration accuracy was assessed by visual inspection of DAPI overlays from aligned rounds. Representative overlay and difference images were generated to document registration quality (**Fig. S8A-B, S9A**).

Following registration, fluorescence channels were background-subtracted using Gaussian filter from scipy package, using sigmas determined by morphology features to reduce local pixel noise while preserving punctate structures. The same preprocessing procedure was applied consistently to images within each plate and fluorescence channel.

Cell boundaries were initially segmented from the aligned and background-subtracted α-tubulin image acquired during round 16 using Cellpose SAM. The resulting segmentation masks and corresponding image arrays were manually reviewed and edited in Napari (version 0.6.1). Masks were corrected to separate merged cells, remove spurious objects, and exclude improperly segmented cells. Cells intersecting an image boundary or failing predefined image-quality criteria were excluded. Cells were additionally required to be identifiable in the peptide-acquisition round (round 0) and in all immunofluorescence rounds required for the analysis and edge cells were excluded. Cell identities were propagated across the registered image series so that measurements from different rounds could be assigned to the same cell. Quality-control images were generated showing retained and excluded cell masks, and cell-retention rates were recorded at each filtering step (**Fig. S9A**).

### Marker and peptide-punctum segmentation

Marker-positive structures and peptide puncta were segmented separately from the aligned, background-subtracted images. Because fluorescence intensity, morphology, and background varied among markers and imaging plates, segmentation parameters were optimized independently for each marker. A parameter grid search was performed using representative wells, and the resulting masks were visually compared with the corresponding grayscale fluorescence images (**Fig. S9B**). The parameter combination that most accurately captured marker-positive structures while limiting background segmentation and object merging was selected manually. The selected segmentation method and parameters for every marker and plate were recorded in a structured CSV file and applied to the remaining images from that plate.

Segmentation quality was assessed using proof images in which object boundaries or masks were overlaid in red on the corresponding grayscale fluorescence channel. Proof images were generated for all marker masks from representative wells, together with examples from the parameter grid searches (**Fig. S9B-C**). Additional quality control was performed by calculating pairwise overlap among marker masks (**Fig. S9D**). Markers associated with related intracellular compartments were expected to exhibit greater spatial overlap than markers representing spatially distinct compartments. Unexpected overlap patterns prompted reinspection of the underlying images and, where necessary, resegmentation using more conservative thresholds. Final segmentation parameters were selected before biological comparisons were performed.

### Punctum-level and cell-level quantification

Quantitative measurements were restricted to cells that passed segmentation, registration, and cross-round retention criteria. For each cell, the analysis pipeline quantified marker fluorescence intensity within each peptide punctum normalized by whole cell intensity and divided that measurement by the normalized marker fluorescence intensity within a radial 10-15 pixels away from the centroid of the peptide punctum mask.

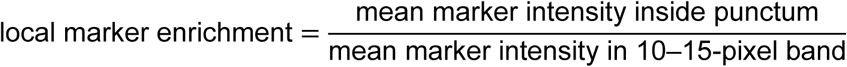

Measurements were retained at the individual-punctum level and linked to the corresponding cell, well, plate, peptide, and treatment time point.

GP30, a negative-control peptide lacking appreciable punctate uptake, was used to estimate nonspecific fluorescence, false positive peptide segmentation and autofluorescence. A GP30-derived intensity threshold was calculated separately for each imaging plate using mean punctum intensity for the false-positive puncta. Candidate peptide puncta with fluorescence intensities less than or equal to the corresponding GP30 threshold were excluded from all downstream punctum-based analyses. The same filtered punctum set was used consistently for marker-intensity, marker-distance, and marker-overlap measurements to prevent analysis-specific differences in punctum inclusion.

### Multichannel visualization

For qualitative visualization of intracellular trafficking, final marker masks from successive imaging rounds were assigned distinct pseudocolors and superimposed for a representative cell (**Fig. 3BC**). The displayed cell was selected from the quality-controlled population and was processed using the same registration, background correction, segmentation, and filtering procedures used for quantitative analysis. Image brightness and contrast were adjusted for visualization only and did not affect the quantitative measurements.

UMAP visualization of endosomal marker enrichment (**Fig. 4E-F**)

To characterize the intracellular trafficking profiles of peptide-containing puncta, individual puncta were represented by the enrichment of 14 subcellular markers: CI-M6PR, EEA1, GAL8, GM130, LAMP1, LAMP2, LC3B, mTOR, p62, RAB5, RAB7, RAB11, RAB14, and TGN46. For each punctum, marker enrichment was calculated as the mean marker fluorescence intensity within the peptide punctum divided by the mean marker fluorescence intensity across the corresponding cell. Enrichment values were log2-transformed and clipped to the range of −4 to 8 to limit the influence of extreme values.Only non-nuclear puncta were included. Markers were required to have valid measurements in at least 90% of puncta and a standard deviation of at least 0.15. Puncta missing a valid value for any retained marker were excluded from the analysis.

Marker features were standardized to zero mean and unit variance before principal component analysis (PCA). Only marker enrichment was included in the PCA, not peptide enrichment or intensity information. Principal components were retained in order of explained variance until at least 90% of the cumulative variance was captured (**Fig. S10A**). The retained PCA scores were then embedded in two dimensions using Uniform Manifold Approximation and Projection (UMAP) with 15 nearest neighbors, a minimum distance of 0.1, Euclidean distance, and a random seed of 42. Additional UMAP parameters were set to a spread of 1.0, learning rate of 1.0, repulsion strength of 1.0, and negative sampling rate of 5. Different nearest neighbors values were tested to ensure the relationships were robust.

The resulting UMAP coordinates were colored after embedding according to individual marker-enrichment values or peptide fluorescence intensity within each punctum based on enrichment distributions (**Fig. S10B-C**). These overlay measurements were used only for visualization and did not influence the UMAP embedding. All marker overlays (See **Fig. S10C**)

### Galectin-enrichment analysis in lysosomes (lamp1) and late endosomes (rab7)

To assess galectin recruitment to peptide-containing endolysosomal compartments, LAMP1and RAB7-defined objects were analyzed independently. Objects were obtained from the marker-object segmentation dataset and assigned to peptide conditions based on plate and well identity. Analyses were performed separately for CROT, GR30, and LL-37, with all sampled timepoints for each peptide pooled for this analysis. For each source-marker population, only objects enriched for the defining marker were retained. Specifically, LAMP1-derived objects were required to exhibit LAMP1 log2 enrichment >2, and RAB7-derived objects were required to exhibit RAB7 log2 enrichment >2. Marker enrichment was calculated as the mean fluorescence intensity within the segmented object relative to the mean whole-cell fluorescence intensity. Within each LAMP1or RAB7-enriched object, GAL3 and GAL8 enrichment were quantified using the same inside-object versus whole-cell normalization. Objects with GAL3 or GAL8 log2 enrichment >2 were classified as galectin-positive, whereas objects in which neither GAL3 nor GAL8 exceeded this threshold were classified as galectin-negative. Peptide accumulation was then compared between galectin-positive and galectin-negative compartments using the mean raw peptide fluorescence intensity within each source object. LAMP1and RAB7-defined compartments were analyzed separately for each peptide. For statistical comparison, peptide intensity was summarized at the punctum level and the resulting punctum-level distributions were compared using a two-sided Kolmogorov– Smirnov test. See **Fig. S8C** for more information.

### CRISPRi genetic screen

To perform the CRISPRi screen, we followed the protocol as described previously^88^ with modifications^87^. The screen was repeated in duplicate.

A previously designed 4 sgRNA/gene CRISPR-Cas9 library was used targeting 5′ ends of conserved exons with sgRNAs varying in length between 19 and 25 base-pairs^174^. The library was generated first by infecting K562 cells with a SFFV-Cas9-BFP vector to create a stably expressed Cas9 cell line. We then infected the lentiviral genome-wide sgRNA library into approximately 120 million cells following the same protocol as the genome-wide shRNA library to maintain at least 1,000-fold representation in cells. Infected cells were selected with puromycin (0.7 μg/mL, Sigma) for 3 days. The CRISPRi cell library was expanded and maintained in RPMI + 10% FBS in spinner flask at 60rpm. Cells were passaged at 1x10^6 cells/ml to ∼0.5x10^6 cells/ml daily, freezing down whole-genome replicates in 5-6ml freezing buffer.

Cells were split to ∼0.65x10^6 in 700ml of RPMI + 10% FBS in a new spinner flask (pre-warmed to 37°C overnight). We added Cy3-Crotamine to a final concentration of 125nM and incubated cells with rotation for 1hr. Note, we used a 125nM concentration as this gave us a good dynamic range to identify enhancers and suppressors (see **Fig. S4A**). The cell suspension was transferred to four 225ml tubes, and concentrated using centrifugation at 500g, 10 minutes, 4°C. Cell pellets were resuspended in 20ml ice cold PBS, pooled into two 50ml conical tubes (each containing 40ml). Cell suspensions were centrifuged as above, and each tube was resuspended in 8ml FACS buffer (PBS pH7.4, 25mM HEPES, 2% FBS, 2mM EDTA). Cell suspensions were filtered through a 30μm filter (CellTrics). Filters were washed with additional 1.5ml of FACS buffer. Strained cells were pooled into a 50ml conical tube. Two ml of the cell suspension was kept aside as an unsorted population (∼50x10^6 cells). The remaining cell suspension was transferred 5ml FACS tubes (4ml/tube) and kept on ice. Cells were sorted using a BD FACSAria II cell sorter with a 70 μm nozzle (BD Biosciences). Cell populations containing the lowest and highest 10% of Cy3-Crotamine levels— adjusted for cell size (FSC-W)—were sorted, respectively. Each sorted population, as well as the unsorted (starting) population, were spun down at 600g for 20 min at room temperature before extracting genomic DNA.

Genomic DNA was extracted immediately after pelleting using the Blood and Tissue DNeasy Maxi Kit (QIAGEN, 51194). The DNA was isolated according to the manufacturer’s instructions, except for eluting with buffer EB, rather than buffer AE. The samples were prepared for deep sequencing using two sequential PCR reactions to 1) amplify sgRNA sequences and 2) barcode each sample (for later pooling all samples for sequencing and to then computationally deconvolute)—as described previously^88^—using Agilent Herculase II Fusion DNA Polymerase Kit. The second PCR products were then run on a 2% TBE-agarose gel and gel purified using QIAquick Gel Extraction Kit (QIAGEN, 28706). The concentrations of the purified second PCR products were measured using a Qubit. After pooling all samples and diluting them further ac-cording to NextSeq 500/550 High Output Kit v2.5 (Illumina, FC-404-2001) instructions, deep se-quencing was performed on an Illumina NextSeq 550 platform to determine library composition. Guide composition between the sorted top 10% and the unsorted (starting) populations were compared using Cas9 high Throughput maximum Likelihood Estimator (casTLE)^88^ to determine genes that, when knocked down, increased or decreased Cy3-Crotamine uptake levels. Briefly, enrichment of individual guides was calculated as median normalized log ratios of counts between the various conditions. Gene-level effects were then calculated from ten guides targeting each gene, and an effect size estimate was derived for each gene with an associated-likelihood ratio to describe the significance of the gene-level effects. By randomly permutating the targeting ele-ments, the distribution of the log likelihood ratio was estimated and p values derived^88^.

### LDLR-eGFP plasmid transfection

The lipofectamine 3000 reagent kit (Invitrogen, L3000001) was used for all transfections based on manufacturer protocol. Briefly, opti-mem (Gibco, 31985070), 100 ng of DNA, p3000 and lipo3000 reagent were combined and added to cells in a cellvis glass 96 well plate and incubated for 16 hours before the experiment was performed. Treatment of +CPPs followed previous methods described and normal immunofluorescence (EEA1, Cell Signaling Technology, #3288) and click chemistry was performed. Cells were transfected with a custom pcDNA3.1(+)-based plasmid encoding human LDLR fused at its C terminus to eGFP (LDLR-pcDNA3.1(+)-C-eGFP; GenScript; internal plasmid ID pBoey0345/SB273) (See **Table S9**)

### Lipoprotein and EEA1 colocalization assays

DiI-labeled LDL, HDL, or VLDL (Kalen Biomedical, #770230-9, #770130-9, #770330-9) was added to cells together with the indicated peptide (500 nM) at a final lipoprotein concentration of 10 µg/mL. For all lipoprotein colocalization experiments, cells were seeded in glass-bottom plates without poly-D-lysine coating to minimize nonspecific interactions between negatively charged lipoprotein particles and the cationic polymer. Following cotreatment at 37 °C for the indicated duration, cells were fixed with 4% paraformaldehyde and limaged immediately to document the localization of DiI-labeled lipoproteins before permeabilization. Cells were subsequently permeabilized with 0.4% Triton X-100 in PBS for 5 min, and alkyne-modified peptides were fluorescently labeled using the click-chemistry procedure described above. The same fields were then reimaged using saved stage coordinates. Preand post-permeabilization images were compared to determine whether permeabilization altered lipoprotein localization. Although permeabilization reduced the overall DiI fluorescence intensity, it did not appreciably alter the spatial distribution of lipoprotein-positive puncta. Post-permeabilization images were therefore considered suitable for colocalization analysis. For EEA1 colocalization experiments, peptide-treated cells were processed using the immunofluorescence and click-labeling procedures described above, with EEA1 detected using the corresponding primary and secondary antibodies (EEA1, Cell Signaling Technology, #3288). Because the peptides differed in punctum morphology and fluorescence intensity, peptide-segmentation parameters were optimized separately for each peptide using a parameter grid search. Candidate segmentation masks were overlaid on the corresponding raw fluorescence images, and the parameter set that most accurately captured visually identifiable peptide puncta while limiting background segmentation was selected. GP30, a negative-control peptide, was used to establish the background fluorescence threshold. The mean intensity of GP30-associated puncta was calculated separately for each imaging plate, and candidate peptide puncta with intensities less than or equal to this threshold were excluded from downstream analyses because they could not be reliably distinguished from nonspecific fluorescence or imaging artifacts. EEA1and lipoprotein-positive structures were segmented using Otsu’s automatic thresholding method and all proofs were visually inspected.

### Microscale thermophoresis/ spectral shift

Cy5-labeled peptides were synthesized by Biosynth, and free Cy5 dye (Sigma-Aldrich, catalog no. 777323-1MG) was used as a control. Peptides and free Cy5 dye were prepared in 1× phosphate-buffered saline (PBS) and maintained at a final concentration of 20 nM in all experiments. Low-density lipoprotein (LDL) (Kalen Biomedical, #770200-1) concentrations were calculated using an estimated molecular mass of 2,700 kDa. Starting from a concentration of 1.85 µM, LDL was prepared as a 16-point twofold serial dilution and mixed with each Cy5-labeled peptide or free-dye control. Samples were loaded into premium capillaries (NanoTemper Technologies, #MOK025) and incubated for 5 minutes. Spectral-shift and microscale thermophoresis measurements were performed using a Monolith X instrument (NanoTemper Technologies) with the excitation power set to 100% and the MST power set to medium. Runs that exhibited significant aggregation or absorption to the capillary were excluded from analysis. Each titration was performed 3-4 times as independent observations. The precise method (i.e., 650nm absorption, 670nm absorption, or 670/650 ratio) used to determine the approximate Ec_50_ was chosen based on the highest R^2^ value of the fitted dose response curves (**See Table S8**)

### Zeta potential assay

Peptides and LDL mixtures were prepared at specified concentrations in aqueous buffer (10mM Tris-HCl, pH 7.5) and loaded into Malvern zetasizer nano series disposable folded capillary cells (Malvern Panalytical, #DTS1070). Zeta potential was measured by electrophoretic light scattering using a Zetasizer Nano ZS90 (Malvern Panalytical) at 25°C. Samples were equilibrated for 3 minutes prior to measurement. Each sample included five measurements with a minimum of 10 runs per measurement, and results are reported as the mean ± SD.

### MIC assays

EcM2.1 ΔtolC *Escherichia coli* (Addgene plasmid #64053) was cultured overnight in cation-adjusted Mueller–Hinton broth supplemented with 100 µg/mL ampicillin. Cultures were adjusted to an optical density at 600 nm (OD₆₀₀) of 0.3 and diluted 1:100 into assay mixtures containing the indicated combinations of peptide and LDL. Assays were prepared in deep-well 96-well LoBind plates (Eppendorf), sealed with Breathe-Easier™ film (Diversified Biotech, BERM-2000), and incubated for 24 h at 37°C with shaking at 900 rpm and 85% relative humidity. Following incubation, samples were transferred to clear, flat-bottom 96-well plates, and bacterial growth was quantified by measuring OD₆₀₀ using a BioTek Synergy H1 microplate reader.

### Cryogenic Electron Tomography

LDL–peptide–Nanogold association was examined by incubating the mixture for ∼15 min at 37°C before vitrification. Peptides were conjugated to 1.4 nm Mono-Alkyne-Nanogold using click chemistry (Nanoprobes, Inc.; Cat. No. 2027A-5X6NMOL). For each sample, 5 µL peptide–Nanogold clicked stock was mixed with 15 µL LDL, giving final concentrations of ∼1.14 µM peptide– Nanogold and 0.15 µg/µL LDL, equivalent to ∼300 nM LDL particles. This corresponds to an approximate 4:1 peptide–Nanogold molar ratio, or ∼4 nanogold-labeled peptides per LDL particle. Cryo-EM grids were prepared using R2/1 200-mesh grids (Quantifoil Micro Tools GmbH; Cat. No. X-102-Cu200). Prior to sample application, grids were glow discharged in negative mode at 15 mA and 0.40 mbar for 15 s. A volume of 4 µL of the LDL–peptide–Nanogold mixture was applied to each grid. Grids were blotted using a Vitrobot Mark IV with a blot time of 1.5–2s, followed by plunge freezing in liquid ethane. Frozen grids were transferred to liquid nitrogen and maintained under cryogenic conditions until imaging.

Cryo-ET data were collected on a ThermoFisher Glacios TEM operated at 200 kV on a Falcon IV direct detector. Tilt series were acquired at a magnification of 120,000×, with a pixel size of 1.24 Å/pixel. Images were collected with spot size 7 and an exposure time of 0.64 s per tilt image. The target defocus was −3.0 µm, with recorded defocus values ranging approximately from −2.8 to −3.7 µm. Dose symmetric tilt series were collected from −60° to +60° with 2° increments. Total exposure was approximately 110 e⁻/Å². Tomograms were reconstructed using EMAN2^175,176^ and distance measurements between LDL particles and nanogold-labeled peptides were quantified using the volume-rendering mode in UCSF ChimeraX^177^. Distance was measured from the center of nanogold particle to the monolayer of the LDL particle. LDL particles were randomly identified and nanogold particle distance within a 60nm cutoff were measured. 60 nm was chosen to include both interacting and non-interacting nearby nanogold particles, to analyze the full spatial distribution around LDL rather than only choosing very close particles.

### Gel shift assay

A 0.75% agarose gel was prepared using 20-well combs positioned in the middle and at the bottom of the gel. Peptides were tested at final concentrations of 9.25, 18.5, 37, and 74 µM. DiIlabeled LDL (LDL-DiI) and unlabeled LDL were each used at a final concentration of 100 µg/mL (approximately 37 nM). Peptide and LDL solutions were prepared at 2× their final concentrations, combined in a 96-well LoBind plate (Eppendorf, #951032107), and incubated for 5 min to allow complex formation. Following incubation, 5 µL of 6× loading dye containing 30% glycerol and bromophenol blue was added to each sample, and 25 µL of the resulting mixture was loaded per well. Samples containing unlabeled LDL were loaded into the upper wells, whereas samples containing LDL-DiI were loaded into the lower wells to compare the migration behavior of labeled and unlabeled LDL. The loading orientation was alternated between independent replicates to control for potential positional effects. Electrophoresis was performed at 4 °C for 90 min at 90 V, followed by 90 min at 120 V. LDL-DiI fluorescence was imaged using the Alexa Fluor 546 channel on a Bio-Rad ChemiDoc MP imaging system. The gel was subsequently stained with Sudan Black B (Sigma-Aldrich, #S0395) for 20 min with gentle agitation. The gel was destained twice for 10 min in a solution containing 160 mL acetic acid, 200 mL acetone, and 650 mL water, followed by overnight destaining at room temperature with gentle agitation. The gel was then imaged using the silver stain setting on the Bio-Rad imaging system.

### Generation of LDLRKO, LDLROE, LDLRWT HEPG2 cell lines

HepG2 LDLR knockout (LDLR-KO) and LDLR-KO cells stably expressing LDLR-EGFP were generously provided by Simon Pfisterer (University of Helsinki). The generation and validation of these cell lines have been described previously^178^.

### RNA sequencing

Total RNA was extracted from HEPG2 wildtype and HEPG2 LDLR KO cells grown in 15 cm dishes using the Qiagen RNeasy Mini Kit (QIAGEN, #74104) according to the manufacturer’s instructions. RNA concentration and purity were assessed using Aligent TapeStation High Sensitivity RNA ScreenTape and 4 biological replicates were analyzed per condition. Purified RNA was submitted to Novogene for RNA quality assessment, library preparation, and sequencing. RNA integrity was assessed before library construction. Bioinformatics analysis was performed by Novogene as described by their standard analysis.

### Statistical analysis and reproducibility

All data were analyzed using custom-built python scripts and statistical analyses were performed using scipy.stats packages. Scripts output SVG plot files that were either aesthetically edited on Adobe Illustrator or CSV’s produced from python scripts were then visualized via GraphPad Prism 9.3.1. If replotted in GraphPad Prism, the same statistical analyses were performed and test details are shown in figure legends. All experiments were performed a minimum of 3 times, but if resources allowed, power analysis was performed to determine the appropriate number of N. Statistical analyses were performed on the independent experiments unless stated otherwise.

### Code availability

All relevant custom python scripts can be found at https://github.com/sovannytaylor/cpp-manuscript-2026.

---

**Figure S1:**
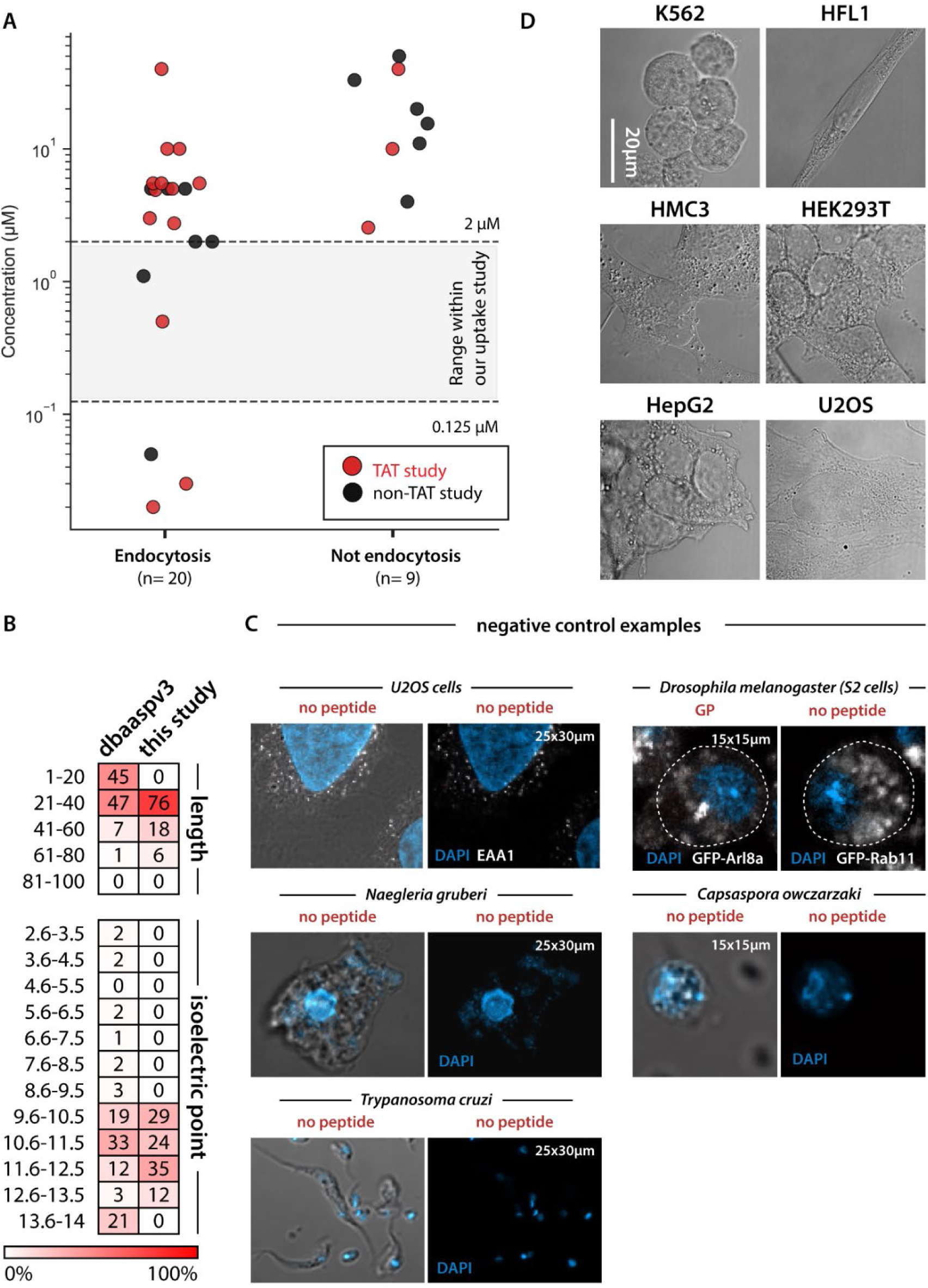
Characteristics of the peptide panel and representative control images. **(A)** Concentrations used in published uptake studies of cationic cell penetrant peptides. Around half of the studies investigated the HIV peptide, TAT. The shaded region indicates the concentration range used in this study (0.125–2 µM) (**See Table S1**) **(B)** Distribution of peptide length and isoelectric point in QBaasov3 and the present study. Values indicate the percentage of peptides within each bin. **(C)** No detected puncta is observed in negative-control images from the indicated cell types and organisms. **(D)** Immortalized cell types exhibit different morphological features.

**Figure S2:**
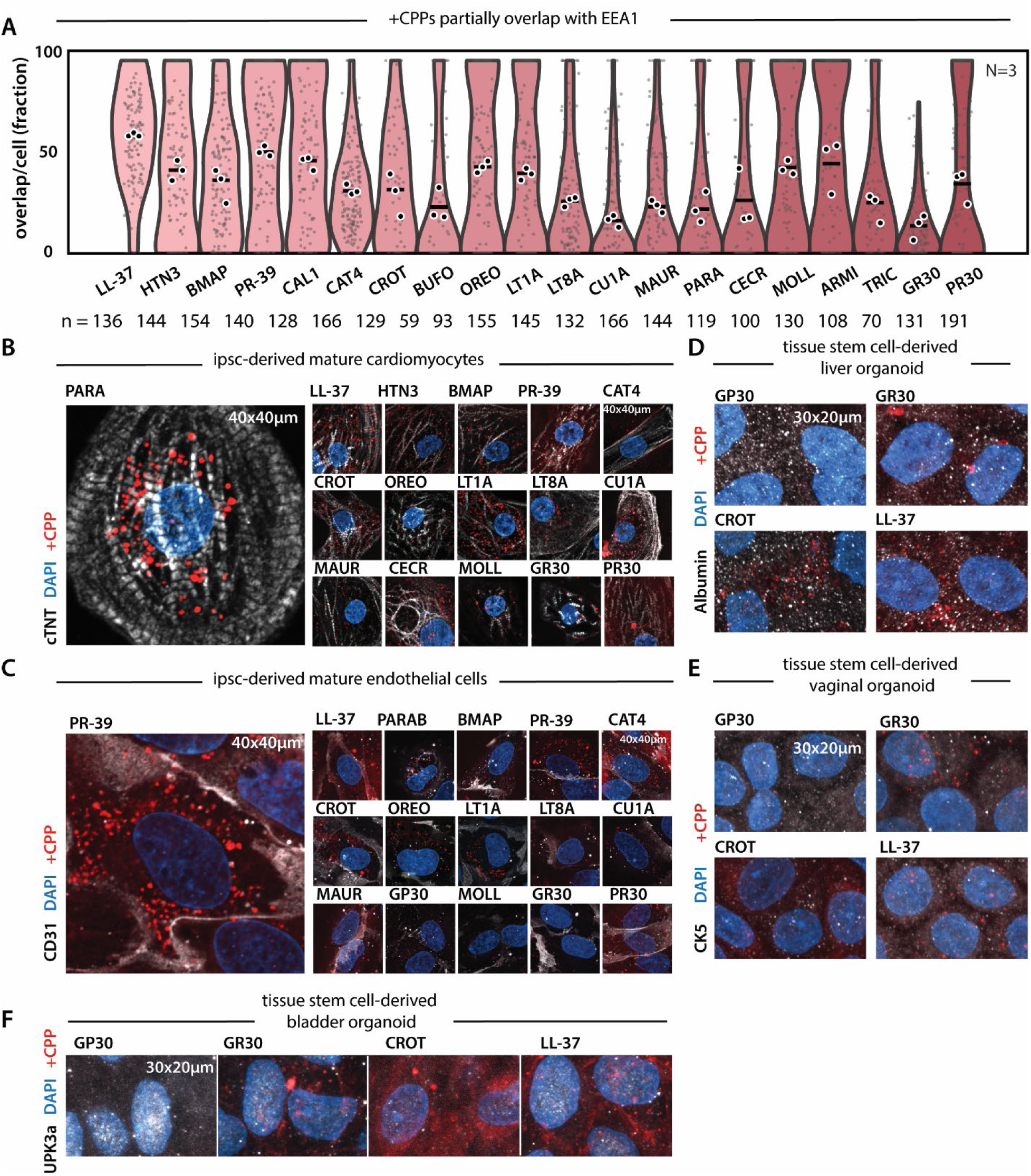
Entire +CPP library overlaps with early endosomal marker EEA1 and vesicle-like structures are observed in all tested cell types. **(A)** Phylogenetically diverse natural +CPPs localize to EEA1+ early endosomes in U2OS cells. Individual dots indicate the percentage of peptide+ puncta that overlap with EEA1 staining per cell. White outlined dots represent averages per independent experiment and black line indicates overall average. (see Fig. 1C**-D**). N = 3 replicates, n = 2,740 cells total. **(B)** +CPP puncta form vesicle-like structures in human iPSC-derived mature cardiomyocytes. cTNT marks cardiomyocytes. (1 µM, 60 min). Representative images. **(C)** +CPP puncta form vesicle-like structures in human iPSC-derived mature endothelial cells. CD31 marks the surface of endothelial cells. (1 µM, 60 min). Representative images. **(D)** +CPP puncta form vesicle-like structures in human tissue stem cell derived liver organoid monolayers. Albumin marks for hepatocytes. (1 µM, 60 min). Representative images. **(E)** +CPP puncta form vesicle-like structures in human tissue stem cell derived vaginal organoid monolayers. CK5 marks for basal and intermediate epithelial cells of the vagina. (1 µM, 60 min). Representative images. **(F)** +CPP puncta form vesicle-like structures in human tissue stem cell derived bladder organoid monolayers. UPK3a marks for urothelial cells. (1 µM, 60 min). Representative images.

**Figure S3:**
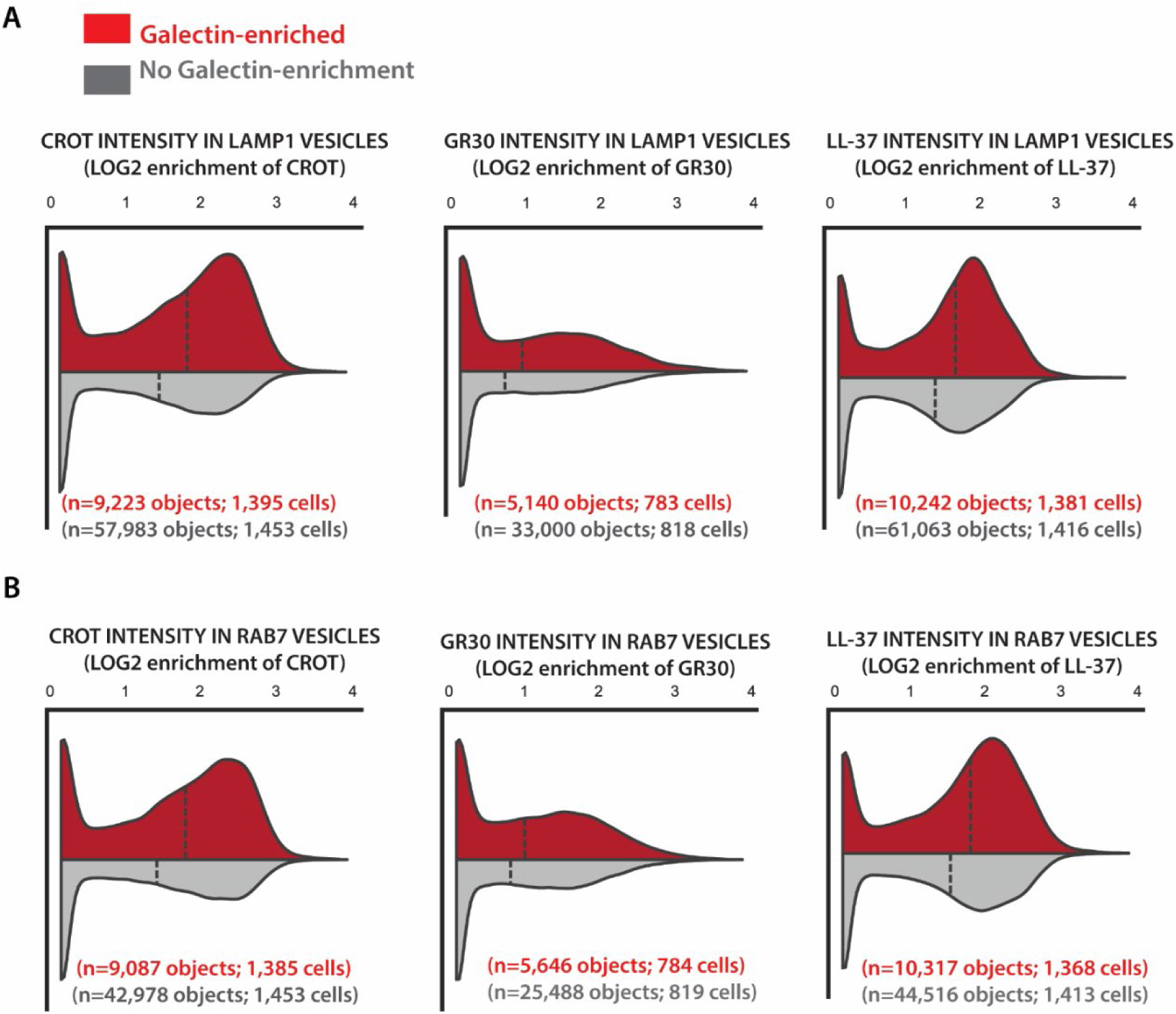
Galectin-enriched lysosomes (LAMP1⁺) and late endosomes (RAB7⁺) contain higher levels of +CPPs. **(A)** LAMP1+ lysosomes with detectable LGALS3/8 enrichment (red; n = 24,605 lysosomes) have on average a higher +CPP intensity than those without LGALS3/8 enrichment (gray; n = 152,046 lysosomes), indicative of lysosomal membrane damage and peptide escape (see also Fig. 3G). **(B)** RAB7+ endosomes with detectable LGALS3/8 enrichment (red; n = 25,050 RAB7+ endosomes) have on average a higher +CPP intensity than those without LGALS3/8 enrichment (gray; n = 112,982 RAB7+ endosomes), indicative of membrane damage and peptide escape (see also Fig. 3G).

**Figure S4:**
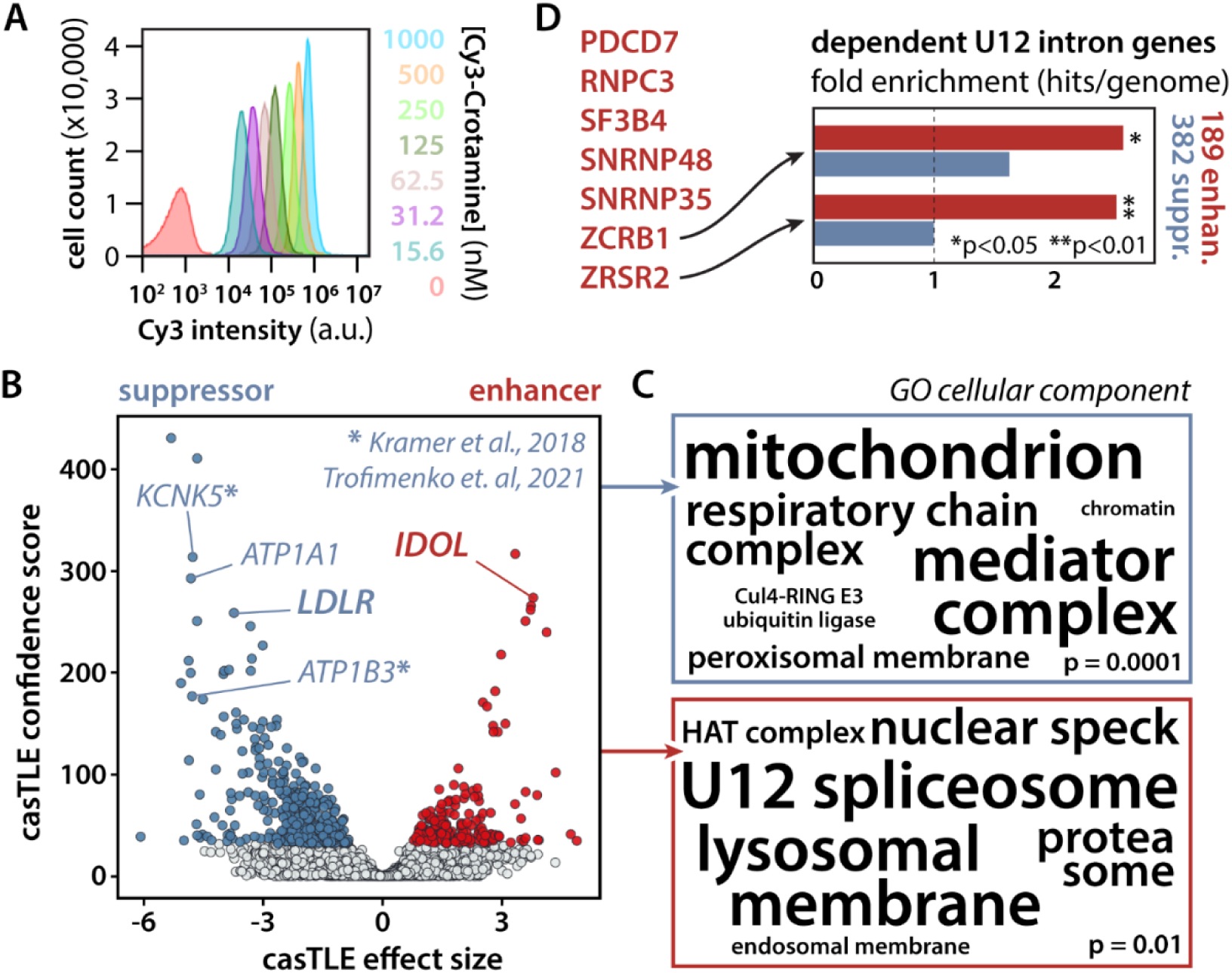
A genome-wide CRISPRi screen identifies regulators of +CPP uptake. **(A)** Dose-dependent uptake of Cy3-crotamine measured by flow cytometry. **(B)** casTLE analysis showing suppressors (blue) and enhancers (red) of uptake. Candidate genes found in other studies indicated by asterisk. (See **Table S3-4**) **(C)** Gene ontology enrichment of screen hits shows wide variety of pathways. (See **Supplemental Text B**) **(D)** Enrichment of U12 intron-containing genes are among enhancer and suppressor hits. *P < 0.05; **P < 0.01. (See **Supplemental Text B**)

**Figure S5:**
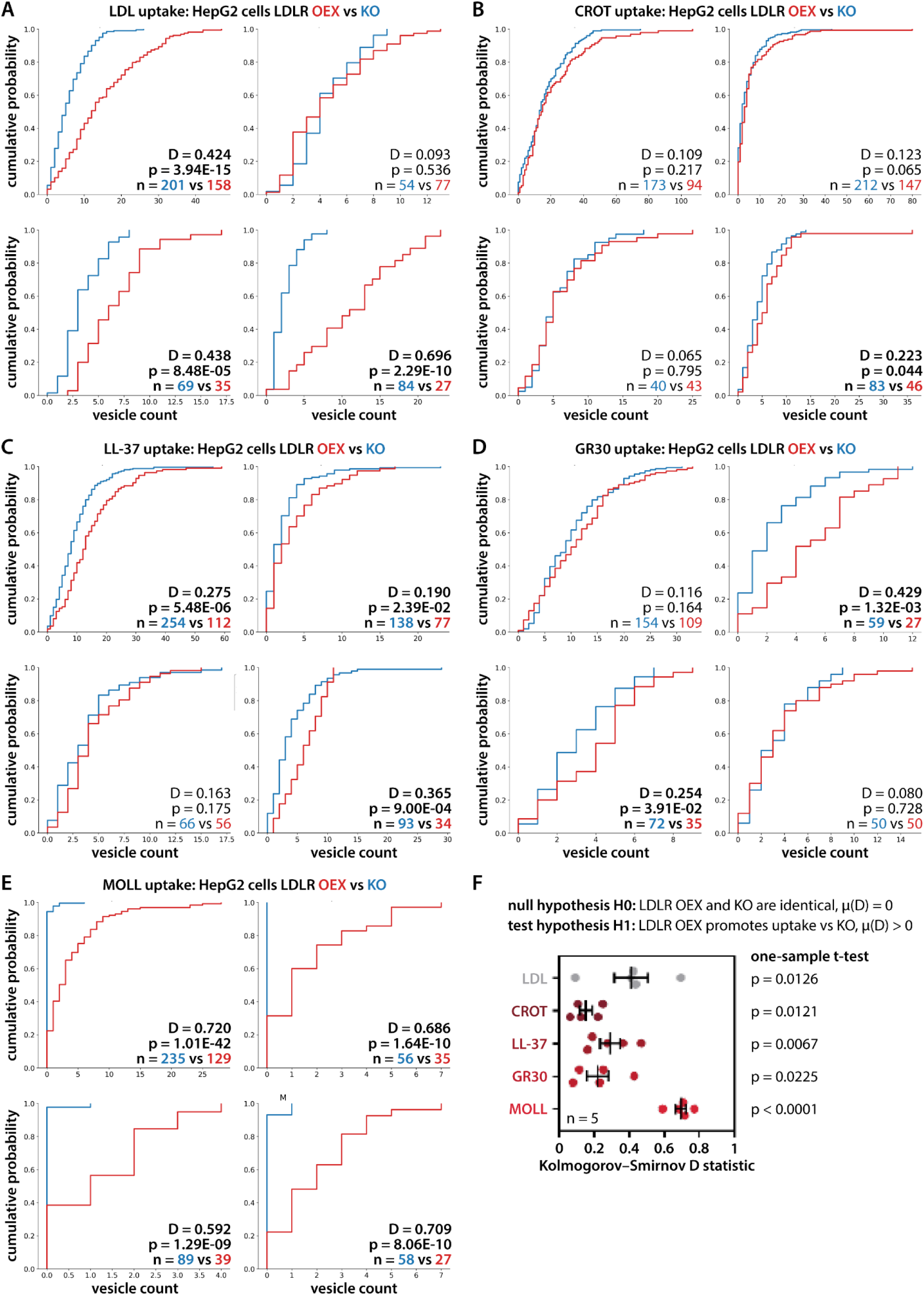
LDLR overexpression increases LDL and +CPP uptake in HEPG2 cells. **(A-E)** Cumulative distributions of LDL, CROT, LL-37, GR30, and MOLL-positive vesicle counts in LDLR-overexpressing and LDLR knockout cells show reduction of uptake in KO across independent experiments. Kolmogorov-Smirnov statistics, P values, and cell numbers are indicated. **(F)** KS D statistics across five independent experiments. A one-sample, one-sided t-test evaluated whether the mean directional D was greater than zero, indicating consistently increased uptake in LDLR-OEX relative to LDLR-KO cells. Points represent individual experiments; black bars show mean ± SEM. (See also Fig. 4G)

**Figure S6:**
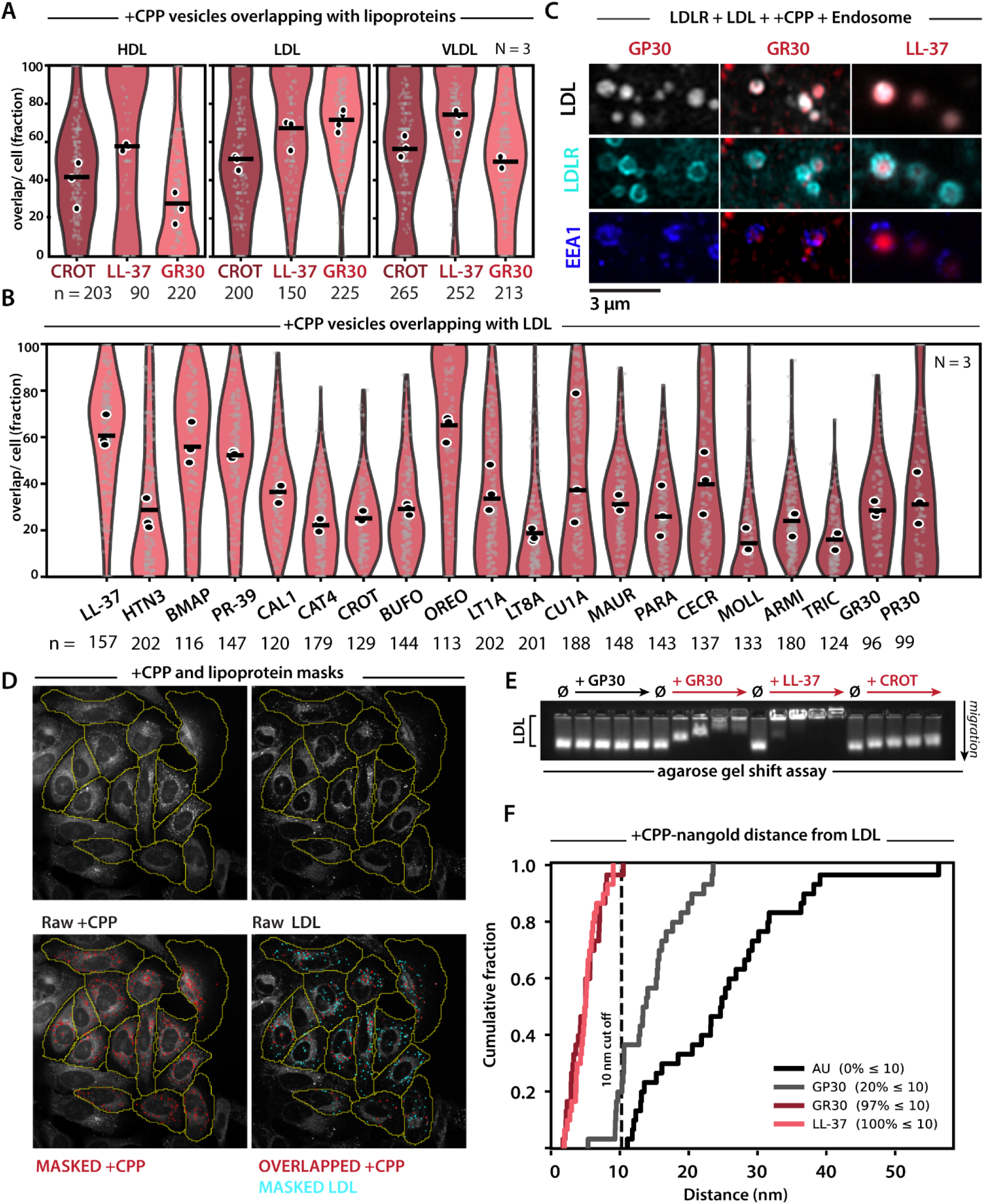
+CPPs associate with extracellular lipoproteins and remain associated during cellular uptake. **(A)** Fraction of CROT-, LL-37-, and GR30-positive vesicles overlapping with HDL, LDL, or VLDL. N = 3 replicates; n = 1,818 cells total. (See also Fig. 5C) **(B)** LDL overlap across a phylogenetically broad panel of +CPPs, demonstrating that lipoprotein association is a shared but peptide-dependent property in U2OS cells. Violin plots show cell-level distributions; white points represent replicate means. N = 3; n = 2,958 cells total. (See also Fig. 5E) **(C)** Representative images showing LDLR, LDL, +CPPs, and the early-endosomal marker EEA1 colocalize across main +CPPs. (See also Fig. 5D) **(D)** Representative raw images and masks used to quantify +CPP–LDL overlap. **(E)** Agarose gel-shift assay demonstrating direct, peptide-dependent interactions with LDL. **(F)** Cumulative nanogold-to-LDL distances, showing that GR30 and LL-37 localize within 10 nm of LDL more frequently than GP30 or the gold-only control. Dashed line indicates 10 nm, relative close distance based on length of peptide and nanogolds. n= 120 nanogold objects total.

**Figure S7:**
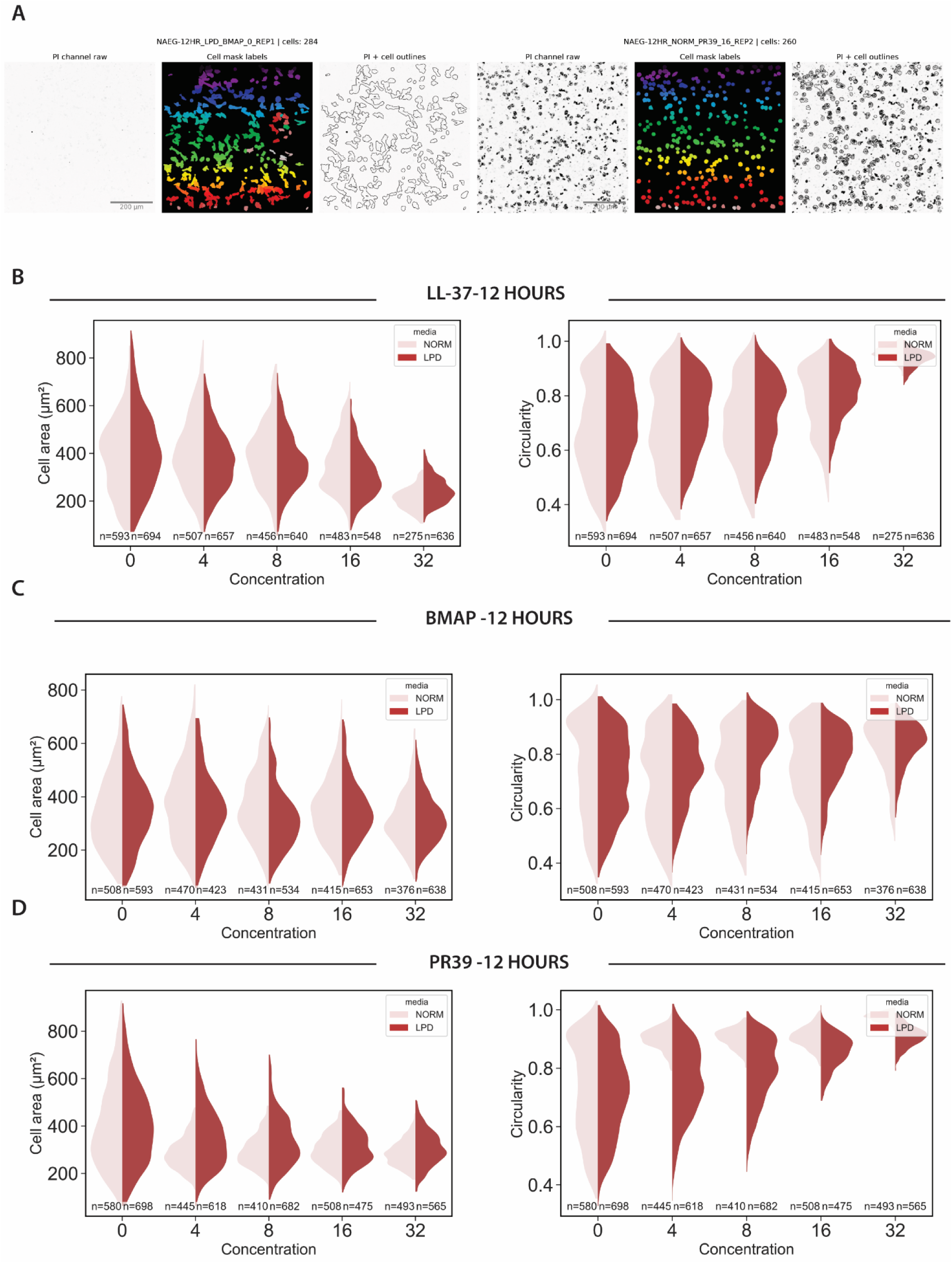
LL-37 and BMAP are not toxic to *Naegleria gruberi*. **(A)** Representative proof images of masks created for naegleria cell body and PI stain channel overlayed. **(B)** LL-37 does not show profound toxicity effects at the lower micromolar concentrations. Toxicity begins past 16 μM. **(C)** BMAP does not show profound toxicity effects at the lower micromolar concentrations. Toxicity is not observed at 32 μM. **(D)** PR39 induces toxicity-associated morphological changes at concentrations as low as 4 μM under normal media conditions, suggesting greater cytotoxic potency.

**Figure S8:**
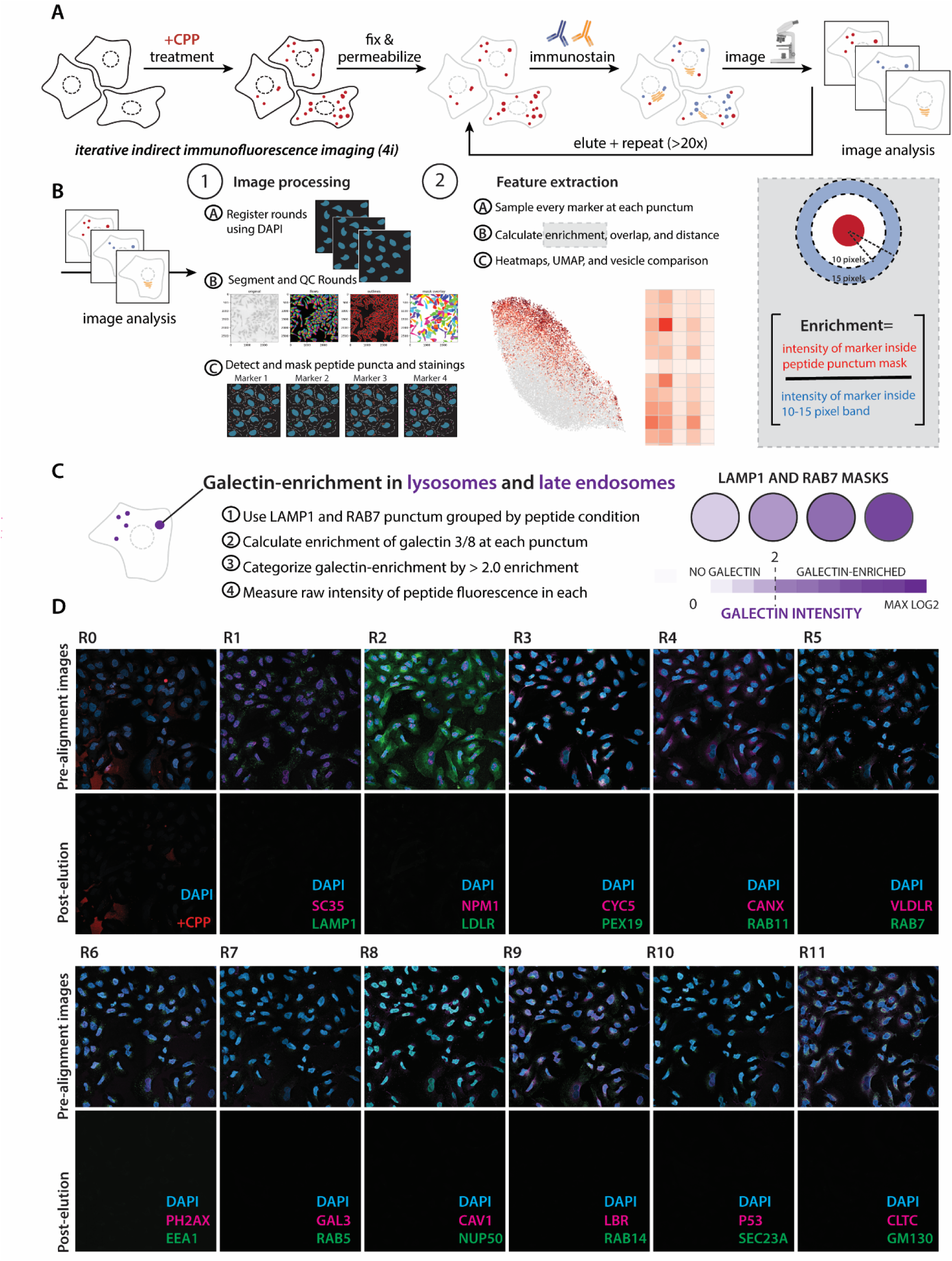
4i images and post-elution quality control. **(A)** Workflow of 4i (iterative, indirect, immunofluorescent imaging). **(B)** Workflow of 4i image analysis. **(C)** Workflow of galectin-enrichment analysis. **(D)** Representative images of pre-alignment images. Post-elution images were taken for 10 images per round. Representative. 10 out of 24 rounds shown. Scale dimensions indicated.

**Figure S9:**
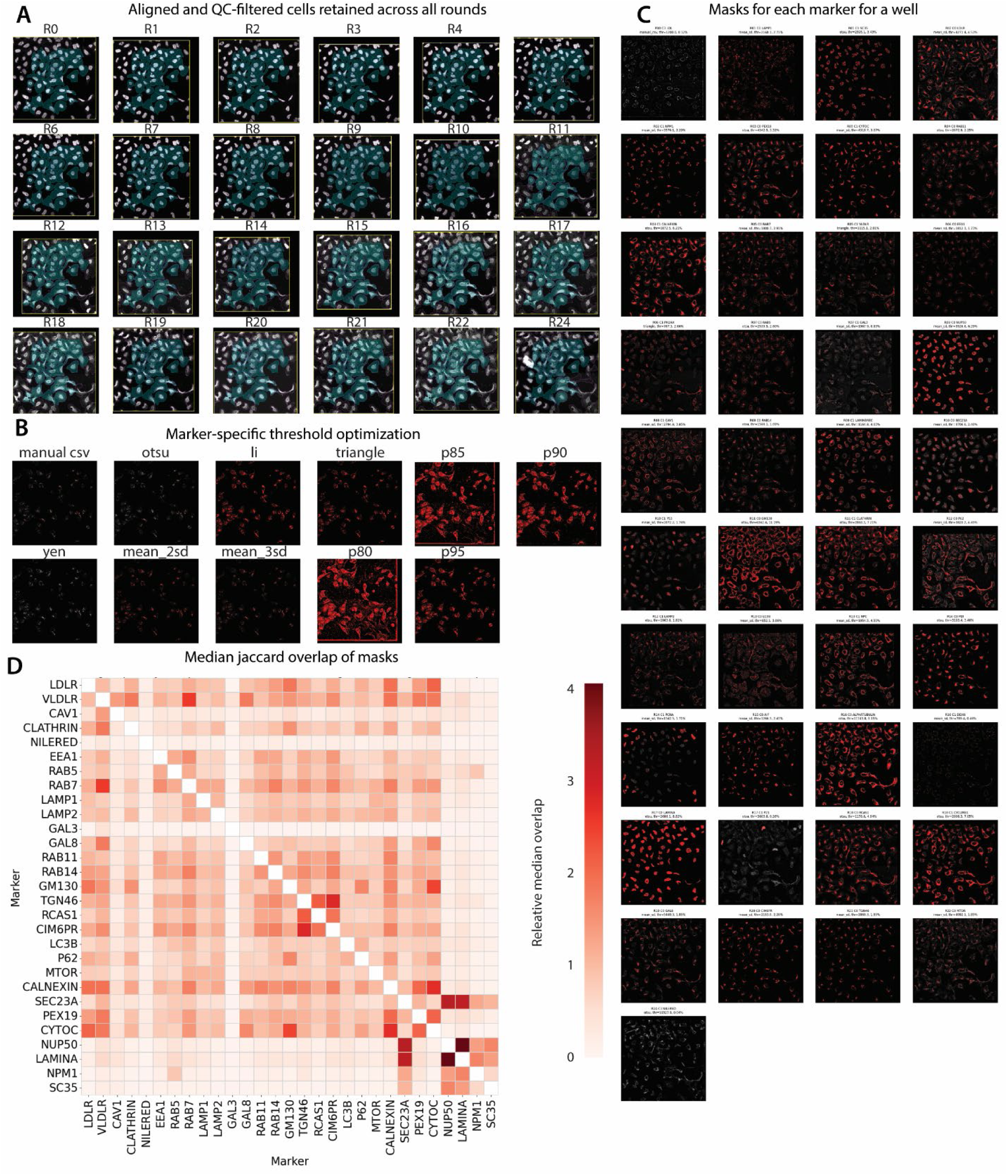
4i analysis quality-control parameters and workflow. **(A)** Images were aligned and cells were retained if present in all rounds and not touching the 10 pixel border in any round (cyan = cells kept, yellow = 10 pixel border region of interest). Representative well. **(B)** Grid search was performed to determine best mask segmentation method for each marker. Representative marker image. **(C)** Marker mask proof. Representative well. **(D)** Pixel-level marker mask overlap based on median jaccard grouped my compartment and normalized to highest off-diagonal overlap.

**Figure S10:**
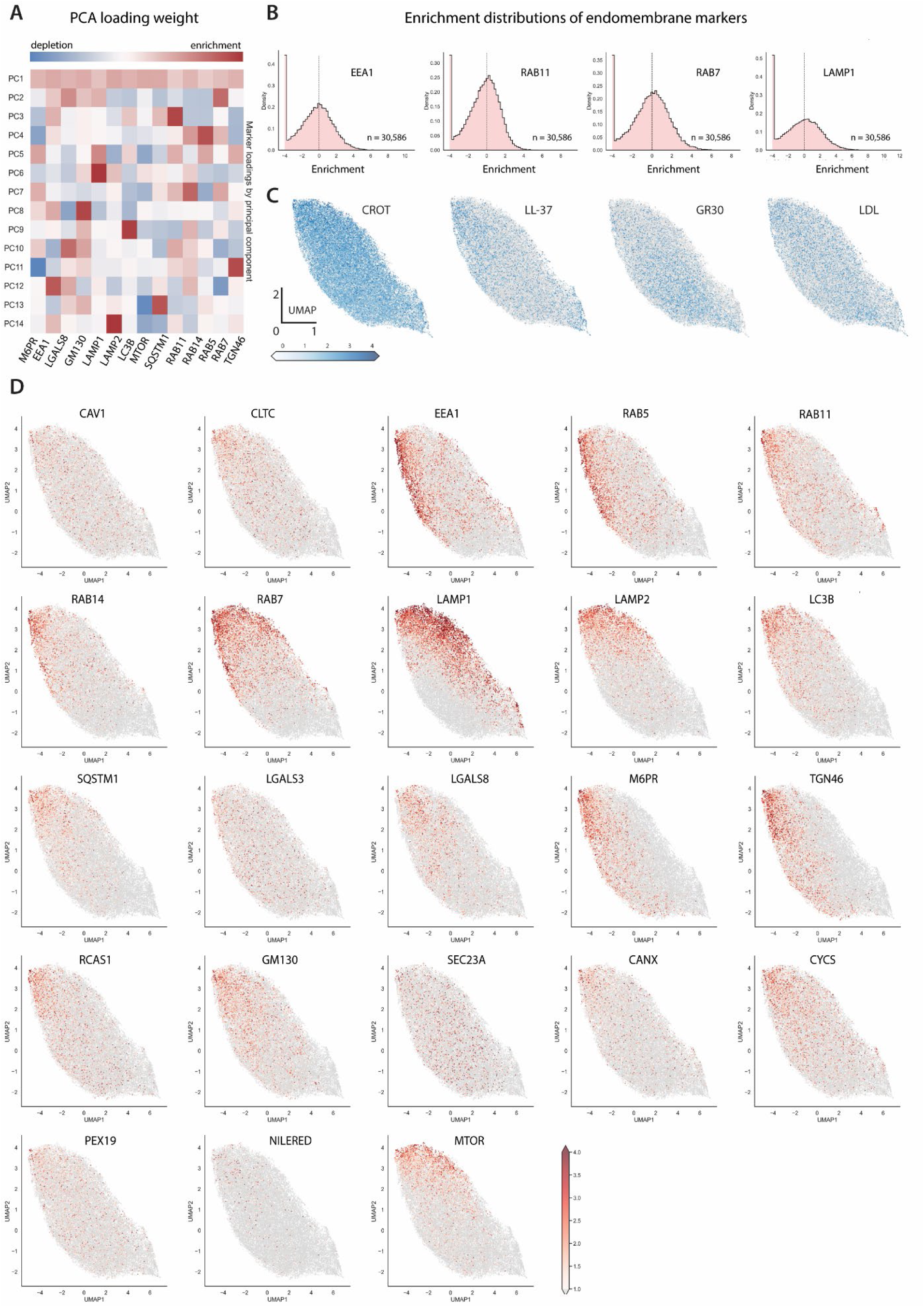
Dimensionality reduction of endomembrane-marker enrichment at individual peptide-containing vesicles. **(A)** Marker loadings for principal components PC1–PC14. Positive and negative values indicate relative marker enrichment and depletion, respectively. **(B)** Vesicle-level marker enrichment distributions. The dashed line indicates no enrichment (log₂ enrichment = 0). Distributions used to inform panel D overlay. **(C)** UMAP of cytoplasmic vesicles from GR30-, LL-37-, CROT-, and LDL-treated cells, colored by enrichment of peptide. LDL and +CPPs are indistinguishable in their endomembrane route. **(D)** UMAP of 30,586 cytoplasmic vesicles from GR30-, LL-37-, and CROT-treated cells, colored by enrichment of the indicated marker; gray points represent all other vesicles.

## REFERENCES

1. Milletti, F. Cell-penetrating peptides: classes, origin, and current landscape. Drug Discov. Today 17, 850–860 (2012).

2. Gori, A., Lodigiani, G., Colombarolli, S. G., Bergamaschi, G. & Vitali, A. Cell penetrating peptides: Classification, mechanisms, methods of study, and applications. ChemMedChem 18, e202300236 (2023).

3. Pooga, M. & Langel, Ü. Classes of cell-penetrating peptides. Methods Mol. Biol. 1324, 3–28 (2015).

4. Boeynaems, S., et al. Aberrant phase separation is a common killing strategy of positively charged peptides in biology and human disease. bioRxiv (2023) doi:10.1101/2023.03.09.531820.

5. Boeynaems, S. et al. Spontaneous driving forces give rise to protein-RNA condensates with coexisting phases and complex material properties. Proc. Natl. Acad. Sci. U. S. A. 116, 7889– 7898 (2019).

6. Guo, Z., Peng, H., Kang, J. & Sun, D. Cell-penetrating peptides: Possible transduction mechanisms and therapeutic applications. Biomed. Rep. 4, 528–534 (2016).

7. Zorko, M. & Langel, Ü. Cell-penetrating peptides. Methods Mol. Biol. 2383, 3–32 (2022).

8. Säälik, P. et al. Protein cargo delivery properties of cell-penetrating peptides. A comparative study. Bioconjug. Chem. 15, 1246–1253 (2004).

9. Guidotti, G., Brambilla, L. & Rossi, D. Cell-penetrating peptides: From basic research to clinics. Trends Pharmacol. Sci. 38, 406–424 (2017).

10. Frankel, A. D. & Pabo, C. O. Cellular uptake of the tat protein from human immunodeficiency virus. Cell 55, 1189–1193 (1988).

11. Green, M. & Loewenstein, P. M. Autonomous functional domains of chemically synthesized human immunodeficiency virus tat trans-activator protein. Cell 55, 1179–1188 (1988).

12. Derossi, D., Chassaing, G. & Prochiantz, A. Trojan peptides: the penetratin system for intracellular delivery. Trends Cell Biol. 8, 84–87 (1998).

13. Vivès, E., Brodin, P. & Lebleu, B. A truncated HIV-1 Tat protein basic domain rapidly translocates through the plasma membrane and accumulates in the cell nucleus. J. Biol. Chem. 272, 16010–16017 (1997).

14. Joliot, A., Pernelle, C., Deagostini-Bazin, H. & Prochiantz, A. Antennapedia homeobox peptide regulates neural morphogenesis. Proc. Natl. Acad. Sci. U. S. A. 88, 1864–1868 (1991).

15. Fretz, M. M. et al. Temperature-, concentration- and cholesterol-dependent translocation of Land D-octa-arginine across the plasma and nuclear membrane of CD34+ leukaemia cells. Biochem. J. 403, 335–342 (2007).

16. Wallbrecher, R. et al. Membrane permeation of arginine-rich cell-penetrating peptides independent of transmembrane potential as a function of lipid composition and membrane fluidity. J. Control. Release 256, 68–78 (2017).

17. Sun, D., Forsman, J., Lund, M. & Woodward, C. E. Effect of arginine-rich cell penetrating peptides on membrane pore formation and life-times: a molecular simulation study. Phys. Chem. Chem. Phys. 16, 20785–20795 (2014).

18. Herce, H. D. et al. Arginine-rich peptides destabilize the plasma membrane, consistent with a pore formation translocation mechanism of cell-penetrating peptides. Biophys. J. 97, 1917– 1925 (2009).

19. Herce, H. D. & Garcia, A. E. Molecular dynamics simulations suggest a mechanism for translocation of the HIV-1 TAT peptide across lipid membranes. Proc. Natl. Acad. Sci. U. S. A. 104, 20805–20810 (2007).

20. Jones, A. T. & Sayers, E. J. Cell entry of cell penetrating peptides: tales of tails wagging dogs. J. Control. Release 161, 582–591 (2012).

21. Trofimenko, E. et al. Genetic, cellular, and structural characterization of the membrane potential-dependent cell-penetrating peptide translocation pore. Elife 10, e69832 (2021).

22. Richard, J. P. et al. Cell-penetrating peptides. A reevaluation of the mechanism of cellular uptake: A reevaluation of the mechanism of cellular uptake. J. Biol. Chem. 278, 585–590 (2003).

23. Potocky, T. B., Menon, A. K. & Gellman, S. H. Cytoplasmic and nuclear delivery of a TAT-derived peptide and a beta-peptide after endocytic uptake into HeLa cells. J. Biol. Chem. 278, 50188–50194 (2003).

24. Mueller, J., Kretzschmar, I., Volkmer, R. & Boisguerin, P. Comparison of cellular uptake using 22 CPPs in 4 different cell lines. Bioconjug. Chem. 19, 2363–2374 (2008).

25. Lundberg, M., Wikström, S. & Johansson, M. Cell surface adherence and endocytosis of protein transduction domains. Mol. Ther. 8, 143–150 (2003).

26. Wadia, J. S., Stan, R. V. & Dowdy, S. F. Transducible TAT-HA fusogenic peptide enhances escape of TAT-fusion proteins after lipid raft macropinocytosis. Nat. Med. 10, 310–315 (2004).

27. Holmes, B. B. et al. Heparan sulfate proteoglycans mediate internalization and propagation of specific proteopathic seeds. Proc. Natl. Acad. Sci. U. S. A. 110, E3138–47 (2013).

28. Trofimenko, E., Homma, Y., Fukuda, M. & Widmann, C. The endocytic pathway taken by cationic substances requires Rab14 but not Rab5 and Rab7. Cell Rep. 37, 109945 (2021).

29. Wang, R. et al. Poly-PR in C9ORF72-Related Amyotrophic Lateral Sclerosis/Frontotemporal Dementia Causes Neurotoxicity by Clathrin-Dependent Endocytosis. Neurosci. Bull. 35, 889– 900 (2019).

30. Patel, S. G. et al. Cell-penetrating peptide sequence and modification dependent uptake and subcellular distribution of green florescent protein in different cell lines. Sci. Rep. 9, 6298 (2019).

31. Kramer, N. J. et al. CRISPR-Cas9 screens in human cells and primary neurons identify modifiers of C9ORF72 dipeptide-repeat-protein toxicity. Nat. Genet. 50, 603–612 (2018).

32. Tyagi, M., Rusnati, M., Presta, M. & Giacca, M. Internalization of HIV-1 tat requires cell surface heparan sulfate proteoglycans. J. Biol. Chem. 276, 3254–3261 (2001).

33. Wittrup, A., Zhang, S.-H. & Belting, M. Studies of proteoglycan involvement in CPP-mediated delivery. Methods Mol. Biol. 683, 99–115 (2011).

34. Perr, J. et al. RNA binding proteins and glycoRNAs form domains on the cell surface for cell penetrating peptide entry. bioRxiv 2023.09.04.556039 (2023) doi:10.1101/2023.09.04.556039.

35. Yu, P., Pearson, C. S. & Geller, H. M. Flexible roles for proteoglycan sulfation and receptor signaling. Trends Neurosci. 41, 47–61 (2018).

36. Hayashida, K., Aquino, R. S. & Park, P. W. Coreceptor functions of cell surface heparan sulfate proteoglycans. Am. J. Physiol. Cell Physiol. 322, C896–C912 (2022).

37. Park, P. W., Reizes, O. & Bernfield, M. Cell surface heparan sulfate proteoglycans: selective regulators of ligand-receptor encounters. J. Biol. Chem. 275, 29923–29926 (2000).

38. Pirtskhalava, M. et al. DBAASP v3: database of antimicrobial/cytotoxic activity and structure of peptides as a resource for development of new therapeutics. Nucleic Acids Res. 49, D288– D297 (2021).

39. Renton, A. E. et al. A hexanucleotide repeat expansion in C9ORF72 is the cause of chromosome 9p21-linked ALS-FTD. Neuron 72, 257–268 (2011).

40. DeJesus-Hernandez, M. et al. Expanded GGGGCC hexanucleotide repeat in noncoding region of C9ORF72 causes chromosome 9p-linked FTD and ALS. Neuron 72, 245–256 (2011).

41. Ash, P. E. A. et al. Unconventional translation of C9ORF72 GGGGCC expansion generates insoluble polypeptides specific to c9FTD/ALS. Neuron 77, 639–646 (2013).

42. Mori, K. et al. The C9orf72 GGGGCC repeat is translated into aggregating dipeptide-repeat proteins in FTLD/ALS. Science 339, 1335–1338 (2013).

43. Zu, T. et al. RAN proteins and RNA foci from antisense transcripts in C9ORF72 ALS and frontotemporal dementia. Proc. Natl. Acad. Sci. U. S. A. 110, E4968–77 (2013).

44. Kwon, I. et al. Poly-dipeptides encoded by the C9orf72 repeats bind nucleoli, impede RNA biogenesis, and kill cells. Science 345, 1139–1145 (2014).

45. Gittings, L. M. et al. Symmetric dimethylation of poly-GR correlates with disease duration in C9orf72 FTLD and ALS and reduces poly-GR phase separation and toxicity. Acta Neuropathol. 139, 407–410 (2020).

46. Westergard, T. et al. Cell-to-cell transmission of dipeptide repeat proteins linked to C9orf72ALS/FTD. Cell Rep. 17, 645–652 (2016).

47. Kerkis, I. et al. State of the art in the studies on crotamine, a cell penetrating peptide from South American rattlesnake. Biomed Res. Int. 2014, 675985 (2014).

48. Szewczyk-Roszczenko, O. K. et al. The chemical inhibitors of endocytosis: From mechanisms to potential clinical applications. Cells 12, 2312 (2023).

49. Dubovskii, P. V. et al. Latarcins: versatile spider venom peptides. Cell. Mol. Life Sci. 72, 4501– 4522 (2015).

50. Kuhn-Nentwig, L. et al. Cupiennin 1, a new family of highly basic antimicrobial peptides in the venom of the spider Cupiennius salei (Ctenidae). J. Biol. Chem. 277, 11208–11216 (2002).

51. Remijsen, Q., Verdonck, F. & Willems, J. Parabutoporin, a cationic amphipathic peptide from scorpion venom: much more than an antibiotic. Toxicon 55, 180–185 (2010).

52. Almaaytah, A., Tarazi, S., Mhaidat, N., Al-Balas, Q. & Mukattash, T. L. Mauriporin, a novel cationic α-helical peptide with selective cytotoxic activity against prostate cancer cell lines from the venom of the scorpion Androctonus mauritanicus. Int. J. Pept. Res. Ther. 19, 281– 293 (2013).

53. Kościuczuk, E. M. et al. Cathelicidins: family of antimicrobial peptides. A review. Mol. Biol. Rep. 39, 10957–10970 (2012).

54. Okello, I., Mafie, E., Eastwood, G., Nzalawahe, J. & Mboera, L. E. G. African animal trypanosomiasis: A systematic review on prevalence, risk factors and drug resistance in sub-Saharan Africa. J. Med. Entomol. 59, 1099–1143 (2022).

55. Traoré, B. M., et al. Free-ranging pigs identified as a multi-reservoir of Trypanosoma brucei and Trypanosoma congolense in the Vavoua area, a historical sleeping sickness focus of Côte d’Ivoire. PLoS Negl. Trop. Dis. 15, e0010036 (2021).

56. Chimelli, L. & Scaravilli, F. Trypanosomiasis. Brain Pathol. 7, 599–611 (1997).

57. Losinno, A. D., Martínez, S. J., Labriola, C. A., Carrillo, C. & Romano, P. S. Induction of autophagy increases the proteolytic activity of reservosomes during Trypanosoma cruzi metacyclogenesis. Autophagy 17, 439–456 (2021).

58. Campbell, P. C. & de Graffenried, C. L. Morphogenesis in Trypanosoma cruzi epimastigotes proceeds via a highly asymmetric cell division. PLoS Negl. Trop. Dis. 17, e0011731 (2023).

59. Alanazi, A. et al. Advancing the understanding of Naegleria fowleri: Global epidemiology, phylogenetic analysis, and strategies to combat a deadly pathogen. J. Infect. Public Health 18, 102690 (2025).

60. Daft, B. M. et al. Seasonal meningoencephalitis in Holstein cattle caused by Naegleria fowleri. J. Vet. Diagn. Invest. 17, 605–609 (2005).

61. Ithoi, I. et al. Morphological characteristics of developmental stages of Acanthamoeba and Naegleria species before and after staining by various techniques. Southeast Asian J. Trop. Med. Public Health 42, 1327–1338 (2011).

62. Ferrer-Bonet, M. & Ruiz-Trillo, I. Capsaspora owczarzaki. Curr. Biol. 27, R829–R830 (2017).

63. Seo, J.-K., Go, H.-J., Kim, C.-H., Nam, B.-H. & Park, N. G. Antimicrobial peptide, hdMolluscidin, purified from the gill of the abalone, Haliotis discus. Fish Shellfish Immunol. 52, 289– 297 (2016).

64. Ros-Rocher, N. et al. Chemical factors induce aggregative multicellularity in a close unicellular relative of animals. Proc. Natl. Acad. Sci. U. S. A. 120, e2216668120 (2023).

65. Kahlenberg, J. M. & Kaplan, M. J. Little peptide, big effects: the role of LL-37 in inflammation and autoimmune disease. J. Immunol. 191, 4895–4901 (2013).

66. Lande, R. et al. Native/citrullinated LL37-specific T-cells help autoantibody production in Systemic Lupus Erythematosus. Sci. Rep. 10, 5851 (2020).

67. Herster, F. et al. Neutrophil extracellular trap-associated RNA and LL37 enable self-amplifying inflammation in psoriasis. Nat. Commun. 11, 105 (2020).

68. Duan, Z. et al. Antimicrobial peptide LL-37 forms complex with bacterial DNA to facilitate blood translocation of bacterial DNA and aggravate ulcerative colitis. Sci. Bull. (Beijing) 63, 1364– 1375 (2018).

69. Altieri, A. et al. LL-37 and citrullinated-LL-37 modulate IL-17A/F-mediated responses and selectively suppress Lipocalin-2 in bronchial epithelial cells. J. Inflamm. (Lond.) 22, 20 (2025).

70. Chen, X. et al. Human antimicrobial peptide LL-37 contributes to Alzheimer’s disease progression. Mol. Psychiatry 27, 4790–4799 (2022).

71. Bhusal, A. et al. Cathelicidin-related antimicrobial peptide promotes neuroinflammation through astrocyte-microglia communication in experimental autoimmune encephalomyelitis. Glia 70, 1902–1926 (2022).

72. Stallaert, W. et al. The molecular architecture of cell cycle arrest. Mol. Syst. Biol. 18, e11087 (2022).

73. Stallaert, W. et al. The structure of the human cell cycle. Cell Syst. 13, 230–240.e3 (2022).

74. Hsu, J., Nguyen, K. T., Bujnowska, M., Janes, K. A. & Fallahi-Sichani, M. Protocol for iterative indirect immunofluorescence imaging in cultured cells, tissue sections, and metaphase chromosome spreads. STAR Protoc. 5, 103190 (2024).

75. Kramer, B. A., Del Castillo, J. S., Pelkmans, L. & Gut, G. Iterative Indirect Immunofluorescence Imaging (4i) on Adherent Cells and Tissue Sections. Bio Protoc 13, e4712 (2023).

76. Gut, G., Herrmann, M. D. & Pelkmans, L. Multiplexed protein maps link subcellular organization to cellular states. Science 361, (2018).

77. Du, Y. et al. Lipid turnover and GEF recruitment collectively determine Rab5 recruitment and activation during the first step of early endosome formation. Nat. Commun. 17, 6896 (2026).

78. Shearer, L. J. & Petersen, N. O. Distribution and Co-localization of endosome markers in cells. Heliyon 5, e02375 (2019).

79. Humphries, W. H., 4th, Szymanski, C. J. & Payne, C. K. Endo-lysosomal vesicles positive for Rab7 and LAMP1 are terminal vesicles for the transport of dextran. PLoS One 6, e26626 (2011).

80. Chahal, G. S., Helbig, K. J., Parton, R. G. & Monson, E. A. The biology of endosomal escape: Strategies for enhanced delivery of therapeutics. ACS Nano 20, 1789–1813 (2026).

81. Mehta, M. J., Kim, H. J., Lim, S. B., Naito, M. & Miyata, K. Recent progress in the endosomal escape mechanism and chemical structures of polycations for nucleic acid delivery. Macromol. Biosci. 24, e2300366 (2024).

82. Thurston, T. L. M., Wandel, M. P., von Muhlinen, N., Foeglein, A. & Randow, F. Galectin 8 targets damaged vesicles for autophagy to defend cells against bacterial invasion. Nature 482, 414–418 (2012).

83. Munson, M. J. et al. A high-throughput Galectin-9 imaging assay for quantifying nanoparticle uptake, endosomal escape and functional RNA delivery. *Commun*. Biol. 4, 211 (2021).

84. Rui, Y. et al. High-throughput and high-content bioassay enables tuning of polyester nanoparticles for cellular uptake, endosomal escape, and systemic in vivo delivery of mRNA. Sci. Adv. 8, eabk2855 (2022).

85. Troncoso, M. F. et al. The universe of galectin-binding partners and their functions in health and disease. J. Biol. Chem. 299, 105400 (2023).

86. Jia, W. et al. Indispensable role of Galectin-3 in promoting quiescence of hematopoietic stem cells. Nat. Commun. 12, 2118 (2021).

87. Kim, G. et al. Genome-wide CRISPR screen reveals v-ATPase as a drug target to lower levels of ALS protein ataxin-2. Cell Rep. 41, 111508 (2022).

88. Morgens, D. W., Deans, R. M., Li, A. & Bassik, M. C. Systematic comparison of CRISPR/Cas9 and RNAi screens for essential genes. Nat. Biotechnol. 34, 634–636 (2016).

89. Hamilton, M. C. et al. Systematic elucidation of genetic mechanisms underlying cholesterol uptake. Cell Genom. 3, 100304 (2023).

90. Berndsen, Z. T. & Cassidy, C. K. The structure of apolipoprotein B100 from human low-density lipoprotein. Nature 638, 836–843 (2025).

91. Ren, G. et al. Model of human low-density lipoprotein and bound receptor based on cryoEM. Proc. Natl. Acad. Sci. U. S. A. 107, 1059–1064 (2010).

92. Mehta, A. & Shapiro, M. D. Apolipoproteins in vascular biology and atherosclerotic disease. Nat. Rev. Cardiol. 19, 168–179 (2022).

93. Byun, J. H. & Daskalopoulou, S. S. Lipoprotein(a) in coronary artery disease. Nat. Rev. Cardiol. 22, 702 (2025).

94. Huang, C. et al. Proteomic and functional analysis of HDL subclasses in humans and rats: a proof-of-concept study. Lipids Health Dis. 22, 86 (2023).

95. Dashty, M. et al. Proteome of human plasma very low-density lipoprotein and low-density lipoprotein exhibits a link with coagulation and lipid metabolism. Thromb. Haemost. 111, 518– 530 (2014).

96. Bancells, C. et al. Proteomic analysis of electronegative low-density lipoprotein. J. Lipid Res. 51, 3508–3515 (2010).

97. Masuda, D. et al. Proteomic analysis of human chylomicron remnants isolated by apolipoprotein B-48 immunoprecipitation. J. Atheroscler. Thromb. 32, 226–238 (2025).

98. Fang, Y. et al. LL-37-ApoB-100 complex serves as a biomarker of coronary artery disease. Arterioscler. Thromb. Vasc. Biol. 46, e323486 (2026).

99. Ciornei, C. D., Sigurdardóttir, T., Schmidtchen, A. & Bodelsson, M. Antimicrobial and chemoattractant activity, lipopolysaccharide neutralization, cytotoxicity, and inhibition by serum of analogs of human cathelicidin LL-37. Antimicrob. Agents Chemother. 49, 2845–2850 (2005).

100. Post, S., Rueschpler, L. & Schloer, S. Endolysosomes as a sorting hub for emerging viruses: Gatekeepers of cellular defense, viral fate and promising therapeutic target. Pharmacol. Res. 222, 108051 (2025).

101. Victoria, G. S. & Zurzolo, C. The spread of prion-like proteins by lysosomes and tunneling nanotubes: Implications for neurodegenerative diseases. J. Cell Biol. 216, 2633–2644 (2017).

102. Miller, C. M., Wan, W. B., Seth, P. P. & Harris, E. N. Endosomal escape of Antisense oligonucleotides internalized by Stabilin receptors is regulated by Rab5C and EEA1 during endosomal maturation. Nucleic Acid Ther. 28, 86–96 (2018).

103. Du Rietz, H., Hedlund, H., Wilhelmson, S., Nordenfelt, P. & Wittrup, A. Imaging small molecule-induced endosomal escape of siRNA. Nat. Commun. 11, 1809 (2020).

104. Chatterjee, S., Kon, E., Sharma, P. & Peer, D. Endosomal escape: A bottleneck for LNP-mediated therapeutics. Proc. Natl. Acad. Sci. U. S. A. 121, e2307800120 (2024).

105. Lechardeur, D., Verkman, A. S. & Lukacs, G. L. Intracellular routing of plasmid DNA during non-viral gene transfer. Adv. Drug Deliv. Rev. 57, 755–767 (2005).

106. Clausen, T. M. et al. SARS-CoV-2 infection depends on cellular heparan sulfate and ACE2. Cell 183, 1043–1057.e15 (2020).

107. Tang, W.-H. et al. The lipid components of high-density lipoproteins (HDL) are essential for the binding and transportation of antimicrobial peptides in human serum. Sci. Rep. 12, 2576 (2022).

108. Nakamura, Y. et al. Increased LL37 in psoriasis and other inflammatory disorders promotes LDL uptake and atherosclerosis. J. Clin. Invest. 134, e172578 (2024).

109. Fang, Y. et al. Cathelicidin LL-37-ApoB-100 interaction promotes LDL clearance and attenuates cholesterol accumulation in the liver. Sci. China Life Sci. 69, 492–505 (2026).

110. Tsao, C. W. et al. Heart disease and stroke statistics-2023 update: A report from the American Heart Association. Circulation 147, e93–e621 (2023).

111. Pham, H. N. et al. Burden of hyperlipidemia, cardiovascular mortality, and COVID-19: A retrospective-cohort analysis of US data. J. Am. Heart Assoc. 14, e037381 (2025).

112. Zheutlin, A. R., Harris, B. R. E. & Stulberg, E. L. Hyperlipidemia-attributed deaths in the U.s. in 2018-2021. Am. J. Prev. Med. 66, 1075–1077 (2024).

113. Babokhov, P., Sanyaolu, A. O., Oyibo, W. A., Fagbenro-Beyioku, A. F. & Iriemenam, N. C. A current analysis of chemotherapy strategies for the treatment of human African trypanosomiasis. Pathog. Glob. Health 107, 242–252 (2013).

114. Güémez, A. & García, E. Primary Amoebic Meningoencephalitis by Naegleria fowleri: Pathogenesis and Treatments. Biomolecules 11, 1320 (2021).

115. Raulin, A.-C., Martens, Y. A. & Bu, G. Lipoproteins in the central nervous system: From biology to pathobiology. Annu. Rev. Biochem. 91, 731–759 (2022).

116. Lane-Donovan, C., Philips, G. T. & Herz, J. More than cholesterol transporters: lipoprotein receptors in CNS function and neurodegeneration. Neuron 83, 771–787 (2014).

117. Jackson, R. J., Hyman, B. T. & Serrano-Pozo, A. Multifaceted roles of APOE in Alzheimer disease. Nat. Rev. Neurol. 20, 457–474 (2024).

118. Lee, S. et al. Downregulation of Hsp90 and the antimicrobial peptide Mtk suppresses poly(GR)-induced neurotoxicity in C9ORF72-ALS/FTD. Neuron 111, 1381–1390.e6 (2023).

119. Lee, S., Silverman, N. & Gao, F.-B. Emerging roles of antimicrobial peptides in innate immunity, neuronal function, and neurodegeneration. Trends Neurosci. 47, 949–961 (2024).

120. Stuart, B. A. R., Franitza, A. L. & E, L. Regulatory roles of antimicrobial peptides in the nervous system: Implications for neuronal aging. Front. Cell. Neurosci. 16, 843790 (2022).

121. Rauch, J. N. et al. LRP1 is a master regulator of tau uptake and spread. Nature 580, 381– 385 (2020).

122. Ranganathan, S. et al. LRAD3, a novel low-density lipoprotein receptor family member that modulates amyloid precursor protein trafficking. J. Neurosci. 31, 10836–10846 (2011).

123. Boni-Mitake, M., Costa, H., Vassilieff, V. S. & Rogero, J. R. Distribution of (125)I-labeled crotamine in mice tissues. Toxicon 48, 550–555 (2006).

124. Nascimento, F. D. et al. The natural cell-penetrating peptide crotamine targets tumor tissue in vivo and triggers a lethal calcium-dependent pathway in cultured cells. Mol. Pharm. 9, 211– 221 (2012).

125. Marinovic, M. P. et al. Crotamine induces browning of adipose tissue and increases energy expenditure in mice. Sci. Rep. 8, (2018).

126. Lima, S. de C., et al. Pharmacological characterization of crotamine effects on mice hind limb paralysis employing both ex vivo and in vivo assays: Insights into the involvement of voltage-gated ion channels in the crotamine action on skeletal muscles. PLoS Negl. Trop. Dis. 12, e0006700 (2018).

127. Bdolah, A. The venom glands of snakes and venom secretion. in Snake Venoms 41–57 (Springer Berlin Heidelberg, Berlin, Heidelberg, 1979).

128. Chen, C. I.-U., Schaller-Bals, S., Paul, K. P., Wahn, U. & Bals, R. Beta-defensins and LL37 in bronchoalveolar lavage fluid of patients with cystic fibrosis. J. Cyst. Fibros. 3, 45–50 (2004).

129. Scott, A. et al. Evaluation of the ability of LL-37 to neutralise LPS in vitro and ex vivo. PLoS One 6, e26525 (2011).

130. Keutmann, M. et al. The ratio of serum LL-37 levels to blood leucocyte count correlates with COVID-19 severity. Sci. Rep. 12, 9447 (2022).

131. Gambichler, T. et al. Serum levels of antimicrobial peptides and proteins do not correlate with psoriasis severity and are increased after treatment with fumaric acid esters. Arch. Derm. Res. 304, 471–474 (2012).

132. Davidopoulou, S. et al. Salivary concentration of the antimicrobial peptide LL-37 in patients with oral lichen planus. J. Oral Microbiol. 6, 26156 (2014).

133. Ganguly, D. et al. Self-RNA-antimicrobial peptide complexes activate human dendritic cells through TLR7 and TLR8. J. Exp. Med. 206, 1983–1994 (2009).

134. Choi, S. Y. et al. C9ORF72-ALS/FTD-associated poly(GR) binds Atp5a1 and compromises mitochondrial function in vivo. Nat. Neurosci. 22, 851–862 (2019).

135. Sakae, N. et al. Poly-GR dipeptide repeat polymers correlate with neurodegeneration and Clinicopathological subtypes in C9ORF72-related brain disease. Acta Neuropathol. Commun. 6, 63 (2018).

136. Krishnan, G. et al. Poly(GR) and poly(GA) in cerebrospinal fluid as potential biomarkers for C9ORF72-ALS/FTD. Nat. Commun. 13, 2799 (2022).

137. Chu, B.-B. et al. Cholesterol transport through lysosome-peroxisome membrane contacts. Cell 161, 291–306 (2015).

138. Kovacs, W. J. et al. Disturbed cholesterol homeostasis in a peroxisome-deficient PEX2 knockout mouse model. Mol. Cell. Biol. 24, 1–13 (2004).

139. Subtil, A. et al. Acute cholesterol depletion inhibits clathrin-coated pit budding. Proc. Natl. Acad. Sci. U. S. A. 96, 6775–6780 (1999).

140. Moyer, D. C., Larue, G. E., Hershberger, C. E., Roy, S. W. & Padgett, R. A. Comprehensive database and evolutionary dynamics of U12-type introns. Nucleic Acids Res. 48, 7066–7078 (2020).

141. Powell-Rodgers, G., et al. Role of U11/U12 minor spliceosome gene ZCRB1 in Ciliogenesis and WNT Signaling. bioRxivorg (2024) doi:10.1101/2024.08.09.607392.

142. Madan, V. et al. Aberrant splicing of U12-type introns is the hallmark of ZRSR2 mutant myelodysplastic syndrome. Nat. Commun. 6, 6042 (2015).

143. Ricoult, S. J. H. & Manning, B. D. The multifaceted role of mTORC1 in the control of lipid metabolism. EMBO Rep. 14, 242–251 (2013).

144. Gierens, H. et al. Interleukin-6 stimulates LDL receptor gene expression via activation of sterol-responsive and Sp1 binding elements. Arterioscler. Thromb. Vasc. Biol. 20, 1777–1783 (2000).

145. Zanoni, P. et al. Posttranscriptional regulation of the human LDL receptor by the U2spliceosome. Circ. Res. 130, 80–95 (2022).

146. Li, H. et al. Identification of mRNA binding proteins that regulate the stability of LDL receptor mRNA through AU-rich elements. J. Lipid Res. 50, 820–831 (2009).

147. Adachi, S. et al. ZFP36L1 and ZFP36L2 control LDLR mRNA stability via the ERK-RSK pathway. Nucleic Acids Res. 42, 10037–10049 (2014).

148. Strøm, T. B., Laerdahl, J. K. & Leren, T. P. Mutations affecting the transmembrane domain of the LDL receptor: impact of charged residues on the membrane insertion. Hum. Mol. Genet. 26, 1634–1642 (2017).

149. Garcia, C. K. et al. Autosomal recessive hypercholesterolemia caused by mutations in a putative LDL receptor adaptor protein. Science 292, 1394–1398 (2001).

150. Wang, J.-Q. et al. SUMOylation of the ubiquitin ligase IDOL decreases LDL receptor levels and is reversed by SENP1. J. Biol. Chem. 296, 100032 (2021).

151. Hahn, J. et al. Genetic loci associated with prevalent and incident myocardial infarction and coronary heart disease in the Cohorts for Heart and Aging Research in Genomic Epidemiology (CHARGE) Consortium. PLoS One 15, e0230035 (2020).

152. Ma, H. et al. LDLRAD3 is a receptor for Venezuelan equine encephalitis virus. Nature 588, 308–314 (2020).

153. Ma, H. et al. The low-density lipoprotein receptor promotes infection of multiple encephalitic alphaviruses. Nat. Commun. 15, 246 (2024).

154. Song, Q. et al. Epidermal SR-A complexes are lipid raft based and promote nucleic acid nanoparticle uptake. J. Invest. Dermatol. 141, 1428–1437.e8 (2021).

155. Srimanee, A., Regberg, J., Hällbrink, M., Vajragupta, O. & Langel, Ü. Role of scavenger receptors in peptide-based delivery of plasmid DNA across a blood-brain barrier model. Int. J. Pharm. 500, 128–135 (2016).

156. Arukuusk, P. et al. Differential endosomal pathways for radically modified peptide vectors. Bioconjug. Chem. 24, 1721–1732 (2013).

157. Ezzat, K. et al. Scavenger receptor-mediated uptake of cell-penetrating peptide nanocomplexes with oligonucleotides. FASEB J. 26, 1172–1180 (2012).

158. Kapinsky, M. et al. Enzymatically degraded LDL preferentially binds to CD14(high) CD16(+) monocytes and induces foam cell formation mediated only in part by the class B scavenger-receptor CD36. Arterioscler. Thromb. Vasc. Biol. 21, 1004–1010 (2001).

159. Miller, Y. I. et al. Minimally modified LDL binds to CD14, induces macrophage spreading via TLR4/MD-2, and inhibits phagocytosis of apoptotic cells. J. Biol. Chem. 278, 1561–1568 (2003).

160. Ly, K., Essalmani, R., Desjardins, R., Seidah, N. G. & Day, R. An unbiased mass spectrometry approach identifies glypican-3 as an interactor of proprotein convertase subtilisin/Kexin type 9 (PCSK9) and low density lipoprotein receptor (LDLR) in hepatocellular carcinoma cells. J. Biol. Chem. 291, 24676–24687 (2016).

161. A. Barrera-Pérez, M., E. Rodríguez-Félix, M., Guzmán-Marín, E., Zavala-Velázquez, J. & Dumonteil., E. Comportamiento biológico de tres cepas de Trypanosoma cruzi de Yucatán, México. Rev Biomed 12, 224–230 (2001).

162. Rajan, A. et al. The human nose organoid respiratory virus model: An ex vivo human challenge model to study respiratory syncytial virus (RSV) and severe acute respiratory syndrome Coronavirus 2 (SARS-CoV-2) pathogenesis and evaluate therapeutics. MBio 13, e0351121 (2021).

163. Adeniyi-Ipadeola, G. O. et al. Infant and adult human intestinal enteroids are morphologically and functionally distinct. MBio 15, e0131624 (2024).

164. Xu, N., Wu, J., Ortiz-Vitali, J. L., Li, Y. & Darabi, R. Directed differentiation of human pluripotent stem cells toward skeletal myogenic progenitors and their purification using surface markers. Cells 10, 2746 (2021).

165. Lian, X. et al. Efficient differentiation of human pluripotent stem cells to endothelial progenitors via small-molecule activation of WNT signaling. Stem Cell Reports 3, 804–816 (2014).

166. Bao, X., Lian, X. & Palecek, S. P. Directed endothelial progenitor differentiation from human pluripotent stem cells via Wnt activation under defined conditions. Methods Mol. Biol. 1481, 183–196 (2016).

167. Liu, X. et al. Differentiation of functional endothelial cells from human induced pluripotent stem cells: A novel, highly efficient and cost effective method. Differentiation 92, 225–236 (2016).

168. Gu, M. Efficient differentiation of human pluripotent stem cells to endothelial cells: Gu. Curr. Protoc. Hum. Genet. 98, e64 (2018).

169. Tohyama, S. et al. Distinct metabolic flow enables large-scale purification of mouse and human pluripotent stem cell-derived cardiomyocytes. Cell Stem Cell 12, 127–137 (2013).

170. Romero-Tejeda, M. et al. A novel transcription factor combination for direct reprogramming to a spontaneously contracting human cardiomyocyte-like state. J. Mol. Cell. Cardiol. 182, 30–43 (2023).

171. Burridge, P. W., Holmström, A. & Wu, J. C. Chemically defined culture and cardiomyocyte differentiation of human pluripotent stem cells: Culture and cardiomyocyte differentiation of human pluripotent stem cells. Curr. Protoc. Hum. Genet. 87, 21.3.1–21.3.15 (2015).

172. Burridge, P. W. et al. Chemically defined generation of human cardiomyocytes. Nat. Methods 11, 855–860 (2014).

173. Deans, R. M. et al. Parallel shRNA and CRISPR-Cas9 screens enable antiviral drug target identification. Nat. Chem. Biol. 12, 361–366 (2016).

174. Tang, G. et al. EMAN2: an extensible image processing suite for electron microscopy. J. Struct. Biol. 157, 38–46 (2007).

175. Chen, M. et al. A complete data processing workflow for cryo-ET and subtomogram averaging. Nat. Methods 16, 1161–1168 (2019).

176. Pettersen, E. F. et al. UCSF ChimeraX: Structure visualization for researchers, educators, and developers. Protein Sci. 30, 70–82 (2021).

177. Islam, M. M., Tamlander, M., Hlushchenko, I., Ripatti, S. & Pfisterer, S. G. Large-scale functional characterization of low-density lipoprotein receptor gene variants improves risk assessment in cardiovascular disease. JACC Basic Transl. Sci. 10, 170–183 (2025).

